# Molecular Networks and Key Regulators of the Dysregulated Neuronal System in Alzheimer’s Disease

**DOI:** 10.1101/788323

**Authors:** Minghui Wang, Aiqun Li, Michiko Sekiya, Noam D. Beckmann, Xiuming Quan, Nadine Schrode, Michael B. Fernando, Alex Yu, Li Zhu, Jiqing Cao, Liwei Lyu, Emrin Horgusluoglu, Qian Wang, Lei Guo, Yuan-shuo Wang, Ryan Neff, Won-min Song, Erming Wang, Qi Shen, Xianxiao Zhou, Chen Ming, Seok-Man Ho, Sezen Vatansever, H. Umit Kaniskan, Jian Jin, Ming-Ming Zhou, Kanae Ando, Lap Ho, Paul A. Slesinger, Zhenyu Yue, Jun Zhu, Sam Gandy, Michelle E. Ehrlich, Dongming Cai, Vahram Haroutunian, Koichi M. Iijima, Eric Schadt, Kristen J. Brennand, Bin Zhang

## Abstract

To study the molecular mechanisms driving the pathogenesis and identify novel therapeutic targets of late onset Alzheimer’s Disease (LOAD), we performed an integrative network analysis of whole-genome DNA and RNA sequencing profiling of four cortical areas, including the parahippocampal gyrus, across 364 donors spanning the full spectrum of LOAD-related cognitive and neuropathological disease severities. Our analyses revealed thousands of molecular changes and uncovered for the first-time multiple neuron specific gene subnetworks most dysregulated in LOAD. *ATP6V1A*, a critical subunit of vacuolar-type H^+^-ATPase (v-ATPase), was predicted to be a key regulator of one neuronal subnetwork and its role in disease-related processes was evaluated through CRISPR-based manipulation of human induced pluripotent stem cell derived neurons and RNAi-based knockdown in transgenic *Drosophila* models. This study advances our understanding of LOAD pathogenesis by providing the global landscape and detailed circuits of complex molecular interactions and regulations in several key brain regions affected by LOAD and the resulting network models provide a blueprint for developing next generation therapeutics against LOAD.

## INTRODUCTION

Sporadic Late Onset Alzheimer’s Disease (LOAD), the most prevalent form of dementia among people over age 65, is a progressive and irreversible brain disorder. Over 5.5 million living in the US are affected by LOAD, which is currently the sixth leading cause of death in the US, and costs more than $200 billion annually ^1^. There is an urgent need to develop effective methods to prevent, treat, or delay the onset or progression of LOAD. The causes of LOAD are poorly understood, with numerous intrinsic and extrinsic factors believed to influence when the disease occurs and how it progresses. Conventional genetic and genome-wide association studies (GWAS) derived primarily from single-nucleotide polymorphism (SNP) analysis have revealed ~30 loci associated with LOAD ^2–5^, and almost 40% of the total phenotypic variance can be explained by these common SNPs ^6^. Translating these genetic findings into biologically meaningful mechanisms of disease pathogenesis and therapeutic interventions remains a huge challenge. A number of studies have performed large-scale molecular profiling (e.g. transcriptomics, proteomics and metabolomics) of postmortem brains from normal control and LOAD patients ^7,8^. Integrating these more functionally relevant datasets holds promise for improving our understanding of the molecular mechanisms of the pathogenesis in sporadic LOAD ^9^.

Leveraging cutting-edge, high-throughput molecular profiling techniques, we recently ^10^ generated a cohort of matched whole-genome sequencing (WGS), whole-exome sequencing (WES), and RNA-sequencing (RNA-seq) data across four brain regions, along with proteomics data generated in one of the regions, from a set of 364 well-characterized brains spanning the full spectrum of LOAD-related cognitive and neuropathological disease severities represented in the Mount Sinai Brain Bank (MSBB) ^8,11^. Systems biology approaches have proven effective for integrating large-scale and diverse biomedical data ^12^, and so this MSBB-AD cohort study offers an unprecedented opportunity to develop more plausible and objective mechanistic models of LOAD. Here we employed a network biology framework to integrate high-dimensional, large-scale DNA, RNA, and clinical data in LOAD to identify predictive molecular signatures and sub-networks underlying early- and late-stage LOAD. The core of our integrative analysis included genetic association, differential expression, correlation, co-expression network ^13^, and causal network ^14,15^ analyses, which together provided an unbiased prediction of novel causal genes and pathways of AD. While the results confirmed some key findings such as elevated immune response from the previous studies of AD^12^, our current study also uncovered a number of new subnetworks and molecular drivers underlying LOAD pathogenesis. This unique collection of multi-Omic LOAD data in tandem with the integrative network biology approaches allowed us to: 1) identify functional pathways dysregulated in LOAD with respect to multiple cognitive/neuropathological outcomes, 2) uncover and prioritize intrinsic co-expressed gene modules across a spectrum of disease stages of LOAD, 3) construct Bayesian probabilistic causal gene regulatory networks by integrating expression quantitative trait loci (eQTLs), transcription factor (TF) and gene expression data, 4) infer the key network hub genes driving key pathways of LOAD, and 5) validate the top ranked novel driver gene by characterizing its functional roles across human induced pluripotent stem cell (hiPSC)-derived neurons and fly models of LOAD (Fig. 1A)..

**Fig. 1.**
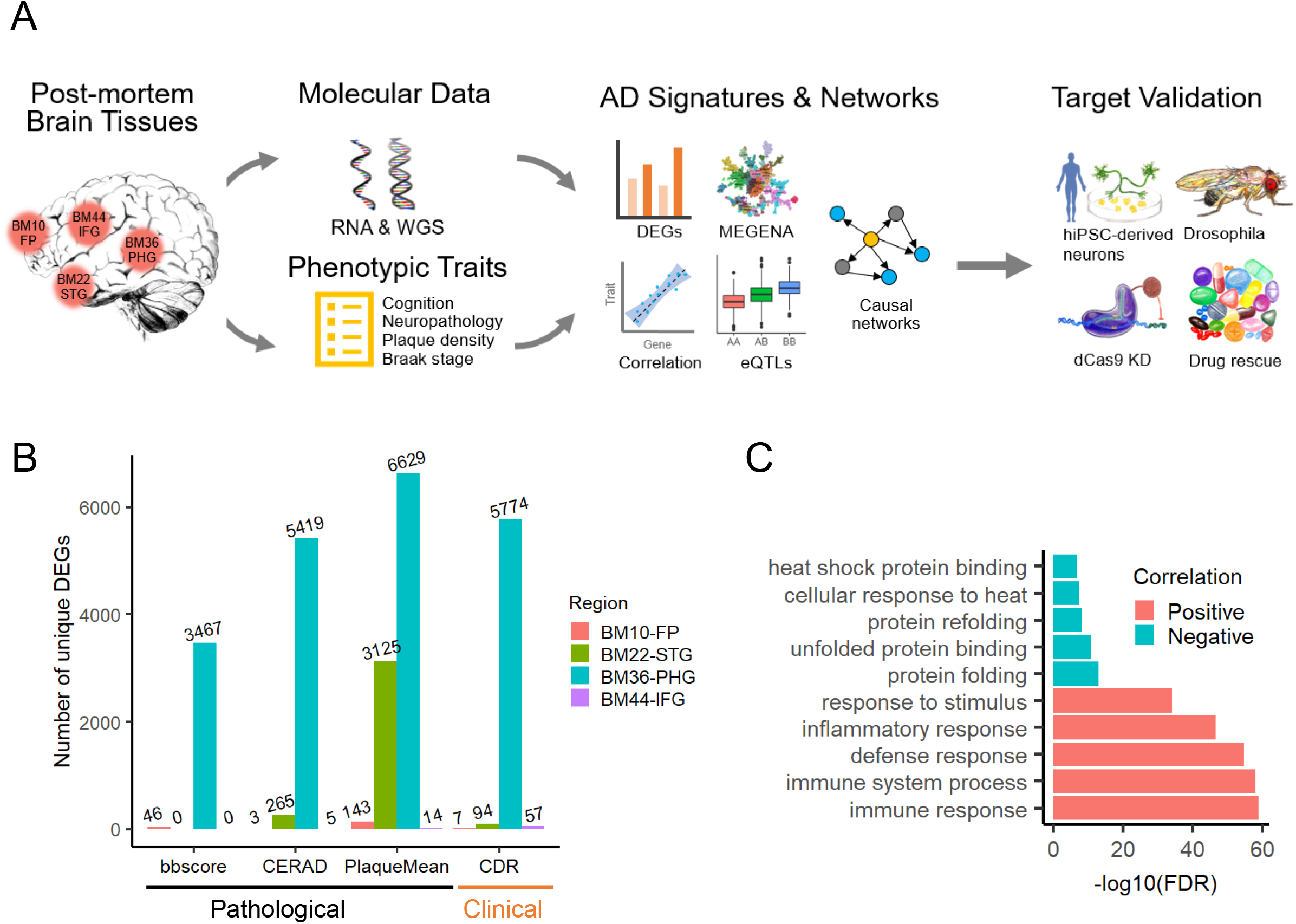
Characterization of MSBB genetics and transcriptomics data. **A**, Schematic of the experimental design and analysis overflow to construct predictive network models and prioritize novel network modules and targets with experiment validation. Postmortem brain tissues from 4 different brain regions in a cohort of 364 brains from the Mount Sinai brain bank (MSBB) with comprehensive AD-related clinical and pathological phenotype characterization are profiled by RNA sequencing (RNA-seq) and whole-genome sequencing (WGS). The molecular data and phenotypic trait data are combined to compute disease signatures and further rank order coexpression network modules that are constructed from the RNA-seq profiles by using MEGENA. The top ranked modules are projected onto Bayesian probabilistic causal networks to identify key driver genes. The disease relevance of the top predicted key driver gene, *ATP6V1A*, is tested in human induced pluripotent stem cell (hiPSC)-derived neurons as well in a Drosophila model of Aβ toxicity through gene perturbations by dCas9 knock-down (KD) or RNA interference (RNAi). Brain region abbreviation: BM10-FP, Brodmann area 10 (BM10) frontal pole; BM22-STG, Brodmann area 22 (BM22) superior temporal gyrus; BM36-PHG, Brodmann area 36 (BM36) parahippocampal gyrus; BM44-IFG, Brodmann area 44 (BM44) inferior frontal gyrus. DEGs, differentially expressed genes. eQTLs, expression quantitative trait loci. MEGENA, multiscale embedded gene co-expression network analysis. **B**, Number of unique DEGs identified in each brain region for each of the clinical/pathological traits. Clinical/pathological traits are bbscore (Braak staging for neurofibrillary tangles), CERAD (neuropathology scale as determined by the Consortium to Establish a Registry for Alzheimer’s Disease protocol), PlaqueMean (mean plaque density), and CDR (clinical dementia rating). **C**, Bar-chart showing the top gene ontology (GO) and pathways enriched in gene pairs with consistent positive or negative correlations across all pairs of brain regions.

## RESULTS

### Study population and molecular data

The MSBB-AD cohort included 364 human brains accessed from the Mount Sinai/JJ Peters VA Medical Center Brain Bank^8,10,11^. The age at the time of death (AOD) of the present population ranged from 61 to 108 years, with a mean and standard deviation (s.d.) of 84.7±9.7. Each donor and corresponding brain sample was assessed for multiple cognitive, medical, and neurological features, including mean plaque density, Braak staging for neurofibrillary tangles (NFT) ^16,17^, clinical dementia rating (CDR) ^18^, and neuropathology scale as determined by the Consortium to Establish a Registry for Alzheimer’s Disease (CERAD) protocol ^19^. These four cognitive/neuropathological traits were scored as semi-quantitative features ranging from normal to severe disease stages, reflecting the continuum and divergence of pathologic and clinical diagnoses of AD beyond a simple case-control classification. Donor brains with no discernable neuropathology or only neuropathologic features characteristic of LOAD were selected. We generated over 1,900 molecular profiles from the MSBB-AD cohort brain specimens, including 1) WGS, 2) WES, and 3) RNA-seq in four brain regions with varying vulnerability to LOAD comprised of Brodmann area 10 (frontal pole, BM10-FP), Brodmann area 22 (superior temporal gyrus, BM22-STG), Brodmann area 36 (parahippocampal gyrus, BM36-PHG) and Brodmann area 44 (inferior frontal gyrus, BM44-IFG) (Fig. 1A). Details about the demographics of the study sample as well as data generation and quality control (QC) have been described previously ^10^. Expanded details regarding the preprocessing of the RNA-seq data, including sample filtering, normalization and covariate correction, are provided in **Supplementary Information (SI)** (**Fig. S1**).

### Identifying gene expression signatures and pathways of LOAD

Differential gene expression analysis was performed to identify genes up-or down-regulated with respect to four LOAD related neuropathological/cognitive traits in each of the four brain regions. The samples were first stratified into multiple disease severity stages for each of the four semi-quantitative traits and then contrasts were made among the severity stages to identify expression changes in each brain region separately (**SI**). The full list of DEGs is provided in **Table S1**. As summarized in Fig. 1B and **Fig. S2**, BM36-PHG had the largest number of differentially expressed genes (DEGs), followed by BM22-STG, BM10-FP and BM44-IFG. This is consistent with our previous pan-cortical transcriptomic analysis of LOAD brains (which were independent of the dataset described herein) in which BM36-PHG was the most impacted region in LOAD at the gene expression level^8^. Our DEG signatures were preserved (adjusted Fisher’s exact test (FET) P value up to 1.0E-100) in ten publicly available AD transcriptomic studies^7,12,20–27^ (**Fig. S3**). Because brain cell-type composition changes over the course of LOAD, we further investigated if the present expression signatures from bulk tissue RNA-seq tend to reflect cell-type changes by overlapping our DEGs with a set of cell type-specific DEGs identified from single-nucleus RNA-seq (snRNA-seq) of LOAD postmortem brains^28^. As shown in **Fig. S4**, we confirmed our down-regulated gene profiles were primarily preserved in down-regulated genes detected in astrocytes, neurons, oligodendrocytes and oligodendrocyte progenitor cell (OPC), while our up-regulated gene profiles were primarily preserved in up-regulated genes in astrocytes and oligodendrocytes (adjusted FET P value up to 6.5E-45), with a few minor exceptions of excitatory neuron or microglia DEGs where there was an enrichment for opposite direction of expression changes. These results demonstrate that we identified a robust set of LOAD-related gene signatures across the four brain regions profiled.

We further analyzed gene ontology (GO) and functional pathways enriched in the DEGs (**Fig. S5** and **Table S2**). As expected, multiple central nervous system (CNS) related gene sets, such as neuronal system, transmission across chemical synapses, and neuroactive ligand receptor interaction, were enriched for down-regulated genes. The KEGG ^29^ Parkinson’s and Alzheimer’ disease gene sets were enriched in the genes down-regulated in demented subjects and in subjects with high plaque density in BM36-PHG. In contrast, matrisome (defined as the ensemble of extracellular matrix (ECM) and related proteins^30^) and associated pathways were most enriched in the up-regulated DEGs in BM36-PHG, followed by immune system, interferon signaling and cytokine signaling pathways. These observations are supported by previous reports that ECM components are associated with the early stages of LOAD ^31^ and memory deficits in APP/PS1 transgenic mice ^32^, perhaps by influencing Aβ fibrillogenesis ^33,34^. Recent genome-wide association studies (GWAS) show enriched AD risk associated with genes expressed by microglia ^2,35–38^, suggesting immunological mechanisms in AD pathogenesis which are consistent with our observation.

We investigated molecular interactions between brain regions by using Pearson’s correlation analysis. Between 176,453 and 1,159,477 (0.03~0.22%) gene pairs showed significant inter-region correlations across the six pairs of brain regions at a conservative Bonferroni corrected P value threshold of 0.05. The correlations for the same genes between any two regions ranged from −0.19 to 0.97 (mean = 0.31 ~ 0.38, median = 0.29 ~ 0.38), indicating the presence of both highly self-correlated genes and inversely-correlated genes. The inverse correlation highlights the importance of multiregional analyses and biological process inferences that can be drawn based on functional connectivity and the complexity of regional interactome. On the other hand, when focusing on the gene pairs from genes with consistent inter-region correlations across all possible region pairs (**Table S3**), immune response pathways were most enriched in the gene pairs with positive inter-region correlations (4.39-fold enrichment (FE), false discovery rate (FDR) = 1.12E-59), while cellular response to unfolded protein (UPR) related pathways were enriched in the gene pairs with negative inter-region correlations (protein folding (8.76-FE, FDR = 1.52-E13), unfolded protein binding (12.93-FE, FDR = 1.57E-11) and protein refolding (29.95-FE, FDR = 9.71E-9)) (**Table S4**, Fig. 1C). Inter-region correlated gene pairs in the immune response and UPR pathways were also correlated within the corresponding individual regions in the same direction, representing coordinated responses to disease-associated triggers within and between the four brain regions.

### Identifying gene modules associated with disease in networks of LOAD brains

To elucidate the coexpression and co-regulation relationships among a large number of gene expression signatures of LOAD, we constructed region-wide gene coexpression networks using MEGENA ^39^, which employs a novel network embedding technique to build up networks (e.g. Fig. 2A) and a multi-scale clustering method to identify coexpressed gene modules (clusters). We identified 475, 527, 441 and 423 coherent gene expression modules in BM10-FP, BM22-STG, BM36-PHG and BM44-IFG, respectively (**Table S5**). Most modules (53.9% to 67.3%) were enriched for MSigDB GO/pathway gene sets (adjusted P value < 0.05) (**Table S6**).

**Fig. 2.**
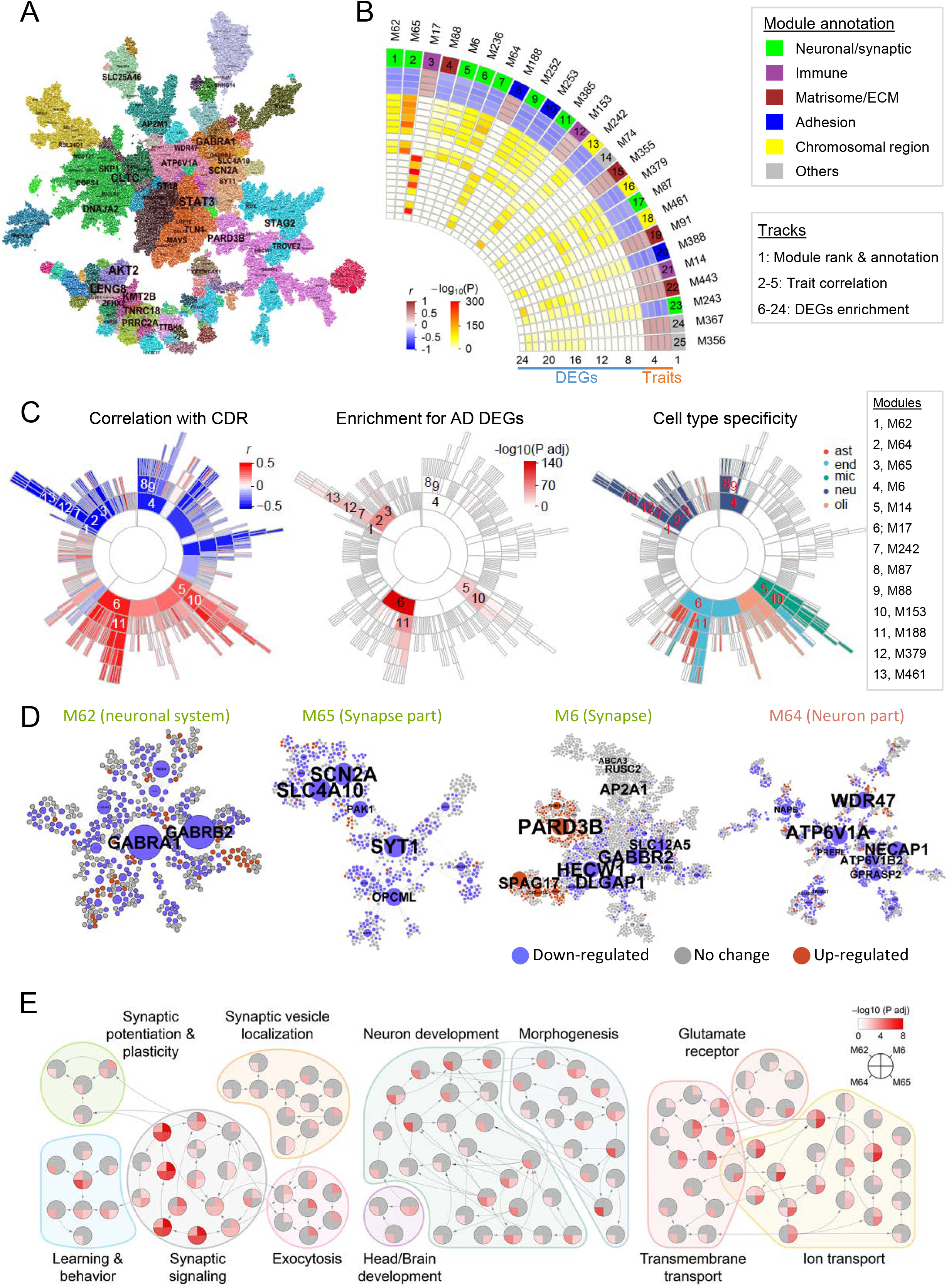
Coexpression network analysis prioritizes neuronal/synaptic modules associated with disease in LOAD brains. **A**, MEGENA network structure in BM36-PHG with nodes colored by module membership. Genes showing high degree of connectivity are highlighted, with font size proportional to the degree of connectivity. **B**, 25 top-ranked MEGENA modules. The heatmap to the left shows the module ranking (number) and GO/pathway annotation (color) in track 1, the module-trait correlation coefficients *r* regarding traits bbscore, CDR, CERAD, and PlaqueMean (tracks 2-5 accordingly), and FDR adjusted P values of enrichment for different sets of DEGs (track 6-24). Specifically, tracks 6-14 denote the enrichment for down-regulated DEGs from AD-vs-normal regarding bbscore, demented-vs-MCI regarding CDR, demented-vs-nondemented regarding CDR, AD-vs-normal regarding CERAD, definite AD-vs-normal regarding CERAD, medium-vs-normal regarding PlaqueMean, severe-vs-medium regarding PlaqueMean, severe-vs-mild regarding PlaqueMean, and severe-vs-normal regarding PlaqueMean, accordingly. Tracks 15-23 denote the enrichment for up-regulated DEGs from the same contrasts as the down-regulated DEGs. An additional track 24 denotes enrichment for up-regulated DEGs from mild-vs-normal regarding PlaqueMean. **C**, Sunburst plots showing the module hierarchy and correlation with CDR (left), and enrichment for CDR demented-vs-nondemented DEGs (middle) and enrichment for cell type-specific markers (right). Numbers 1-13 denote 13 top ranked modules as listed to the right. Cell types are denoted by abbreviations: ast, astrocytes; end, endothelial; mic, microglia; neu, neurons; oli, oligodendrocytes. **D**, Representative visualization of the top ranked neuronal/synaptic modules M62, M65, M6 and M64. Node color denotes whether the gene is up-regulated (red), down-regulated (blue), or no change (grey) in demented brains. Node size is proportional to the network connectivity within each module. **E**, Top ranked neuronal/synaptic modules enriched in GO biological process (BP) hierarchy in relation to synaptic function, neuronal development and transportation. Each node denotes a GO/BP term, with a pie-chart displaying the –log_10_(adjusted P value) of the Fisher’s exact test (FET) enrichment for the 4 top ranked neuronal/synaptic modules (i.e. M6, M62, M64 and M65). The terms are grouped into general categories as indicated by different background color blocks. Arrows denote the direction from a parent term to a child term. The GO hierarchy was extracted from the R/Bioconductor package GO.db and the GO/BP annotation gene sets were obtained from the R/Bioconductor package org.Hs.eg.db.

To prioritize the gene modules with respect to their association to LOAD pathology, we applied an ensemble ranking metric^8^ across multiple feature types (Fig. 2B-C), including 1) correlations between module eigengenes (i.e. the first principal component of module gene expression profile) and cognitive/pathological traits associated with LOAD, and 2) enrichment for the DEG signatures identified above. A more complete description of the information used to rank the modules is included in **Table S7**. The ranking of the top 25 MEGENA modules are illustrated in Fig. 2B, with all of the top modules coming from the BM36-PHG region.

Distinct from our previous transcriptomic network analysis in an independent cohort and brain regions which prioritizes a diverse set of LOAD-related subnetworks including immune response, glutathione transferase, and cell junction, etc. ^12^, the present network analysis highlights the significance of multiple neuronal/synaptic modules in driving the disease process in the MSBB-AD cohort. Nine of the top 25 modules were associated with synaptic transmission/neuronal systems, negatively correlated with disease traits, and enriched for DEG signatures down-regulated in LOAD, including M62 (ranked 1^st^, denoted as #1 for simplicity) and M65 (#2), M6 (#5), M236 (#6), M64 (#7), M252 (#9), M385 (#11), M87 (#17) and M243 (#23). These nine modules, along with three other chromosomal region related modules M242 (#13), M379 (#16) and M461 (#18), showed neuron specific expression^40^ (Fig. 2C). M64 module was overrepresented with inhibitory neuron-enriched genes, while six others (M6, M87, M65, M236, M62, and M252) were overrepresented with excitatory neuron-enriched genes ^41^ (**Table S8**), suggesting different neuron subtypes might be involved in these modules. The topological structures for four of the top ranked neuronal/synaptic modules are shown in Fig. 2D. A number of LOAD genetic candidate risk genes were present in these top ranked modules, such as *MEF2C* in the first-ranked module M62, *CELF1*, *MADD*, *PLD3*, *PTK2B*, and *ZCWPW1* in the fifth-ranked module M6, and *APP* and *SORL1* in the seventh-ranked module M64.

We further investigated whether the top ranked neuronal/synaptic modules were involved in distinct BPs by overlapping the genes of the 4 top ranked parental neuronal/synaptic modules (i.e. M6, M62, M64 and M65) onto the GO/BP hierarchy (Fig. 2E and **S6**). All these 4 modules are enriched in synaptic signaling to different degrees, but M6 and M64 are also enriched in regulation of long-term synaptic potentiation, synaptic vesicle trafficking and localization. M64 is also enriched in learning and memory related pathways. M6 is highly enriched in exocytosis such as regulation of synaptic vesicles exocytosis. M64 and M65 are enriched in CNS development and morphogenesis, but M64 is also enriched in neurogenesis, neuron development related pathways. While all 4 neuronal/synaptic modules are enriched in transporter activities, they are likely to participate in different aspects of transporter activities. For example, M64 is significantly enriched in ATP hydrolysis coupled proton transport. On the other hand, the other modules are enriched in glutamate receptor signaling, which is consistent with the above neuron subtype enrichment analysis. Taken all together, M64 is the most distinct module in terms of potential involvement in biological signaling pathways.

We validated the biological coherence (“preservation”) of the top ranked neuronal/synaptic modules in two previously published independent LOAD-related postmortem brain cohort studies, including 1) the Harvard Brain Tissue Resource Center (HBTRC) microarray data ^12^ and 2) the ROSMAP RNA-seq data ^7^. Among 111 modules in the coexpression network from the HBTRC dataset, a synaptic transmission enriched module (purple) was ranked the 16^th^ for relevance to LOAD pathology and this module was enriched in all the nine top ranked neuronal/synaptic modules in the current study (FE ranging from 2.7 to 8.7, FDR up to 1.2E-62). Of four neuronal/synaptic modules (m16, m21, m22 and m23) in the coexpression network from the ROSMAP dataset, m21 and m23 were significantly overlapped with all the current nine top ranked neuronal/synaptic modules, while m16 and m22 were enriched in three and seven of the current top ranked neuronal/synaptic modules, respectively (FE ranging from 1.4 to 14.1, FDR up to 3.5E-39). It is noted that the m23 module showed a positive correlation with cognitive decline (P value = 9.1E-4), and the m22 and m23 modules showed a negative correlation with amyloid β burden (P value = 3.6E-3 and 3.8E-3, respectively), in the ROSMAP cohort.

Three of the present top modules, M17 (#3), M153 (#12) and M14 (#21), were enriched with immune response pathways. All were positively correlated with disease traits and enriched for up-regulated DEGs in AD. While M14 (#21) and its sub-module M153 (#12) were enriched for microglia markers, M17 (#3) was enriched for endothelial markers (Fig. 2C). Two LOAD genetic candidate risk genes, *CLU* and *CR1*, were members of M17 while seven LOAD genetic candidate risk genes (*APOE*, *CASS4*, *CD33*, *HLA-DRB1*/*HLA-DRB5*, *INPP5D*, *MS4A4A*/*MS4A6A* and *TREM2*) fell into M153 and M14, resulting in a 15.4-FE for AD risk genes in M153 (FET P = 3.8E-9) and an 11.6-FE in M14 (FET P = 4.2E-8). M153 included five GWAS genes (*CD33*, *HLA-DRB1*, *INPP5D*, *MS4A6A* and *TREM2*) that were up-regulated in LOAD (**Fig. S7**). The clustering of the four up-regulated risk genes (*CD33*, *INPP5D*, *MS4A6A*, and *TREM2*) in the same module was previously observed in an independent coexpression network analysis from the ROSMAP cohort ^42^, indicating a potential core component of LOAD-related immune response pathway conserved in multiple datasets.

Besides neuronal system and immune response enriched modules known to be associated with LOAD, several less studied pathways emerged in the top ranked modules, including matrisome enriched modules M88 (#3) and M91 (#19), ECM enriched modules M355 (#15) and M443 (#22), cell proliferation enriched module M367 (#24), and alpha hemoglobin stabilizing protein (AHSP) pathway enriched module M356 (#25). These results indicate important potential pathways of LOAD that warrant further investigation.

### Bayesian network analysis predicts novel key drivers of synaptic transmission/neuronal system pathways implicated in LOAD

Coexpression network analysis revealed the global gene coregulatory landscape and identified synaptic transmission and neuronal system enriched modules as highly associated with LOAD pathology. To determine potential casual relationships between coexpressed genes in these top ranked modules for predicting key regulators of the corresponding pathways underlying disease initiation and progression, we constructed Bayesian probabilistic causal networks (BNs) by integrating genetics (e.g. WGS SNP variants), gene expression, and known TF-target relationships. We first mapped eQTLs by integrating the RNA-seq and WGS-based SNP genotype data. We identified more than 92,336 SNP-gene pairs that were associated locally (so-called *cis*-SNP-gene pair) in any brain region, with more than 66.1% shared in at least another region. Details about the identification and replication of brain region-wide eQTLs are provided in **SI** (see also **Table S9-13** and **Fig. S8-12**). Notably the current eQTL analysis indicated marked common genetic regulation occurring across different brain regions. We applied causal inference test (CIT) ^43,44^ to use eQTLs as instrumental variables to compute the probability of the possible causal regulatory chain among genes associated with the same eQTL; these causal chain probability results were incorporated as structure priors into the BN inference procedure. Fig. 3A shows the BN for BM36-PHG.

**Fig. 3.**
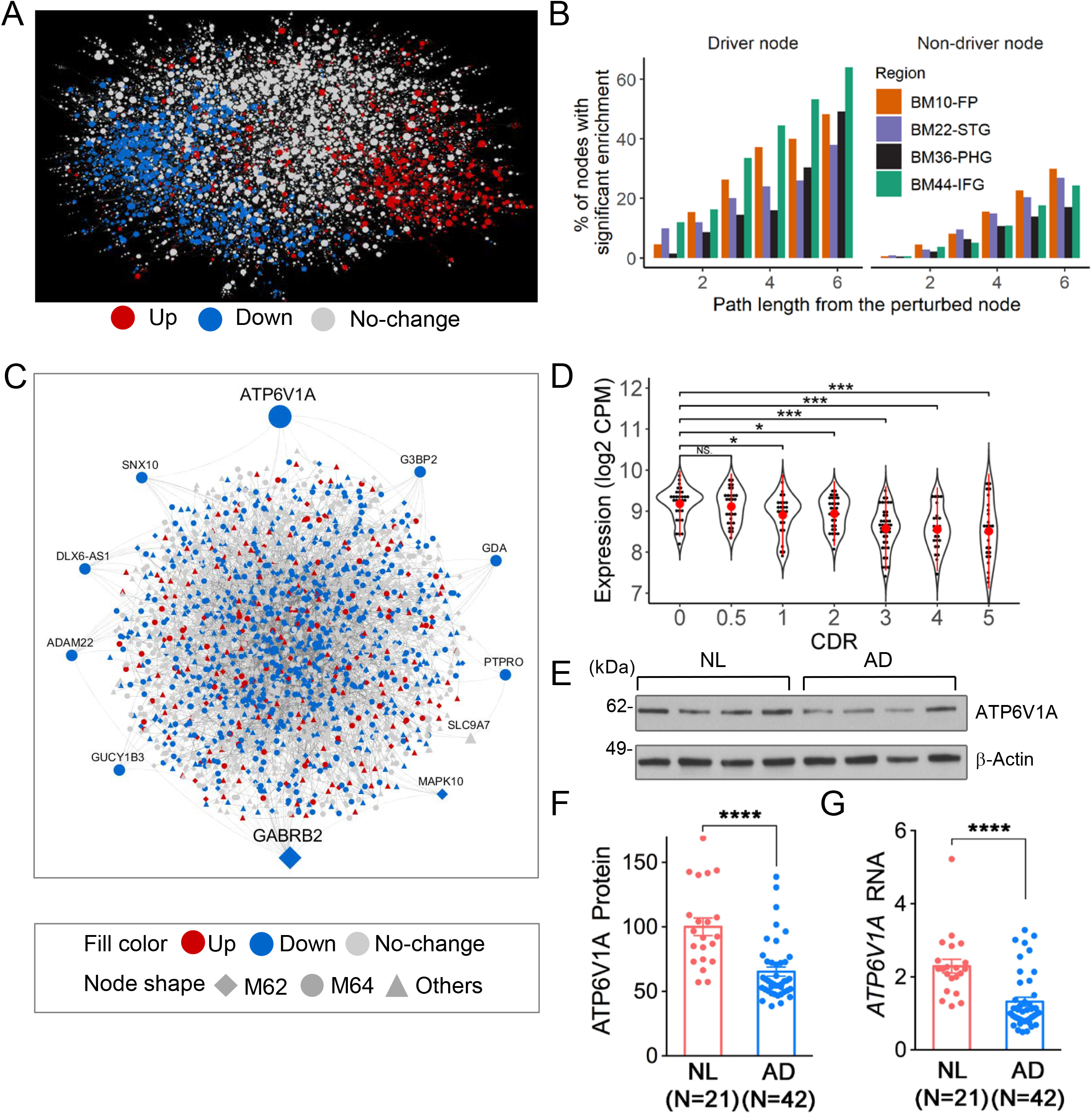
Bayesian probabilistic causal network (BN) analysis predicts novel key drivers of LOAD. **A**, Global network topology of the BN in BM36-PHG. Node color denotes whether the gene is up-regulated (red), down-regulated (blue), or no change (grey) in demented brains. **B**, Summary of validation of BN structure using gene perturbation signatures. The left panel shows percentage of the global BN key driver nodes (genes) whose network neighborhood was enriched for perturbation signature at different path length from the node. The right panel shows the same analysis results for non-driver nodes. **C**, Projection of neuronal modules M62 and M64 on to the BM36-PHG BN. Node labels are shown for the module key drivers, such as the top ranked driver *ATP6V1A*. Node shape denotes the module membership. Node color denotes expression change in demented brains. **D-G**, A novel network key driver *ATP6V1A* is down-regulated in LOAD patient. **D**, Violin plot shows the *ATP6V1A* expression in the RNA-seq analysis of BM36-PHG brain region as stratified by CDR. **E-G** illustrate the validation of *ATP6V1A* expression change in MSBB BM36-PHG brain samples using western blot (**E-F**) and qRT-PCR (**G**) analyses. **E**, Representative western blot of ATP6V1A protein level. (Student’s t-test. *p < 0.05. **p < 0.01. ***p < 0.001. ****p < 0.0001. NS, no significance.). NL, normal control.

Due to limited availability of replication data, validating the BN structure is not trivial and it has been reported that replication of edge-to-edge is strongly dependent on the sample size ^45^. In contrast, highly connected key driver nodes tend to be more stable than network edges ^45^. Indeed, a significant number of global key drivers ^46^ were shared between any two BNs (**Fig. S13**). In light of these findings, we sought to examine whether publicly available experimental gene perturbation signatures of the key drivers were predicted by our networks, by testing whether the genes in these signatures were enriched for genes in the network neighborhood of the key driver in our BNs. As illustrated in Fig. 3B, we demonstrated that 50~60% of the key driver perturbation signatures were enriched in the network neighborhoods of the corresponding key drivers across the four region-wide BNs, while the proportion of significantly enriched perturbation signatures decreased to 20~30% in the network neighborhood of non-driver genes.

We projected each of the nine top ranked neuronal/synaptic modules (M62, M65, M6, M236, M64, M252, M385, M87, and M243) (Fig. 2B) onto the BM36-PHG BN and identified network key drivers that might potentially modulate a large number of nodes in these modules using the key driver analysis ^46^. We identified 48 key drivers (42 unique genes) across nine modules (**Table S14**), including ten key drivers that were root nodes in the BN without parental nodes (**Table S15**). To further verify the root node status beyond a single region-based network, we integrated information from all four brain region-wide BNs by building a union BN, combining directed links from four region-wide BNs following previous practices ^47,48^. Two key drivers, *ATP6V1A* in M64 and *GABRB2* in M62, remained as root nodes in this union BN. We considered these two key drivers as the most likely process initiators of the neuronal/synaptic modules underlying the LOAD pathogenesis in the current dataset. Fig. 3C shows the projection of M62 and M64 onto the BM36-PHG BN, while **Fig. S14** illustrates more detailed network structures surrounding *ATP6V1A* and *GABRB2* on the BM36-PHG BN, respectively.

While *GABRB2* exhibits a region-dependent up- or down-regulation in LOAD based on prior studies (**SI**), the other prioritized key driver, *ATP6V1A*, was more consistently down-regulated across brain regions and disease stages in LOAD. This gene encodes a component (V1 subunit A, V1a) of vacuolar ATPase (v-ATPase), a multi-subunit enzyme that mediates acidification of eukaryotic intracellular organelles. v-ATPase is a component of the mTOR pathway and functions as a lysosome-associated machinery for amino acid sensing ^49,50^. A potential role of *ATP6V1A* in neuronal development was suggested by studying *de novo* mutations in this gene ^51^. Yet, there are no prior studies linking changes in *APT6V1A* expression with LOAD pathogenesis. *ATP6V1A* was significantly down-regulated in the BM36-PHG (−1.43 fold, P value = 1.5E-6) and BM22-STG (−1.25 fold, P value = 2.1E-3) regions of persons with dementia (CDR ≥ 1), and it was marginally down-regulated in the BM10-FP region of persons with MCI and frank dementia (CDR = 0.5) (−1.11 fold, P value < 0.098) (Fig. 3D and **Fig. S15**). In addition, *ATP6V1A* expression was negatively correlated with cognitive/neuropathological traits in BM22-STG and BM36-PHG (Spearman correlation coefficients ranges between −0.21 and −0.44 and the corresponding P values are between 5.9E-11 to 3.3E-4), suggesting down-regulation of *ATP6V1A* was a consistent event at both early and late stages of the disease. We validated the reduced expression of *ATP6V1A* in LOAD brains by quantitative reverse transcription polymerase chain reaction (qRT-PCR) and western blot analyses using BM36 brain samples (Fig. 3E-G; 42% decrease at mRNA level and 35% decrease at protein level, P < 1.0E-4). Down-regulation of *ATP6V1A* was also previously identified in cortical neurons of the superior frontal gyrus ^21^ and the hippocampus CA1 area ^26^ of LOAD brains. In addition, it was also found to be down-regulated in the excitatory (0.8 fold, adjusted P value 2.6E-117) and inhibitory (0.83 fold, adjusted P value 6.7E-22) neurons in brains with early-pathology of LOAD compared to no-pathology brains in the ROSMAP cohort ^28^. Thus, multiple lines of evidence indicate that *ATP6V1A* is a novel therapeutic target for LOAD. To validate the functional role of *ATP6V1A* in LOAD, we performed gene perturbation experiments in both *in vitro* (hiPSC-derived neurons) and *in vivo* (transgenic flies) models.

### Decreased neuronal activity in *ATP6V1A*-deficient *NGN2*-neurons

As *ATP6V1A* was both down-regulated in the LOAD brains and enriched for neuronal expression, we developed a model of hiPSC-derived neurons with reduced expression of *ATP6V1A*. To repress endogenous *ATP6V1A* expression, we utilized CRISPR inhibition (CRISPRi) ^52^, in which dCas9 (dead Cas9) is fused to the transcriptional repressor of Krüppel associated box (KRAB) ^53^. We designed six sgRNAs (crispr-era.stanford.edu) to target the promoter region for knockdown (KD) of the *ATP6V1A* gene (Fig. 4A; see **SI** for sgRNA sequences), and individually cloned each into a lentiviral vector (lentiGuide-Hygro-mTagBFP2)^52^. By qRT-PCR and western blot, we identified two sgRNAs (*ATP6V1A*-i1 and i2) that efficiently repressed *ATP6V1A* expression in NPCs from two donors stably expressing dCas9^−KRAB^ (**Fig. S16A-C**). Following *NGN2*-induction (**Fig. S16D**)^54^, both *ATP6V1A* RNA (60~70% transcriptional repression, P < 0.001, Fig. 4B) and protein levels (80~90% translational repression, P < 0.001, Fig. 4C-D) were significantly reduced in 21-day (D21) cultured neurons.

**Fig. 4.**
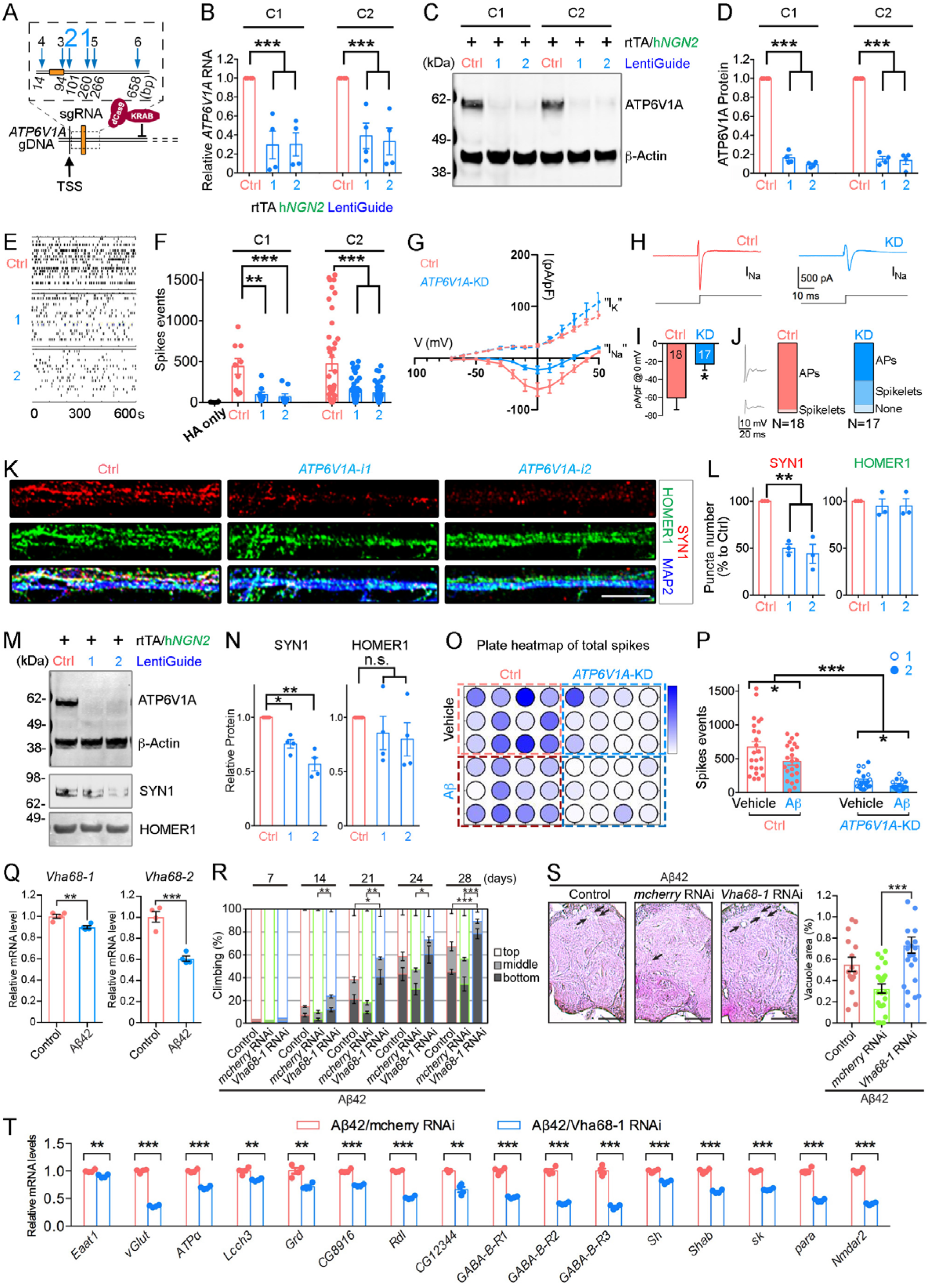
Repression of *ATP6V1A* leads to neuronal malfunction in human *NGN2*-neurons and Aβ42 transgenic flies. **A**, Schematic diagram of the *ATP6V1A* gene editing by CRISPR/dCas9-KRAB system. Six different sgRNAs are designed for targeting the *ATP6V1A* promoter. **B**, qRT-PCR analysis (n = 4 biological replicates) confirms the decreased *ATP6V1A* RNA by sgRNA candidate 1 & 2 (i1 and i2) in two independent *NGN2*-neuron cell lines (i.e. C1 and C2). **C**-**D**, Representative western blot and quantitative analysis (n = 4) of ATP6V1A protein level in *NGN2*-neurons. β-Actin is a loading control. **E**-**F**, Representative raster plots of spike events over 10 min and analysis (n = 6~45 wells) of day 21 (D21) *NGN2*-neurons. **G**, Current-voltage (I-V) plot for inward sodium (I_Na_) and outward potassium (I_K_) currents. Current density (pA/pF) is shown. Holding potential was −80 mV. **H**, Representative examples of putative inward voltage-gated sodium current at 0 mV. **I**, Bar plot shows mean inward sodium current densities at 0 mV for *ATP6V1A* KD neurons (n=17) and control neurons (n=18), (*p = 0.0151; unpaired t-test). **J**, Box plots show fraction of neurons that displayed a full action potential (AP), spikelets, or no events with a current injection step (0.1 nA) positive to threshold for control and KD neurons. Inset shows representative examples of AP & spikelet. **K**, Representative confocal microscopic images of synapse plasticity-related proteins (SYN1, red; HOMER1, green) and pan-neuronal marker MAP2 (blue). Bar, 20 µm. **L**, Analysis of SYN1 and HOMER1-immunoreactive puncta numbers (n = 3). **M-N**, Representative western blot and quantitative analysis (n = 4) of SYN1 and HOMER1 levels. **O-P**, Multi-electrode array after exposure to 5 µM beta-amyloid at 24 hours. **O**, plate map of total spike events; **P**, analysis of spike events (n = 12 wells). (*t*-test and ANOVA; *p < 0.05; **p < 0.01; ***p < 0.001, n.s., no significance; Error bars represent SE.) **Q**, mRNA expression levels of *Vha68-1*and *Vha68-2* were decreased in the heads of flies carrying Aβ42 transgene. mRNA levels were analyzed by qRT-PCR. n = 4, **p < 0.01 and ***p < 0.001. **R**, Knockdown of *Vha68-1* in neurons exacerbated locomotor deficits caused by Aβ42 as revealed by climbing assay. Average percentages of flies that climbed to the top (white), climbed to the middle (light gray), or stayed at the bottom (dark gray) of the vials. Percentages of flies that stayed at the bottom were subjected to statistical analyses. n = 5 independent experiments except for 7 days (n =2), *p < 0.05, **p < 0.01 and ***p < 0.001 by Student’s t-test. **S**, Neuronal knockdown of *Vha68-1*significantly worsened neurodegeneration in the neuropil region in Aβ42 fly brains. Representative images show the central neuropil in paraffin-embedded brain section with hematoxylin and eosin (HE) staining from 33-day-old flies. Scale bars: 50 μm. Percentages of vacuole areas (indicated by arrows in the images) were subjected to statistical analyses. n = 12-24 hemispheres, ***p < 0.001 by Student’s t-test. The genotypes of the flies were: (Control): *elav*-GAL4/Y, (Aβ): *elav*-GAL4/Y; UAS-Aβ42/+, (Aβ/*mcherry*RNAi): *elav*-GAL4/Y; UAS-Aβ42/+; UAS-*mcherry*RNAi/+, and (Aβ/*Vha68-1* RNAi): *elav*-GAL4/Y; UAS-Aβ42/+; UAS-*Vha68-1*RNAi/+. **T**, mRNA expression levels of 16 selected genes related to synaptic activities were significantly reduced in Aβ42-expressing files with neuronal KD of *ATP6V1A*/*Vha68-1*. mRNA levels were analyzed by qRT-PCR (n = 4).

*ATP6V1A* plays a unique role in synapse function ^55^. Therefore, we determined whether *ATP6V1A* repression influenced spontaneous neuronal electric activity. Isogenic pairs of control and *ATP6V1A* CRISPRi *NGN2*-neurons (co-cultured with human fetal astrocytes to enhance synaptic maturation) were evaluated across a panel of assays to explore their impact on synaptic function and synapse plasticity-associated proteins. We applied an Axion multi-electrode array (MEA) to assess the impact of *ATP6V1A* repression on population-wide neuronal activity, including frequency and coordination of network firing. Significantly reduced neuronal activity was observed following perturbations with either sgRNA (average 4.32-fold down in D21 *NGN2*-neurons, P < 0.01; Fig. 4E-F). We further measured the amplitude of voltage-gated potassium (I_K_) and sodium current (I_Na_) using whole-cell patch-clamp recordings (Fig. 4G-I). *ATP6V1A* KD neurons exhibited significantly smaller I_Na_ current density (P = 0.015), but no significant change in I_K_ current (Fig. 4G-I). Consistent with a decrease of I_Na_, RNA sequencing of *NGN2*-neurons at 21 days (detailed below) revealed significantly reduced mRNA expression of different voltage gated sodium channel subunits, such as *SCN3A*, *SCN2A*, and *SCN4B* (**Fig. S17**). Lastly, we observed a decrease in the number of full action potentials and increase in immature spikes (e.g. spikelets) in the *ATP6V1A* CRISPRi group (Fig. 4J).

To explore the effect of *ATP6V1A* on synaptic components, *NGN2*-neurons were immunostained against the presynaptic marker SYN1 and the postsynaptic marker HOMER1 and analyzed by confocal imaging (Fig. 4K). A significant reduction in SYN1^+^ puncta number following *ATP6V1A* CRISPRi was observed (1.1-fold down, P < 0.001; Fig. 4K-L) across two sgRNAs, whereas CRISPRi had had limited effect on HOMER1 (Fig. 4L). Western blot assay showed similar results. A 25-45% reduction of SYN1 (P < 0.05) was observed, while HOMER1 was expressed at comparable levels regardless of CRISPRi (Fig. 4M-N). qRT-PCR indicated that in *ATP6V1A*-deficient *NGN2*-neurons, only presynaptic components (SYN1, vGLUT1) were significantly decreased in RNA (~20% down, P < 0.05 and ~38% down, P < 0.01, respectively **Fig. S18A**). While postsynaptic components (HOMER1 and PSD95) showed no significant change, vGLUT1 protein level decreased by approximately 22% (P < 0.05, **Fig. S18A-C**).

AD neuronal pathology is associated with extracellular beta-amyloid (Aβ) aggregates ^56^. Aβ administration (24 hours, 5 µM) significantly decreased spontaneous neuronal activity (P < 0.05), without altering *ATP6V1A* gene expression (Fig. 4O-P and **Fig. S19A-C**). Moreover, *ATP6V1A* repression in combination with Aβ42 exposure further impaired neuronal activity (p < 0.05, Fig. 4O-P and **Fig. S19B-C**).

### Neuronal knockdown of *Vha68-1*, a fly ortholog of *ATP6V1A*, worsens behavioral deficits and neurodegeneration in Aβ42 flies

In addition to the *in vitro* validation of neuronal activity impairment by *ATP6V1A* deficit in hiPSC-derived neurons, we also evaluated whether there were behavior and cellular consequences caused by knocking down ortholog of *ATP6V1A* using transgenic *Drosophila* models. According to the DIOPT (DRSC Integrative Ortholog Prediction Tool), *Drosophila* Vacuolar H^+^ ATPase 68kD subunit 1 (*Vha68-1*, *CG12403*) and *Vha68-2* (*CG3762*) are the best orthologs of human *ATP6V1A* proteins.

We first examined the effects of neuronal KD of *Vha68-1* and *Vha68-2* on neuronal integrity during aging in flies. The pan-neuronal *elav*-GAL4 driver was used to express shRNAi targeting *Vha68-1* or *Vha68-2* in neurons. Since both *Vha68-1* and *Vha68-2* are essential genes for viability in flies and RNAi-mediated KD of *Vha68-2* in neurons resulted in lethality of flies, we selected an RNAi line that could modestly reduce mRNA expression levels of *Vha68-1* **(Fig. S20A**). We found that neuronal KD of *Vha68-1* by itself caused modest decline in climbing ability in aged flies (**Fig. S20B**).

We next examined whether neuronal KD of *Vha68-1* worsened neuronal dysfunction and neurodegeneration in a transgenic *Drosophila* model that expresses human Aβ42 in the brain^57^. qRT-PCR analysis detected significant reduction in mRNA expression levels of both *Vha68-1* and *Vha68-2* in Aβ42 fly brains (Fig. 4Q), suggesting that downregulation of fly orthologs of human *ATP6V1A* may play a role in Aβ42-mediated toxicity. The pan-neuronal *elav*-GAL4 driver was used to express both Aβ42 and shRNAi targeting *Vha68-1* or control shRNAi in neurons. The Aβ42 flies showed age-dependent locomotor deficits assessed by the forced climbing assay ^57^, which was significantly exacerbated by neuronal KD of *Vha68-1* compared to control flies (Fig. 4R). To minimize potential off-target effects of RNAi, the climbing assay was repeated in an independent transgenic line carrying shRNAi targeting a different region of *Vha68-1* (**Fig. S20C-E**). Although the KD efficiency of *Vha68-1* by this shRNAi was weaker (**Fig. S20C**) and *Vha68-1* KD by itself did not cause climbing defects (**Fig. S20D**), locomotor deficits in Aβ42 flies were slightly enhanced (**Fig. S20E**).

In *Drosophila*, brain vacuolation is a morphological hallmark of neurodegeneration and neuronal expression of Aβ42 causes age-dependent appearance of vacuoles in the central neuropils and cell bodies in the fly brains ^57^. We found that KD of *Vha68-1* significantly worsened neurodegeneration in the neuropil region in Aβ42 fly brains (Fig. 4S).

To further ask whether altered neuronal activity underlies toxic interactions between *ATP6V1A*/*Vha68-1* deficiency and Aβ42 in flies, we examined mRNA expression levels of 16 genes related to synaptic activities, focusing on GABAergic and glutamatergic systems as well as ion channels (Fig. 4T and **S20F-G**). We found that mRNA expression levels of 9 out of 16 genes were significantly reduced by neuronal KD of *ATP6V1A*/*Vha68-1* in fly brains (**Fig. S20F**), while 5 out of 16 genes were significantly reduced in Aβ42 fly brains (**Fig. S20G**). Three genes, including *SLC1A2/Eaat1*, *SLC17A6-8/vGlut* and *ATP1A1-3/ATPα*, were commonly reduced in both conditions (**Fig. S20F-G**). By contrast, neuronal KD of *ATP6V1A*/*Vha68-1* in Aβ42 flies dramatically reduced mRNA expression levels of all 16 genes compared to control (Fig. 4T). Of particular interest, three key driver genes including *GABRA1*/*Grd* in M62, *SCN2A*/*para* in M65 and *GABBR2*/(*GABA-B-R2, 3*) in M6 were downregulated in these fly brains, suggesting functional links between these networks and *ATP6V1A*/*Vha68-1* in M64 module.

In summary, these results suggest that *Vha68-1/ATP6V1A* deficiency and Aβ42 synergistically downregulate key regulator genes of neuronal activity and exaggerate Aβ42-induced toxicities in flies.

### *ATP6V1A* KD signatures are enriched in *ATP6V1A* regulated networks in human LOAD brains

To characterize the molecular changes and validate the sub-network regulated by *ATP6V1A*, we performed RNA-seq analysis on RNA from four groups of hiPSC-derived *NGN2*-neurons (designated WT-V and WT-Aβ for vehicle-treated and Aβ-treated *ATP6V1A* wild-type (WT) neurons, respectively, and KD-V and KD-Aβ for vehicle-treated and Aβ-treated *ATP6V1A* KD neurons, respectively), with five independent neuronal differentiation experiments in each group (**SI**). At a cutoff of FDR adjusted P value ≤ 0.05 and FC ≥ 1.2, no significant expression change was observed between Aβ-treated cells and vehicle-treated cells in either *ATP6V1A* KD or WT genotype. In contrast, three genes (1 up-regulated and 2 down-regulated) showed significant expression change in *ATP6V1A* deficit neurons in vehicle-treated condition (KD-V vs WT-V), while 55 DEGs (18 up-regulated and 37 down-regulated) were detected in *ATP6V1A* deficit neurons under Aβ-treated condition (KD-Aβ vs WT-Aβ) (**Table S16 and Fig. S21**). Furthermore, the contrast between KD-Aβ and WT-V presented 326 DEGs (110 up-regulated and 216 down-regulated ones) (**Table S16**), the largest number among all the comparisons.

We examined the GO/pathways impacted by *ATP6V1A* deficit and/or Aβ treatment by employing the Gene Set Enrichment Analysis (GSEA) ^58^. Fig. 5A illustrates the top GO/pathways enriched in the molecular profiles; the full list of significantly enriched MSigDB GO/pathways is provided in **Table S17**. A number of GO/pathways were commonly impacted by KD (KD-V vs WT-V) and Aβ treatment (WT-Aβ vs WT-V), such as up-regulation of ribosome, translation, and DNA packaging, and down-regulation of axoneme assembly and cilium movement. As expected, we found V-ATPase transport and phagosome maturation/acidification were down-regulated in KD-V vs WT-V. Consistent with the functional assay above, we noted that KD-V vs WT-V led to down-regulation of multiple synaptic and presynaptic pathways, with the down-regulation even more significant after the exposure to Aβ treatment in KD-Aβ vs WT-Aβ. In addition, it is noted that UPR and ER stress response were up-regulated in conditions with Aβ treatment.

**Fig. 5.**
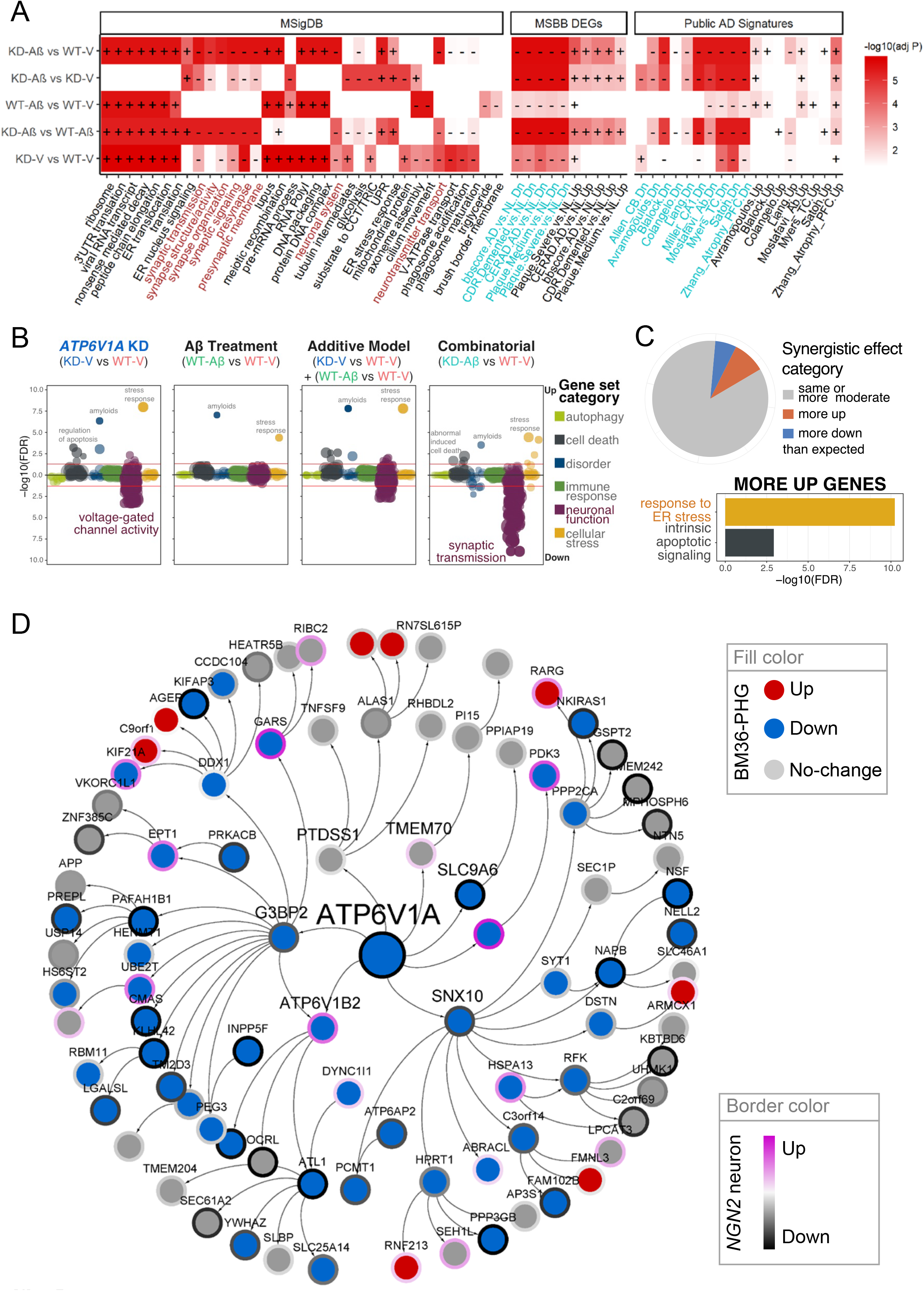
RNA-seq analysis reveals that *ATP6V1A* KD impacts neuronal function related pathways and *ATP6V1A* KD signatures are enriched in *ATP6V1A* regulated networks in human LOAD brains. **A**, Summary of top MSigDB GO/pathway gene sets and human AD signatures, including the present MSBB BM36-PHG DEGs and public AD DEGs, which were significantly enriched in molecular profiles from the perturbations of *NGN2*-neurons. Plus (+) and minus (−) symbols denote the sign of enrichment scores as computed by gene set enrichment analysis (GSEA). Brown color in the x-axis of the left panel highlights the neuronal and synaptic related terms. Cyan color in the x-axis of the two panels to the right highlights the down-regulated signatures. **B-C**, Analysis of synergistic effect between *ATP6V1A* KD and Aβ treatment in *NGN2*-neurons. **B**, Summary of the functional categories that are likely to be impacted by the synergistic effect between *ATP6V1A* KD and Aβ treatment in *NGN2*-neurons. **C**, Pie chart in the top panel shows percentages of genes that exhibit similar or more moderate differential expression (grey) following combinatorial treatment in comparison with the expected additive model, as well as genes that are more downregulated (blue) and more upregulated (red). Heat-map in the bottom panel shows over-representation analysis of those “more down” and “more up” genes with significant synergistic differential expression (FDR < 10%), ranked by significance. “More down” genes did not exhibit significant enrichment in any category. **D**, Genes within a path length of 3 from *ATP6V1A* on the BM36-PHG BN were enriched for down-regulated signals of Aβ-KD vs V-WT (GSEA normalized enrichment score = 2.3, adjusted P value = 8.3E-6). Node fill color denotes the expression change in demented brains and node border color gradient denotes the magnitude of expression change (t-statistics from −3 to 3) induced by *ATP6V1A* KD in Aβ-treated *NGN2*-neurons. Genes with known symbols are labeled.

As combination of *ATP6V1A* KD and Aβ treatment led to a greater amount of molecular expression change than individual factor perturbation, we explored potential synergistic effects between the two factors^59^. We first modeled the additive effect of differential expression in the individually modulated samples computationally. Hierarchical clustering of the log FCs of all genes for each contrast showed notable differences between the predicted and observed cumulative effects (including inverse differential expression of some genes) (**Fig. S22**). Similarly, competitive gene set enrichment analysis using 877 curated gene sets found strong enrichment of disorder and cellular stress gene sets following individual KD or Aβ treatment, while the KD showed further associations with cell death and negative correlation with neuronal function signatures. The latter was markedly amplified in the combinatorial modulation (Fig. 5B). To examine these synergistic effects in more detail, we grouped genes into synergism categories based on differential expression between the additive model and the combinatorial modulation. Genes were classified as “more” differentially expressed in the combinatorial modulation than predicted, if their logarithmic fold change differed by at least the average standard error in all samples. Most genes were altered approximately as predicted or less, while 6% (1152 genes) were more downregulated and 9% (1773 genes) were more upregulated than expected (Fig. 5C). Overrepresentation analysis in our curated gene sets revealed that genes more upregulated than expected from an additive model were significantly enriched for cell death and cellular stress gene sets (Fig. 5C).

The genes in response to *ATP6V1A* KD and Aβ perturbation were significantly enriched in the human postmortem brain LOAD signatures identified from the current MSBB cohort as well as 10 published datasets (Fig. 5A and **Table S18**). The down-regulated genes identified in human LOAD patients tended to show down-regulation in the current perturbation settings, while the up-regulated gene signatures identified in human LOAD patients were up-regulated in the current perturbation settings. We further overlaid the *ATP6V1A* KD and Aβ challenge perturbation profiles onto the BM36-PHG MEGENA modules and BN. Using GSEA, the top ranked neuronal/synaptic modules (M64 which contains *ATP6V1A*, M62, M65, M6, M236, M252, M385, M87, and M243) were down-regulated in KD-Aβ cells compared to WT-V cells (**Table S19**). Meanwhile, several immune response modules (M14, M153, M366 and M428) were up-regulated in KD-Aβ cells compared to WT-V cells. As summarized in **Table S20** and exemplified in Fig. 5D, the genes surrounding *ATP6V1A* on the BM36-PHG BN were enriched for down-regulation signals of *ATP6V1A* KD with or without Aβ treatment, with the most significant enrichment coming from KD-Aβ vs WT-V (FDR 4.1E-6). In summary, the *ATP6V1A* deficit signature in *NGN2*-neurons is consistent with the predicted topological structures in the human LOAD gene networks.

## DISCUSSION

Conventional largest scale GWAS have identified ~30 genetic susceptibility loci associated with the risk of LOAD ^2–5^, but little progress has been made in translating these genetic associations into therapeutics, reflecting the low penetrance of individual pathogenic factor and the lack of knowledge regarding causal mechanisms. Here we generated multi-*Omics* data from multiple brain regions of a large number of LOAD and control subjects. This unique MSBB-AD cohort enabled region-wide as well as cross-region analyses of genomic changes in LOAD brains. We identified up to 126,799 cis-associated SNP-gene pairs in region-wide analysis, majority (66.1~90.7%) of which were shared between brain regions (supplementary text). However, less than 6% of the region-wide DEGs were detected to be *cis-*regulated (FET P value > 0.1), suggesting a lack of detectable *cis*-genetic regulation among the genes dysregulated in LOAD brains. This highlights the challenges in identifying the genetic regulator that may drive the transcriptomic response in LOAD. Yet, we observed a marginally significant enrichment for LOAD genetic association signals at the *cis*-regulating SNPs (**Table S13**), warranting further investigation to link eQTLs to GWAS hits for a better interpretation of the clinical and biological relevance of the GWAS signal. In this study, *cis*-eQTLs were detected at two AD GWAS loci, including one region at the *HLA* locus and another region near gene *ZCWPW1*. A fine-mapping analysis which integrates eQTLs and GWAS signals with a summary-data-based Mendelian randomization (SMR) ^61^ method suggests that *HLA-DRB1* and *HLA-DRB6* are the most plausible functionally relevant targets underlying the GWAS hits at the HLA locus and *PVRIG* is supported as the most likely target mediating the GWAS effect at the *ZCWPW1* locus. The present analysis shows that prioritized genes may not be necessarily the genes nearest to the peak SNP as reported in the association studies, indicating caution interpreting the reported candidate risk genes by GWAS and potential power of using transcriptomic data to dissect the complex genetic association signals.

The MSBB cohort contains a continuous spectrum of LOAD-related neuropathology and clinical dementia symptom, allowing for detection of molecular changes at both early and late stage of LOAD. We built coexpression networks and BNs using all samples with an aim to maximize the power to identify the most coherent regulatory relationships across disease stages. Our integrative network analyses prioritized functional pathways (gene modules) by considering module association with the cognitive/neuropathological traits and enrichment for LOAD-related DEGs. The top-ranked modules are related to neuronal system/synaptic transmission, immune response, ECM and matrisome related, cell adhesion, etc, which is in line with our prior knowledge about the molecular feature of LOAD including reduced neuronal activity or loss of synapses^62^, alterations in synaptic adhesion^63^, dysfunction in the immune system^64^, and changes of ECM structure^31,32^. Interestingly, a number of LOAD genetic candidate risk genes were present in these top ranked modules, including *MEF2C* (M62), *CELF1*, *MADD*, *PLD3*, *PTK2B*, and *ZCWPW1* (M6), and *APP* and *SORL1* (M64), *CLU* and *CR1* (M17), and *APOE*, *CASS4*, *CD33*, *HLA-DRB1*/*HLA-DRB5*, *INPP5D*, *MS4A4A*/*MS4A6A* and *TREM2* (M153 and M14). The mechanism underlying the clustering of GWAS risk genes in the top modules is unknown. One possible reason is that they express in common cell types, especially those microglia-specific genes *CASS4*, *CD33*, *HLA-DRB1*/*HLA-DRB5*, *INPP5D*, *MS4A4A*/*MS4A6A* and *TREM2*.

One of the primary goals of this paper was to discover novel pathways and genes that are central to LOAD and could be potentially pursued with therapeutic prospects. We took advantage of the coexpression networks to prioritize modules and then used BNs to predict key driver nodes that regulated a substantial network of genes disrupted in LOAD brains. By identifying process initiators (i.e. key drivers) and reactive genes, it is possible to unravel the key genetic elements modulating pathogenic molecular mechanisms^65^. Thus, we hypothesize that key drivers play prominent roles in regulating biological pathways underlying disease pathogenesis and that manipulating the key drivers might reverse the aberrant pathway activities which in turn holds promise to reverse the disease phenotypes. The integrative network analysis-based target nomination method complements the conventional genetic linkage and linkage disequilibrium-based gene mapping technique in identifying the most functionally relevant genes that could be followed in functional studies. As 9 of the top 25 modules were related to neuronal system/synaptic transmission, we highlighted such modules to be of particular relevance to LOAD pathology and clinical severity, and predicted key molecular regulators of the modules using BNs, and one top driver, *ATP6V1A*, was tested experimentally for its disease relevance.

*ATP6V1A*, an ATP-dependent proton pump, is well known for its role in the acidification of intracellular compartments such as the lysosome; morpholino-knockdown of *ATP6V1A* impaired acid secretion in zebrafish^66^, while siRNA-mediated knockdown induced autophagy activity in U87-MG cells^67^, and KD of *ATP6V1A* in HeLa cells prevented drug-induced lysosomal acidification and autophagy activation^50^. Under our experimental conditions, *ATP6V1A* CRISPRi in hiPSC neurons did not alter lysosomal acidification (data not shown) or impact autophagy-related gene pathways. Instead, *ATP6V1A* CRISPRi down-regulated neuronal activity-associated functional pathways, particularly in the presence of Aβ42 peptides. Similar results were obtained from transgenic fly models: mRNA expression levels of fly orthologs of *ATP6V1A*, *Vha68-1*/*Vha68-2*, were reduced and neuronal KD of *Vha68-1* exacerbated age-dependent behavioral deficits and neurodegeneration accompanied by downregulation of synaptic genes in Aβ42 flies, suggesting evolutionary conserved roles of *ATP6V1A* in maintaining neuronal activity and integrity. Mounting evidence suggests that *ATP6V1A* may play an additional and unique role in synapse function; although *de novo* heterozygous mutations (p.Asp349Asn and p.Asp100Tyr) in *ATP6V1A* in rat hippocampal neurons revealed contradictory effects on lysosomal acidification, but both mutations lead to abnormalities in neurite outgrowth, branching and synaptic connectivity^51^. The synaptic role of *ATP6V1A* in LOAD brains requires further investigation.

hiPSC-based models recapitulate disease-relevant features, gene expression signatures, and identify deregulated genes with potential clinical implications (reviewed in^68^). Induced neurons also possess age-related signatures that share similarities with the transcriptomic aging signatures detected in postmortem human brain samples (reviewed in^69^). Likewise, *ATP6V1A* deficit signatures in MSBB post-mortem AD cohort were conserved in *ATP6V1A*-deficient *NGN2*-neurons. Here, we show that *ATP6V1A* KD signatures in hiPSC-neurons were highly enriched for LOAD DEGs and the sub-network surrounding ATP6V1A, indicating that hiPSC modeling is a unique and promising avenue to success for devastating diseases such as LOAD when living tissues are not available.

The present transcriptomic profiles were generated from bulk brain tissues which were mixtures of different cells of different types. Since variation in cell type proportions across individuals can influence the expression pattern, a limitation with the current transcriptomic data is that the expression level change in response to external stimuli or disease state may be confounded by cell type composition. With the recent technology advancement in single-cell gene expression analysis, including single-nucleus RNA-seq (snRNA-seq), it is now possible to study diseased tissues at the single-cell level^72^. We compared our bulk-tissue derived gene profiles with a recent snRNA-seq analysis of dorsolateral prefrontal cortex (DLPFC) region of LOAD patients and controls ^28^ and found significant preservations of our gene signatures in the snRNA-seq DEGs with the same direction of change (**Fig. S4**), despite different brain regions analyzed. This suggests that cell type proposition may have a limited impact on our gene signatures identification. Nonetheless, it is widely accepted that there exists selective regional and cell type-specific vulnerability to LOAD^8,73^. We anticipate generating region-specific single-cell multi-Omics data of LOAD and developing cell type-specific network models will offer invaluable in-depth understanding of the cellular complexity and etiology underlying the devastating disease.

In summary, we systematically identified molecular signatures, constructed multiscale gene networks and uncovered regulators of LOAD in four brain regions. We uncovered a number of relatively independent, neuron/synaptic transmission enriched gene subnetworks that were highly dysregulated in LOAD. We validated one predicted top key driver of the dysregulated neuronal system, *ATP6V1A, in silico, in vitro* and *in vivo*, and demonstrated *ATP6V1A* to be a promising therapeutic candidate target for treating LOAD.

## Funding

This work was supported in parts by grants from the National Institutes of Health (NIH)/National Institute on Aging (R01AG046170, RF1AG054014, RF1AG057440, R01AG057907, U01AG052411, R01AG062355, U01AG058635), NIH/National Institute of Allergy and Infectious Diseases (U01AI111598), NIH/National Institute on Drug Abuse (R01DA043247), NIH/National Institute of Dental and Craniofacial Research (R03DE026814), NIH/National Institute of Diabetes and Digestive and Kidney Diseases (R01DK118243), Dept. of the Army (W81XWH-15-1-0706), Japan Society for the Promotion of Science KAKENHI (JP16K08637), and the Research Funding for Longevity Science from the National Center for Geriatrics and Gerontology, Japan (grant numbers 19-49 to MS and 19-7 to KI). All hiPSC research was conducted under the oversight of the Institutional Review Board (IRB) and Embryonic Stem Cell Research Overview (ESCRO) committees at Icahn School of Medicine at Mount Sinai (ISSMS). Informed consent was obtained from all skin cell donors as part of a study directed by Judith Rapoport MD at the National Institute of Mental Health (NIMH).

## AUTHOR CONTRIBUTIONS

BZ and ES conceptualized the study. BZ designed the study; ES, KI, and KB participated in the study design and the discussion of the results; MW, AL, MS, NB, and XQ performed research; NS, RN, YW, WS, EW, QS, XZ, LG, CM, SV, EH, JZ, SG, ME, and VH participated in data analysis and interpretation; LL, AY, MF, PS, SH, LZ, JC, QW, HK, JJ, MZ, KA, DC, and YZ, contributed to validation experiments. MW, AL, MS, KI, KB, and BZ wrote the paper. All the authors reviewed and revised the paper.

## DECLARATION OF INTERESTS

The authors declare no competing interests.

