## Supplementary text and figures for "Molecular Networks and Key Regulators of the Dysregulated Neuronal System in Alzheimer’s Disease"

### Supplementary information

#### 1 Human postmortem brain Omics data analysis

##### 1.1 MSBB RNA-seq data quality control and preprocessing

Through an iterative QC and adjustment procedure which examined the genetic similarity between every pair of molecular profiles across different data types and multiple brain regions, we identified mislabeled or duplicated molecular profiles [1]. In this paper, we excluded all mislabeled samples for downstream analyses. For RNA-seq, we further removed RNA-seq libraries with RNA integrity number (RIN) less than 4 or rRNA rate larger than 5%, and then selected one with the best sequencing coverage for the duplicated sequencing libraries (see table below for demographics of RNA-seq samples). To avoid any artificial regional difference, the data from all four brain regions were merged and processed together. Genes with at least 1 count per million (CPM) reads in at least 10% of the libraries were considered expressed and hence retained for further analysis; others were removed. After filtering, 23,201 genes were retained. The gene read counts data were normalized using the trimmed mean of M-values normalization (TMM) [2] method in the R/Bioconductor edgeR package to adjust for sequencing library size differences. It is critical to identify and correct for confounding factors in the RNA-seq data. For this purpose, we used R/Bioconductor variancePartition [3] package to evaluate the impact of multiple sources of biological and technical variation in gene expression experiments, including sex, race, age, RIN, postmortem interval (PMI), sequencing batch, rate of exonic reads, and rate of rRNA reads, together with the four cognitive/neuropathological features described in the main text. **Fig. S1** illustrates the principal component analysis and variance partition analysis of the RNA-seq data. We found sequencing batch, exonic rate and brain donor contributed to the most variance. The contributions from the cognitive/neuropathological variables were similar and ranked in the middle among all the variables. While rRNA rate generally did not explain a large proportion of variation, it contributed more overall variance than did sex and race. Therefore, in addition to the usual confounding factors that are commonly corrected in postmortem brain gene expression data, including batch, sex, race, age, RIN, and PMI, we included exonic rate and rRNA rate as covariates. As there were more than 30 batches, the batch was firstly regressed out with a random effect model using variancePartition [3], and the other covariates were corrected by linear regression in R.

**Demographics of the MSBB RNA-seq samples after quality control processing.**

| Sample size or age | Brain region | | | |
|---|---|---|---|---|
| | **BM10-FP** | **BM22-STG** | **BM36-PHG** | **BM44-IFG** |
| Total (male %) | 261 (35.6%) | 240 (37.5%) | 215 (38.1%) | 222 (36%) |
| Age (years ± s.d.) | 85.2±9.6 | 85.0±9.6 | 85.0±9.7 | 85.3±9.7 |
| Cognitive normal (CDR = 0) | 35 | 33 | 32 | 27 |
| Mild cognitive impairment (CDR = 0.5) | 39 | 33 | 32 | 38 |
| Demented (CDR >= 1) | 187 | 174 | 151 | 157 |

#### 1.2 Differential expression analysis

For each neuropathological/cognitive trait in each brain region, we grouped the samples into multiple disease severity stages and compared the gene expression between every two groups using limma's moderated t-test analysis[4]. Specifically, for CDR, samples were classified into cognitive normal (nondemented) (CDR = 0), mild cognitive impairment (MCI) (CDR = 0.5), and demented (CDR ≥ 1). For Braak score, samples were classified into normal (NL) when Braak score ≤ 2, and AD when Braak score > 2. For plaque mean density (PlaqueMean), samples were classified into 4 categories, namely normal (PlaqueMean = 0), mild (0 < PlaqueMean ≤ 6), medium (6 < PlaqueMean ≤ 12), and severe (PlaqueMean > 12) groups. With CERAD score, two types of samples classification schemes were used. First, samples were classified into normal (NL) (CERAD = 1), definite AD (CERAD = 2), probable AD (CERAD = 3) and possible AD (CERAD = 4). Second, samples were classified into two groups, normal (NL) when CERAD = 1 and AD when CERAD > 1. To adjust for multiple tests, false discovery rate (FDR) was estimated using the Benjamini-Hochberg (BH) method[5]. Genes showing at least 1.2-fold change (FC) and FDR adjusted P values less than 0.05 were considered significant. The gene showing the largest fold increase in all comparisons is *LTF* (lactotransferrin) (3.8-fold, adjusted P value 3.9E-5) as identified in BM36-PHG with respect to the PlaqueMean trait. Lactotransferrin is a major component of mammals' innate immune system, protecting from direct antimicrobial activities to anti-inflammatory and anticancer activities[6]. *NEUROD6* (neuronal differentiation factor 6) showed the largest fold decrease across all contrasts (0.34-fold, adjusted P value = 6.3E-9). *NEUROD6* encodes a transcription activator that may be involved in neuronal development and differentiation. Down-regulation of *NEUROD6* in LOAD has been consistently observed in several previous studies [7,8].

To systematically validate the present DEG signatures of LOAD related traits, we assembled public ALOD signatures from 10 studies, including Zhang et al 2013 [9], Webster et al 2009 [10], Satoh et al 2014 [8], Miller et al 2013 [11], Avramopoulos et al 2011 [12], Liang et al 2008 [13], Colangelo et al 2002 [14], Blalock et al 2004 [15], Mostafavi et al 2018 [16], and Allen et al 2018 [17]. Then we evaluated the overlap between the present DEGs and these previously published LOAD signatures using the Fisher's exact test (FET). We observed a highly significant overlap (adjusted P value up to 1.0E-100) for almost every differential contrast in public LOAD signatures as illustrated in **Fig. S2**. We note that when up- and down-regulated DEGs were separated, we observed significant enrichments in consistent directions with respect to expression changes in this analysis. The relatively mild enrichment for the signatures in BM10-FP and BM44-IFG was due to the small number of genes identified in the two regions. To further investigate if the present expression signatures from bulk tissue RNA-seq tend to reflect cell-type changes, we collected a set of cell type-specific DEGs identified from a recent single-nuclei RNA-seq (snRNA-seq) analysis of LOAD postmortem brains [18]. Here, we used cell type-specific DEGs computed from the cell-level model. **Fig. S3** shows the FET of the enrichment between our bulk-tissue DEGs and the cell type-specific DEGs detected in Ex (excitatory neurons), In (inhibitory neurons), Oli (oligodendrocytes), Opc (oligodendrocyte progenitor cells), Ast (astrocytes), or Mic (microglia) in brains with LOAD pathology. We observed a strong preservation of both up- and down-regulated genes in a cell type-specific manner. These results demonstrate a robust set of LOAD related gene signatures across all brain regions profiled.

To understand what biological processes are represented in the DEGs, we tested these signatures for enrichment of gene ontology (GO) and canonical functional pathway gene sets from the Molecular Signatures Database (MSigDB) gene annotation database v6.1 [19,20]. For convenience, the MSigDB gene set collections have been assembled into an R package called "msigdb" which is publicly available from https://github.com/mw201608/msigdb. We overlapped the DEGs with the MSigDB gene sets and computed the fold enrichment (FE) and P value significance using the algorithms described in the next section "Functional enrichment analysis". The top GO and functional pathways identified are summarized in **Fig. S4,** and the full list of significant enrichments is provided in **Table S2**.

#### 1.3 Functional enrichment analysis

Functional enrichment analysis (or overlap test) P value was calculated using the hypergeometric test (equivalent to the Fisher's exact test, FET) assuming the sets of genes, such as DEGs, were identically independently sampled from the genome-wide genes profiled. Under the same assumption, FE was calculated as the ratio between observed overlap size and expected overlap size. To control for multiple testing, we employed the Benjamini-Hochberg (BH) approach [5] to constrain the FDR. For GO and pathway enrichment analysis, we utilized the functional gene set collections from the Molecular Signatures Database (MSigDB) v6.1 [19,20]. For brain cell type marker gene enrichment analysis, we focused on the 5 major brain cell types, i.e. neurons, microglia, astrocytes, oligodendrocytes and endothelial, and for each type used the top 500 ranked consensus cell type-specific genes derived from a meta-analysis of 5 cell type-specific or single cell RNA-seq datasets [21].

#### 1.4 Region-region correlation analysis

We investigated molecular interactions between brain regions by using Pearson's correlation analysis. There were 6 distinct pairs of brain regions in our dataset, each comprised of 215-240 RNA-seq matched samples. Using Pearson's correlation analysis, between 176,453 and 1,159,477 (0.03~0.22%) gene pairs showed significant inter-region correlations across the 6 brain region pairs at a conservative Bonferroni corrected P value threshold of 0.05 (**Fig. 1C**). There was a relatively smaller number of correlated genes between BM36-PHG and BM10-FP and between BM36-PHG and BM44-IFG, possibly due to the distal physical interconnectivity from BM36-PHG to these two regions. The correlations for the same genes between any two regions ranged from -0.19 to 0.97 (mean = 0.31 ~ 0.38, median = 0.29 ~ 0.38), indicating the presence of both highly self-correlated genes and inversely-correlated genes. The later highlights the importance of using the right tissue to study a given biological process. Excluding the gene pairs from the same genes, the number of the positively correlated gene pairs was 3.3~6.7-fold of that of the negatively correlated gene pairs. 12,013 gene pairs were significantly correlated across all 6 pairwise inter-region correlation analyses (**Table S3**). Immune response pathways (adjusted P value = 1.12E-59, 4.39-fold enrichment (FE)) were most enriched in the gene pairs with positive inter-region correlations (**Table S4**, **Fig. 1C**). On the other hand, gene pairs showing strong negative inter-region correlations were related to cellular response to unfolded protein (UPR), including protein folding (8.76-FE, adjusted P value = 1.52-E13), unfolded protein binding (12.93-FE, adjusted P value = 1.57E-11) and protein refolding (29.95-FE, adjusted P value = 9.71E-9). Using Spearman's correlation coefficient analysis, 5 UPR genes, *BAG3*, *SERPINH1*, *CDKN1A*, *HSPA6*, and *SOCS3*, were found to be positively correlated with CDR (correlation coefficient r > 0.2), and 4 UPR genes, including *HSPA4L*, *CIRBP*, *PPID*, and *HSPH1*, were negatively correlated with CDR (correlation coefficient r < -0.2), at adjusted P value ≤ 0.05. We confirmed that inter-region correlated gene pairs in the immune response and UPR pathways, were also correlated within the corresponding regions and in the same direction, suggesting that immune response and UPR are common and coordinated molecular activities of the four brain regions.

#### 1.5 MEGENA gene coexpression network analysis

For MEGENA [22], Pearson correlation coefficients (PCCs) were computed for all gene pairs in every brain region. Significant PCCs at a permutation-based FDR cutoff of 0.05 were ranked and iteratively tested for planarity to grow a Planar Filtered Network (PFN) by using the PMFG algorithm. Multiscale Clustering Analysis (MCA) was conducted with the resulting PFN to identify coexpression modules at different network scale topology. We identified 475, 527, 441 and 423 coherent gene expression modules in BM10-FP, BM22-STG, BM36-PHG and BM44-IFG, respectively (**Table S5**). To annotate the potential biological functions associated with the modules, we performed MSigDB gene set enrichment analysis using FET as described above. Most of these modules (53.9% to 67.3%) were enriched for MSigDB GO/pathway gene sets (adjusted P value < 0.05) (**Table S6**), indicating that MEGENA is capable of capturing data-drive biologically meaningful, context-dependent co-regulation signals beyond what is represented in canonical pathways from ontology databases. For simplicity, modules were annotated by the top enriched functional category. It is noted that MEGENA modules are formed in a hierarchy with parent-child relationships.

We found many top ranked modules were enriched for neuronal or microglia-specific cell types (**Fig. 2E**) based on enrichment analysis of recently determined cell type-specific markers of five major brain cell types, including neurons, microglia, endothelial, astrocytes and oligodendrocytes [21]. To test whether the top ranked synaptic transmission/neuronal system modules reflected distinct neuronal subtypes, we utilized a large-scale single-nucleus RNA-seq data of inhibitory and excitatory neurons isolated from six different regions of the human cerebral cortex [23]. We downloaded the preprocessed gene expression data (transcripts per million, TPM) from this published study and further selected genes with at least 1 TPM in at least 10% of the cells in one subtype. Then we computed genes which showed differential expression between cell types using limma's moderated t-test analysis [4]. Source of brain region origination of the cells was incorporated as a covariate. While the original study identified up to 16 different subtypes of neurons, we focused on the genes differentiate between the two major neuron cell types (i.e. inhibitory and excitatory). We called genes inhibitory neuron-enriched if they presented at least 4-fold higher expression in inhibitory neuron cells than in excitatory neuron cells with FDR < 0.05, and excitatory neuron-enriched if they presented at least 4-fold higher expression in excitatory neuron cells than in inhibitory neuron cells with FDR < 0.05. As a result, we identified 1008 excitatory neuron-enriched genes and 413 inhibitory neuron-enriched genes. Lastly, we overlapped the cell type-enriched genes with the top ranked synaptic transmission/neuronal system modules and found that module M64 was overrepresented with inhibitory neuron-enriched genes while M6, M87, M65, M236, M62, and M252 were overrepresented with excitatory neuron-enriched genes (**Table S8**).

#### 1.6 Discovery of region-wide expression quantitative trait loci (eQTLs)

Given the well-established relationships between gene expression and interactions with genetic and environment factors, we mapped expression quantitative trait loci (eQTLs) by integrating the RNA-seq and WGS-based SNP genotype data. SNPs significantly associated with gene expression traits were identified using the MatrixEQTL package [24]. Significant SNPs (eSNPs) were classified into *cis*- and *trans*-acting elements according to whether they are located within 1-MB from the gene or not. At a conservative Bonferroni corrected P value threshold of 0.05 (equivalent to a nominal P value cutoff of 3.0E-10), 1214, 922, 762, and 1054 genes were identified to be regulated by at least one proximal SNP within 1 million base (Mb) from the gene, termed *cis*-eSNP, in BM10-FP, BM22-STG, BM36-PHG, and BM44-IFG, respectively (**Table S9**). For simplicity, we refer to genes with significant eSNPs as eGenes and a significant association between a SNP and a gene as an eSNP-eGene pair. By such a definition, 126,799, 101,705, 92,336, and 112,139 *cis*-eSNP-eGene pairs were identified in BM10-FP, BM22-STG,

BM36-PHG and BM44-IFG, respectively. It is noted that there are redundant eSNPs for the same eGene due to linkage disequilibrium (LD) of the SNPs. **Fig. S8** shows the overlap of these *cis*-eSNP-eGene pairs among the four brain regions. 66.1% to 90.7% of the *cis*- eSNP-eGene pairs identified in one brain region were also detected in at least one other brain region. In addition, 71,298 *cis*-eSNP-eGene pairs from 548 unique genes were shared by all 4 brain regions.

We detected 20,657, 14,011, 14,766, and 17,125 *trans*-eSNP-eGene pairs from BM10-FP, BM22-STG, BM36-PHG and BM44-IFG, respectively. For each brain region, 28.5 to 70.1% of the *trans*-eSNP-eGene pairs identified were also detected in at least one other brain region (**Fig. S8**). We grouped Bonferroni corrected significant SNPs within a 5-Mb interval into a single peak because of insufficient resolution to break LD over such narrow windows[25,26]. Each peak was represented by the most significant eSNP in the window, referred to as the lead eSNP, for a given *trans*-eGene. We identified 2,411, 1,965, 1,392 and 2,460 *trans*-eQTL peaks from BM10-FP, BM22-STG, BM36-PHG and BM44-IFG, respectively. Early eQTL studies noted the existence of master *trans*-genetic regulators, which we refer to as eQTL hotspots [27], that regulate many genes throughout the human genome. We defined *trans*-eQTL hotpots as those peaks associated with 10 or more *trans*-eGenes. From this definition we identified 24, 12, 2 and 27 *trans*-eQTL hotpots from BM10-FP, BM22-STG, BM36-PHG and BM44-IFG, respectively (**Fig. S9** and **Table S10**), with nine *trans*-eQTL hotspots shared between 2 or 3 brain regions (**Fig. S10**). Each of these hotspots were associated with 10 to 36 *trans*-eQTL genes. The hotspot associated with the greatest number of *trans*-eQTL genes (36 genes) was located at a region near 84.4-Mb on chromosome 17 (lead eSNP rs10264300) in BM44-IFG. SNP rs10264300 is 181 kilobases upstream of *AC003984.1* (a long intergenic noncoding RNA, lincRNA) and 82.5 kilobases downstream of *AC093716.1* (a pseudogene gene). About half (16) of the gene targets of this hotspot encode enzyme binding proteins (6.4-FE, adjusted FET =7.1E-5) (**Table S11**). Interestingly, synaptic pathway genes were enriched for the targets of a hotpot near lead SNP rs34072069 on chromosome 10 in BM10-FP (17.1-FE, adjusted FET P=4.9E-6). SNP rs34072069 is 44.5 kilobases upstream of RNU6-535P (a small nuclear RNA gene) and 1.9 kilobases downstream of RP11-385N23.1 (an antisense gene).

We evaluated whether any modules were enriched for our cis-eGenes. Twelve MEGENA modules were significantly enriched for *cis*-eQTL genes (**Table S12**), among which four were associated with GTPase mediated signal transduction (one from each brain region (>17.9-FE, adjusted FET P < 3.5E-11) and three were associated with transferase activity (one from each of 3 brain regions except BM44-IFG; > 10.9-FE, adjusted FET P < 2.3E-4). We noted that the genes in the GTPase mediated signal transduction modules were concentrated in chromosome region 17q21, while the transferase activity modules in chromosome region 8p23, suggesting the genetic regulation of these modules by common eQTLs shared by multiple brain regions.

We attempted to replicate eQTLs in an independent LOAD postmortem brain RNA-seq dataset generated from the ROSMAP cohort [16], which is, to our knowledge, the largest sampled RNA-seq based eQTL analysis of LOAD in a single brain region (494 individuals). *Cis*-eQTLs were identified for 3,388 genes from the ROSMAP cohort as published by Ng et al[28]. However, since no *trans*-eQTLs were reported for the ROSMAP cohort[28], we focused on the replication of *cis*-eQTLs in this paper, and particularly the *cis*-eSNP-eGene pairs that were available in both datasets. To avoid including dependent signals induced by LD among adjacent SNPs, only the associations comprising the top SNP for each eGene were included in the replication rate calculations. To circumvent the statistical power difference caused by different sample sizes (494 individuals in ROSMAP and 215~261 individuals across the present four brain regions), we first followed Ng et al [28] to assess the replication rate of LOAD brain *cis*-eSNP-eGene discovered in the ROSMAP cohort in our data set using the $\pi_1$ statistic [29], which estimated the proportion of reported ROSMAP *cis*-eSNP-eGene pairs that are also significant in the current data set based on their P-value distribution. $\pi_1$ values of the ROSMAP *cis*-eSNP-eGene pairs were 0.698, 0.674, 0.637, and 0.670 in the present brain regions BM10-FP, BM22-STG, BM36-PHG, and BM44-IFG, respectively. These values were significantly larger than their empirical null mean of 0.025~0.038 from 10,000 random samples of P values of associations that did not overlap with the eQTLs (one-tailed P value < 0.0001). Analogously, we applied the same $\pi_1$ statistic to estimate the replication rate of the present region-wide eQTLs in the ROSMAP data but were unsuccessful because the P value distributions of the MSBB *cis*-eSNP-eGene pairs were truncated (maximum P value = 0.92) with majority of the values approaching 0 (93% to 96% were less than 0.05) in the ROSMAP data. In fact, 81.6%, 84.3%, 89.0% and 82.5% of the *cis*-eSNP-eGene pairs identified in BM10-FP, BM22-STG, BM36-PHG, and BM44-IFG, respectively, were also called genome-wide significant in the ROSMAP data, thus indicating most of the present cis-eQTLs were replicated. These results indicate marked common genetic regulation occurring across different brain regions.

#### 1.7 Integrating eQTL, gene expression traits, and LOAD GWAS loci to identify causal LOAD genes

We did not observe significant enrichment for *cis*-eGenes in the LOAD-related DEGs or brain cell type-specific markers in each brain region (with less than 8% of the *cis*-eGenes detected as DEGs and less than 6% of the DEGs detected as *cis*-eGenes, FET P value > 0.1), suggesting a lack of detectable *cis*- genetic regulation among the genes dysregulated in LOAD brains. However, for the most strongly associated SNPs across all *cis*-eGenes, we observed a significant enrichment for LOAD genetic association signals based on the SNP-level summary statistics from IGAP [30] (http://web.pasteur-lille.fr/en/recherche/u744/igap/igap_download.php) ($P < 0.05$), compared to random samples of SNPs of the same size (**Table S13**). In this analysis, we first selected the strongly associated SNPs across all *cis*-eGenes and then extracted their SNP-level LOAD GWAS chi-square statistics from IGAP [30]. The mean chi-square statistics among those *cis*-eSNPs was compared to a null distribution which was obtained by randomly sampling the same number of SNPs for 10,000 times. Enrichment P value was computed as the proportion of randomly sampled SNP sets with mean chi-squared values larger than the observed one.

Moreover, *cis*-eQTLs overlapped the genome-wide significant LOAD GWAS SNPs at GWAS risk loci *HLA-DRB1*/*HLA-DRB5* and *ZCWPW1*. To aid in the identification of candidate causal genes in these GWAS loci, we applied the summary-data-based mendelian randomization (SMR) [31] method to test if the effects of the top GWAS SNPs in the *HLA-DRB1*/*HLA-DRB5* and *ZCWPW1* loci were mediated by gene expression associated with eQTL coincident with the GWAS loci. By integrating eQTLs and GWAS signals, we aimed to prioritize the most possible functional relevant genes underlying the effects of causal variants on the disease phenotype at two LOAD GWAS risk loci. We reformatted the eQTL results and IGAP SNP-level summary statistics data files in accordance to the manual of the SMR software [31]. Then we ran region-wide SMR analysis using the default parameter. For each locus, we used the region-wide rather than the experiment-wise significance threshold because we were interested in gene discovery for each specific locus in each brain region than the joint analysis of all regions as a whole. For the genes with significant association by the SMR test, the heterogeneity in dependent instruments (HEIDI) test [31] was further employed to distinguish whether the association was caused by pleiotropy of the same causal variant underlies the disease risk, or due to linkage of distinct variant to the one causal to the disease.

**Fig. S10** shows the GWAS and eQTL P value profiles at the *HLA-DRB1*/*HLA-DRB5* locus as well as the SMR test results in four brain regions. In a 2-Mb region centered on *HLA-DRB1*, there were 8 to 11 genes with cis-eQTLs across the four brain regions. For example, 11 genes were found to have *cis*-eQTLs in BM10-FP; the SMR test was significant for 5 of these genes at a Bonferroni corrected P value threshold of 4.5E-3, including *HLA-DRB1*, *HLA-DRB6*, *HLA-DQA1*, *HLA-DQA2*, and *HLA-DQB2*, while the gene *HLA-DRB5* was not significant. To distinguish whether the significant association in the SMR test was caused by pleiotropy that gene expression and the trait affected by the same underlying causal variant, or due to linkage that the top associated *cis*-eQTL being in LD with two distinct causal variants, one affecting the disease trait and the other affecting the gene expression, we further performed the heterogeneity in dependent instruments (HEIDI) test as in Zhu et al [31]. Of the 5 genes surpassing the SMR test in BM10-FP, 3 genes, including *HLA-DRB1*, *HLA-DRB6* and *HLA-DQA1*, showed no significant heterogeneity by the HEIDI test (P value > 0.05), supporting the null hypothesis that there is a single causal variant affecting both gene expression and disease trait phenotype. Using the same analytic procedure, we found that *HLA-DRB6* passed both SMR and HEIDI tests in all the other three brain regions, and *HLA-DRB1* passed both SMR and HEIDI tests in BM36-PHG and BM44-IFG (**Fig. S10**), suggesting that the *HLA-DRB1* and *HLA-DRB6* are the most plausible functionally relevant targets underlying the GWAS hits at this locus.

At a 2-Mb region surrounding gene *ZCWPW1*, there were 3 to 6 genes with *cis*-eQTLs across the different brain regions (**Fig. S11**). *ZCWPW1* did not pass the SMR test, indicating that our data do not support that the expression of *ZCWPW1* mediates the causal effect on the disease phenotype. However, an adjacent gene *PVRIG* passed the SMR test in all brain regions except BM44-IFG, and another adjacent gene *ZSCAN21* passed the SMR test in BM22-STG. *PVRIG*, also known as *CD112R*, encodes a protein that recruits tyrosine phosphatases for signal transduction and could act as a coinhibitory receptor that suppresses T cell receptor (TCR) signaling [32]. *ZSCAN21* is a transcription factor containing zinc finger and SCAN domains and may regulate alpha-synuclein (SNCA) in primary neuronal cultures [33,34]. We noted that only *PVRIG* passed the HEIDI test in BM10-FP, indicating that in our data *PVRIG* is supported as the most likely target mediating the effect of the LOAD-associated variant on disease compared to other genes in this region.

The present analysis shows that prioritized genes may not be necessarily the genes nearest to the peak SNP as reported in the association studies. Further independent replications and experimental validations are required to verify the potential causal relationships inferred from the current integrative analysis.

##### 1.8 Bayesian probabilistic causal network inference and key driver analysis

To construct Bayesian probabilistic causal Network (BN), we made use of genetic perturbations in biological systems (e.g. WGS SNP variants) and known transcription factor (TF)-target relationships from the ENCODE project as prior for inferring regulatory relationships between genes. In the causal network construction, the TFs are allowed to be parent node of their target genes; but targets are inhibited to be parent nodes of their TFs. To infer gene regulatory relationship from genetic data, we first computed *cis*- and *trans*-eQTLs for each expression trait using WGS-based SNP variants as described above and then employed a causal inference to infer the causal probability between gene pairs associated with the same eQTL. Since a gene pair associated with the same eSNP may be causally regulated from one to another or independently regulated by a genetic factor in LD with the eSNP, we derived genetic priors under two scenarios. In the first scenario, genes with *cis*-acting eSNP could be parent nodes of genes with *trans*-acting eSNP, but the opposite direction was not permitted following previous practices [35,36]. In the second scenario where the genes are both *cis*-regulated or both *trans*-regulated, either gene can be the parent node of the other and hence there are two possible directions. For the latter scenario, we applied a formal causality inference test (CIT) [37,38] to distinguish the causal/reactive and independent relationships between the gene expression traits by modeling the gene pair and associated eSNP with a "chain" of mathematical conditions. For each trio (a gene pair and one eSNP), CIT will compute the  probability of the causal "chain" in which one gene is mediating the causal impact of the eSNP to the other gene when the regulatory direction is allowed [38]. In cases that the gene pair is associated with multiple common eSNPs, the individual causality test P values of each trio were aggregated using Fisher's method to make a collective call for the gene pair. As the conservative Bonferroni corrected P value threshold of 0.05 in the eQTL analysis gave a very limited number of gene pairs associated with common SNPs, we relaxed the cutoff to a BH FDR adjusted P value threshold of 0.05 to increase the pool of potential causal-reactive gene pairs. The causal relationships thus inferred by CIT were combined with TF-target relationships, and together they were used as structure priors for building a brain region-wide BN from all 23,201 expressed genes through a Monte Carlo Markov Chain (MCMC) simulation based procedure [39]. Following previous practices [35,36], we employed a network averaging strategy in which 1,000 networks were generated by this MCMC process starting with different random structure, and links that appeared in more than 30% of the networks were used to define a final consensus network. If loops were present in the consensus network, the weakly supported link involved in a loop was removed to ensure the final network structure was a directed acyclic graph.

From the region-wide BNs, we identified network key drivers that are predicted to modulate a large number of downstream nodes, and as a result, modulate the state of the network, by using the Key Driver Analysis (KDA) [9,40,41]. Here we loaded all the BN nodes as input in the KDA and hence the resulting key drivers were called global network key drivers which were different from the pathway context dependent key drivers such as the synaptic transmission/neuronal system module focused key drivers described later. There were 1,545, 1,418, 1,454, and 1,371 global key drivers in the BNs from BM10-FP, BM22-STG, BM36-PHG and BM44-IFG, respectively. Strikingly, the key drivers were significantly conserved across the region-wide BNs, with any two BNs sharing a significant number of key drivers ($7.2 < FE \leq 8.2$, FET P values < 1.0E-320) while 325 key drivers were shared across four BNs (929.4-FE, Super Exact Test P value < 1.0E-320) (**Fig. S12**), demonstrating a high-degree of conservation of the regulatory architecture in the brain regions we profiled.

To perform a comprehensive validation of the BN topological structures and the global key drivers that modulate them, we downloaded a library of 2,460 single gene perturbation signatures curated at the Enrichr server [42]. After filtering for central nervous system (CNS) or immune system-related studies and requiring the perturbed genes to be present in the current dataset, we obtained 649 signatures from 320 studies for 287 unique perturbed genes. These gene perturbation signatures were collected from the gene expression omnibus (GEO) database, and the original experiments were conducted in a diversity of conditions (different cell lines or tissues from different species). 66 of these perturbed genes were global key drivers in at least one of our four brain region-wide BNs. For each of these perturbed global key drivers, we examined whether the experimental perturbation signature was predicted by our networks by examining whether the genes in these signatures were enriched for genes in the network neighborhood of the key driver in our BNs (examining genes that were within a path length of 6 of the key driver gene). Despite the vast heterogeneity of the gene perturbation studies compared to the present human postmortem brain tissues used to generate our data for the region-wide BNs, 50 to 60% of the key driver perturbation signatures were enriched in the network neighborhoods of the corresponding key drivers across the four region-wide BNs (**Fig. 3B**). The significance levels of the enrichments are observed to increase as path lengths defining the network neighborhoods are increased, given the BNs were sparse, with a limited number of neighboring nodes closer to the key driver gene, which serves to reduce the power to make such detections, especially in the context of multiple testing. In contrast, the proportion of significantly enriched perturbation signatures decreased to 20~30% in the network neighborhood of non-driver genes.

We further performed KDA on the top ranked MEGENA synaptic transmission and neuronal system modules to identify their master regulators. In this analysis, we projected the module genes onto the region-wide BN and searched for key driver genes whose network neighborhood were enriched for the module genes. Different from the global key drivers described above, here the key drivers were context dependent, in this case, related to synaptic transmission and neuronal system. This yielded 42 unique key driver genes across 9 modules (**Table S14**) predicted to be the key drivers of the synaptic transmission and neuronal system modules. 10 key drivers were root nodes in the BM36-PHG BN without parental nodes. To further verify the root node status beyond a single region-based network, we sought to integrate information from all four region-wide BNs by building a union BN, which contained a union of directed links from all four individual BNs. Like region-wide BNs, loops in the union BN were broke by removing the weakly supported links. Two key drivers, *ATP6V1A* (module M64), and *GABRB2*, (module M62), remained as root nodes in this union BN.

Gamma-aminobutyric acid (GABA) is the major inhibitory neurotransmitter in the mammalian brain and GABA type A (GABA-A) receptors mediate the inhibition effect [43]. GABA-A receptors form pentameric complexes by combinations of more than 10 subunits and marked functional remodeling GABA-A receptors, including change of subunit composition and reduced expression of principal subunits, had been observed in LOAD brains [44]. It has been reported that GABA-A β2 (*GABRB2*) subunit, paralleled with some other subunits like α1, α2, α5, β3, and γ2, showed altered brain region- and cell layer-specific expression [45]. Its protein level was significantly decreased in the dentate gyrus stratum moleculare, but increased in the stratum oriens and stratum radiatum of the hippocampal CA2 region, and stratum radiatum of the hippocampal CA3 region [45]. In this paper, we observed a significant down-regulation of the subunits α1-6, β2-3 and γ2-3 in diseased brains compared to control (**Table S1**).

### 2 In vitro functional validation of *ATP6V1A* deficit in *NGN2*-neurons

#### 2.1 gRNA design and cloning

gRNA design and cloning were performed as previously described (*Ho et al, 2017*). Specifically, 6 gRNA candidates for ATP6V1A were designed by using CRISPR-ERA web tool (crispr-era.stanford.edu): 6 gRNA sequences targeting promoter region (between +658 bps and transcription start site) of ATP6V1A. For lentiviral cloning, the gRNA sequences were inserted into LentiGuide-Hygro-mTagBFP2 (Addgene #99374). Oligonucleotides encoding gRNA sequences were annealed, diluted and then ligated into BsmBI-digested LentiGuide vectors as previously described (*Ho et al, 2017*). Sanger sequencing using U6 promoter confirmed all constructions.

| **Oligo ID** | **Location** | **gRNA** (E,S score) | **gRNA Sequence** (5'-3') |
|---|---|---|---|
| **ATP6V1i_#1-1** | +260 | #1 (20,0) | **5'-CACCG***GCGGGAACGACCACACTTGG* |
| **ATP6V1i_#1-2** | | | **5'-AAAC***CCAAGTGTGGTCGTTCCCGC***C** |
| **ATP6V1i_#2-1** | +101 | #2 (20,0) | **5'-CACCG***GGCGACCGGTAACTGGCGAG* |
| **ATP6V1i_#2-2** | | | **5'-AAAC***CTCGCCAGTTACCGGTCGCC***C** |
| **ATP6V1i_#3-1** | +94 | #3 (20,0) | **5'-CACCG***GGTGAGCGGCGACCGGTAAC* |
| **ATP6V1i_#3-2** | | | **5'-AAAC***GTTACCGGTCGCCGCTCACC***C** |
| **ATP6V1i_#4-1** | +14 | #4 (20,-2) | **5'-CACCG***GGGGAAGTCCTCAGCTGCAC* |
| **ATP6V1i_#4-2** | | | **5'-AAAC***GTGCAGCTGAGGACTTCCCC***C** |
| **ATP6V1i_#5-1** | +266 | #5 (20,-2) | **5'-CACCG***GTGGTCGTTCCCGCTACTT* |
| **ATP6V1i_#5-2** | | | **5'-AAAC***AAGTAGCGGGAACGACCAC***C** |
| **ATP6V1i_#6-1** | +658 | #6 (15,0) | **5'-CACCG***GATGTTCACGTGCTTCGGAT* |
| **ATP6V1i_#6-2** | | | **5'-AAAC***ATCCGAAGCACGTGAACATC***C** |

#### 2.2 Lentivirus generation

Third-generation VSV.G pseudotyped HIV-1 lentiviruses (below) were produced by polyethylenimine (PEI, Polysciences #23966- 2)-transfection of HEK293T cells and packaged with VSVG-coats using established methods (Tiscornia et al., 2006). Lentiviral FUW-M2rtTA (Addgene #20342), pLV-TetO-hNGN2-eGFP-neo (TBD), lentiGuide-Hygro-mTagBFP2, and 6 lentiGuide vectors with insertion were generated. Physical titration of lentivirus was performed by qPCR (qPCR Lentivirus Titration Kit, ABM good #LV900). Lentiviruses were then used to transduce cells according to their physical titer as described below, calculated through the company's website (https://www.abmgood.com/High-Titer-Lentivirus-Calculation.html).

#### 2.3 hiPSC-NPC culture and *NGN2* neuronal differentiation

Two stable hiPSC-derived neuronal progenitor cells (hiPSC-NPCs) (553KRAB and 2607KRAB) expressing dCas9$^{-KRAB}$ (Addgene 99372) were generated as previously described[46] and cultured in hNPC media (DMEM/F12 (Life Technologies #10565), 1x N2 (Life Technologies #17502-048), 1x B27-RA (Life Technologies #12587-010), 20 ng/ml FGF2 (Life Technologies), and 0.3 µg/mL puromycin) on Matrigel (Corning, #354230). NPCs at full confluence (1-1.5x$10^7$ cells/well of a 6-well plate) were dissociated with Accutase (Innovative Cell Technologies) for 5 mins, spun down (5 mins X 1000g), resuspended and seeded onto Matrigel-coated plates at 3-5x$10^6$ cells/well. Media was replaced every two days for four to seven days until next split.

At day -2, NPCs were seeded as 4-6x$10^5$ cells/well in a 24-well plate coated with Matrigel (coverslips are put in a plate and coated with Matrigel for immunostaining). At day -1, cells were transduced with rtTA, pLV-TetO-hNGN2-eGFP-Neo and *ATP6V1A*i gRNA or empty lentiguide-Hygro-mTagBFP2 (Addgene 99374) lentiviruses via spinfection. Medium was switched to non-viral medium three hours post-spinfection. At Day 0, 1 µg/ml dox was added to induce NGN2-expression. At Day 1, transduced hiPSC-NPCs were treated with corresponding antibiotics to the lentiviruses (300 ng/ml puromycin for dCas9-effectors-Puro, 1 mg/ml G-418 for hNGN2-eGFP-neo and 1 mg/ml HygroB for lentiguide-Hygro-mTagBFP2) in order to increase the purity of transduced NPCs. At day 3, NPC medium was switched to neuronal medium (Brainphys (Stemcell Technologies, #05790), 1x N2 (Life Technologies #17502-048), 1x B27-RA (Life Technologies #12587-010), 1 µg/ml Natural Mouse Laminin (Life Technologies), 20 ng/ml BDNF (Peprotech #450-02), 20 ng/ml GDNF (Peptrotech #450-10), 500 µg/ml Dibutyryl cyclic-AMP (Sigma #D0627), 200 nM L-ascorbic acid (Sigma #A0278)) including 1 µg/ml Dox, along with antibiotic withdrawal. 50% of the medium was replaced with fresh neuronal medium (lacking dox once every second day. At day 11, full medium change withdrew residual dox completely. At day 13, NGN2-neurons were treated with 200 nM Ara-C to reduce the proliferation of non-neuronal cells in the culture, followed by half medium change by day 17. At Day 17, Ara-C was completely withdrawn by full medium change, followed by half medium changes until the neurons were fixed or harvested around day 21-24.

#### 2.4 Primary human astrocyte (pHA) co-culture

Commercially available pHAs (Sciencell, #1800; isolated from fetal female brain) were thawed onto a matrigel-coated 100 mm culture dish with commercial astrocyte medium (Sciencell, #1801). While their growing, the astrocytes were fed with fresh astrocyte medium for five days according to the company's manual. Upon their confluence at 90%, astrocytes were detached by TrypLE$^{TM}$ (Thermo Fisher Scientific, #12605010), spun down (200g x 5 mins), resuspended with freezing medium (astrocyte medium supplemented with 10% DMSO) and banked in liquid nitrogen.

At day -2, pHAs were thawed and seeded onto the matrigel-coated 100 mm culture dish and cultured for five days. At day 3, cells were detached, spun down and resuspended with Brainphys basal medium supplemented with Antibiotic-Antimycotic (Anti/Anti; Thermo Fisher Scientific, #15240062) and 2% fetal bovine serum (FBS; Sigma, F4135). Then, cells were split as $1x10^5$ cells / well on a matrigel-coated coverslip. At day 5, pHAs were fed by full medium change with the Brainphys medium (2% FBS + Anti/Anti). At day 7, neurons were split on the pHAs with neuronal medium supplemented with 2% FBS.

Until day 7, NGN2-neurons, when co-cultured with pHAs, were prepared as described above. At day 7, NGN2-neurons were gently detached with Accutase, spun down (1000g × 5 mins) and resuspended in neuronal medium supplemented with 2% FBS. After counting cells with a hemacytometer, NGN2-neurons were seeded on astrocyte culture at different cell densities according to assays (4.5-6×$10^5$ cells/coverslip for presynaptic ICC and 7.5-10×$10^4$ cells/well for MEA). Since day 9, the culture was fed by half medium change along with treatment with 2 µM Ara-C until the day of analysis.

#### 2.5 RNA sequencing data processing of *ATP6V1A* KD and Aβ-treated neurons (New York Genome Center)

RNA Sequencing libraries were prepared using the Kapa Total RNA library prep kit. Paired-end sequencing reads (100bp) were generated on a NovaSeq platform. Raw reads were aligned to hg19 using STAR aligner (v2.5.2a) and gene-level expression were quantified by featureCounts (v1.6.3) based on Ensembl GRCh37.70 annotation model. Genes with over 1 count per million (CPM) in at least 1 sample were retained. After filtering, the raw read counts were normalized by the voom function in limma and differential expression was computed by the moderated t-test implemented in limma.

We examined the GO/pathways impacted by *ATP6V1A* deficit and/or Aβ treatment by employing the Gene Set Enrichment Analysis (GSEA) [19], a weighted enrichment test using all genes devoid of setting a hard threshold to select significant ones since there was a relatively small number of DEGs in KD-V vs WT-V passing the stringent multiple-test correction and no individual gene met the threshold for statistical significance between the Aβ-treated cells and the vehicle-treated cells. In these analyses, the t-test statistics from the differential expression contrast were used to rank genes in the GSEA. Permutations (up to 100,000 times) were used to assess the GSEA enrichment P value.

#### 2.6 *ATP6V1A* KD and Aβ-treatment synergistic effect analysis from the RNA-seq data

The synergistic effect between *ATP6V1A* KD and Aβ-treatment was performed by limma's linear model analysis with formula: Gene expression ~ Sample treatment. The coefficients, standard deviations and correlation matrix were calculated, using *contrasts.fit*, in terms of the comparisons of interest. Empirical Bayes moderation was applied using the *eBayes* function to obtain more precise estimates of gene-wise variability. P-values were adjusted for multiple hypotheses testing using false discovery rate (FDR) estimation, and differentially expressed genes were determined as those with an estimated FDR ≤ 5%, unless stated otherwise. Details about the synergistic effect analysis method were described in [47].

The expected additive effect was modeled through addition of the individual comparisons: (KD-V vs WT-V) + (WT-Aβ vs WT-V). The synergistic effect was modeled by subtraction of the additive effect from the combinatorial perturbation comparison: (KD-Aβ vs WT-V) - (KD-V vs WT-V) - (WT-Aβ vs WT-V). Fitting of this model for differential expression gives genes that show a difference in the differential expression computed for the additive model and that computed for the combinatorial perturbation. However, interpretation of the resulting DEGs depends on several factors, such as the direction of fold change (FC) in all three models. To identify genes of interest, namely those whose magnitude of change is larger in the combinatorial perturbation vs. the additive model, we categorized all genes by the direction of their change in both models and their $\log_2$(FC) in the synergistic model. First, $\log_2$(FC) standard errors (SE) were calculated for all samples. Genes were then grouped into 'positive synergy' if their FC was larger than SE and 'negative synergy' if smaller than -SE. If the corresponding additive model $\log_2$(FC) showed the same or no direction, the gene was classified as "more" differentially expressed in the combinatorial perturbation than predicted. 2925 genes were computed to be in this category (1152 more down, 1773 more up).

GSEA was performed on a curated subset of the MAGMA collection using the limma package camera function, which tests if genes are ranked highly in comparison to other genes in terms of differential expression, while accounting for inter-gene correlation. Due to the small sample size in this study and moderate fold changes in Aβ treatment, changes in gene expression may be small and distributed across many genes. However, similar to previous studies more powerful enrichment analyses in the limma package were used. These evaluate enrichment based on genes that are not necessarily genome-wide significant, and identify sets of genes for which the distribution of t-statistics differs from expectation. Over-representation analysis (ORA) was performed when subsets of DEGs were of interest, such as the synergistic 'more up' and 'more down' genes. The genes of interests were ranked by –log10 (p-value) and enrichment was performed against a background of all expressed genes using the WebGestaltR package.

##### 2.7 Quantitative reverse transcription PCR (qRT-PCR) of *ATP6V1A*

Quantitative reverse transcription PCR (qRT-PCR) was performed as previously described (*Ho et al, 2017*). Specifically, cell cultures were harvested with Trizol and total RNA extraction was carried out following the manufacturer's instructions. Quantitative transcript analysis was performed using a QuantStudio 7 Flex Real-Time PCR System with the Power SYBR Green RNA-to-Ct Real-Time qPCR Kit (all Thermo Fisher Scientific). Total RNA template (25 ng per reaction) was added to the PCR mix, including primers listed below. qPCR conditions were as follows; 48°C for 30 min, 95°C for 10 min followed by 40 cycles (95°C for 15 s, 60°C for 60 s). All qPCR data is collected from at least 3 independent biological replicates of one experiment. Data analyses were performed using GraphPad PRISM 6 software.

| Gene_id | Primer_FWD | Primer_REV | Length |
|---|---|---|---|
| ATP6V1A | GAGATCCTGTACTTCGCACTG | GGGATGTAGATGCTTTGGGTC | 130 |
| β-Actin | TGTCCCCCAACTTGAGATGT | TGTGCACTTTTATTCAACTGGTC | 109 |

#### 2.8 Preparation of cell lysates and western blotting

Cells were rinsed with ice-cold phosphate-buffered saline (PBS), pelleted, and lyzed in RIPA Lysis and Extraction Buffer (Thermo Fisher Scientific, #89900) containing Halt™ Protease and Phosphatase Inhibitor Cocktail (Thermo Fisher Scientific, #78440). Alternatively, MSBB BM36 brain samples were homogenized with similar methods. Samples were sonicated for 1 minute then centrifuged at 13,000× rpm for 10 min. The supernatant was collected, and total protein concentration was determined using Quick Start™ Bradford Protein Assay (Bio-Rad, 5000201) following the manufacturer's instructions.

Western blotting was performed as previously described using antibodies listed in the table below. Images were captured and quantified using the Odyssey® Imaging Systems (LI-COR. Inc.).

| Item Name | Catalog | Species | WB | IF |
|---|---|---|---|---|
| ATP6V1A | ab199326 | Rabbit | 1:1000 | - |
| Homer 1 | 160 003 | Rabbit | 1:1000 | 1:200-500 |
| PSD-95 | clone K28/43 | Mouse IgG2 | 1:500 | 1:1000 |
| Synapsin 1 | 106 011 | Mouse | 1:1000 | 1:500 |
| Synaptophysin 1 | 101 002 | Rabbit | 1:2000 | 1:500 |
| VGLUT 1 | 135 303 | Rabbit | 1:2000 | 1:1000 |
| SOX1 | AF3369 | Goat | - | 1:100 |
| TUJ1 | 801202 | Mouse IgG2a | 1:1000 | 1:1000 |

#### 2.9 Immunofluorescence and microscopy

*NGN2*-neurons (on coverslips) were washed with PBS and fixed with 4% paraformaldehyde (PFA) at pH 7.4 for 10 mins, room temperature. Then, fixative solution was replaced with PBS. After 3 times wash, NGN2-neurons were incubated with blocking solution (0.1% Tween-20, 0.5% bovine serum albumin in PBS) for 1 hour, room temperature. The blocking solution was aspirated and replaced with the same solution with primary antibodies listed above and incubated overnight at 4°C. Neurons were then incubated with secondary antibodies in blocking solution, for one hour at room temperature, followed by PBS-washing 3 times.

20 µL of AquaPolymount mounting solution (Polysciences Inc., #18606-20) per coverslip was placed onto each microscopic slide and the coverslips were gently mounted onto the slides with the neuron side facing down. Mounted coverslips were air-dried for two days at ambient temperature. For synaptic ICC imaging, images were acquired using a confocal microscope (LSM 780, Zeiss) with a 63× objective lens. These puncta analyses were assessed using NIH ImageJ. Total synapsin1 and homer1 puncta number per image were divided by that image's respective MAP2-positive area in order to calculate synapsin1 and homer1 puncta counts normalized to MAP2 levels. Data from 3 independent experiments were analyzed using GraphPad PRISM 6 software.

#### 2.10 β-Amyloid treatment

Human β-amyloid (1-42) peptide was purchased from GenScript (#RP10017). 500 µM β-amyloid was prepared in the vehicle solution of Tris-HCl, pH 7.5 and stored at –20°C. 21-day isogenic pairs of ATP6V1A-manipulated NGN2-neurons were exposed to 5 μM β-amyloid for 24 hours, and then used in qPCR, MEA assays and RNA sequencing.

#### 2.11 Multi-electrode array (MEA)

In order to evaluate electrical activity of NGN2-neurons by MEA, density-matched isogenic NGN2-neuronal populations, co-cultured with pHAs, were prepared as described above. Specifically, at day 3, pHAs were split as 17,000 cells/well in a Matrigel-coated 48W MEA plate (Axion Biosystems, M768-tMEA-48W) and maintained as above. At day 7, NGN2-neurons were detached, spun down and seeded on the pHA culture. Outer space of each well in the plate was filled up with autoclaved/deionized water to minimize the evaporation of marginal wells ("edge effect") during long-term culture. Half volume of neuronal medium (supplemented with 2% FBS) was replaced with fresh medium including 200 nM Ara-C from day 9 until the end of MEA recording. Electrical activity of neurons was daily-recorded during day 14~24. On the recording day, the plate was loaded into the Axion Maestro MEA reader (Axion Biosystems). Recording was performed via AxiS 2.4 for 10 mins. Quantitative analysis of the recording was exported as a Microsoft excel sheet. Data were analyzed using GraphPad PRISM 6 software.

#### 2.12 Electrophysiology

For whole-cell patch-clamp recordings, 1.0-1.5 x 104 human astrocytes were first seeded onto Matrigel-coated 12-mm glass coverslips in 24-well plates, and then seeded with 1.0 x 105 neurons after ~5 days. Neurons were recorded at 4-5 weeks following dox-induction, with media exchange every 3-4 days. Cells were visualized on a Nikon inverted microscope equipped with fluorescence and Hoffman optics. Neurons were recorded with an Axopatch 200B amplifier (Molecular Devices), digitized using a Digidata 1320a (Molecular Devices) and filtered between

1-10 kHz, using Clampex 10 software (Molecular Devices). Series resistance compensation was applied (70-100%). Patch pipettes were pulled from borosilicate glass electrodes (Warner Instruments) to a final tip resistance of 3-5M Ω using a vertical gravity puller (Narishige). Neurons were bathed in artificial cerebral spinal fluid (ACSF) containing (in mM): NaCl, 119, CaCl22; KCl, 2.5; MgCl2, 1.3; d-glucose, 11; NaHCO3, 26.2; NaPO4, 1, at a pH of 7.4. The internal patch solution contained (in mM): K-d-gluconate, 140; NaCl, 4; MgCl2, 2; EGTA, 1.1; HEPES, 5; Na2ATP, 2; sodium creatine phosphate, 5; Na3GTP, 0.6, at a pH of 7.4. Osmolarity was 290-295 mOsm. All chemicals were purchased from Sigma-Aldrich Co. (St. Louis, MO). Neurons were chosen at random using DIC or with BFP+ expression. Current-clamp recordings were used for measuring evoked (current injected to hyperpolarize to approx. -80 mV) activity. Spikelets were defined as small, outward spikes with a peak amplitude of less than 25 mV and occurred at voltages positive to threshold (~ -30 mV). In voltage-clamp recordings, voltage steps were applied from -80 mV to +50 mV (10 mV increments) to elicit voltage-gated ionic currents All recordings were made at room temperature (~22 C). Difference between sodium current densities at 0 mV were tested for statistical significance ($P < .05$) using a student's t-test between control (n=18) and ATP6V1A KD neurons (n=17), pooling over two experimental replicates. Voltages are corrected for a junction potential of ~ -15 mV. Values are reported as mean ± SEM.

#### 3 *Drosophila* genetics analysis

Flies were maintained in standard cornmeal media at 25 ºC. Transgenic fly lines carrying *UAS-Aβ42* were previously described [48]. The *elav*-GAL4 (#458), UAS-*mcherry* RNAi (#35785), UAS-*Vha68-1* RNAi (#50726 and #42888) were obtained from the Bloomington Stock Center.

Genotypes and ages of all flies used in this study are provided in Figure legends. Experiments were performed using age-matched male flies.

#### 3.1 ATP6V1A orthologs in fly

According to the DIOPT (DRSC Integrative Ortholog Prediction Tool), *Drosophila* Vacuolar $H^+$ ATPase 68kD subunit 1 (*Vha68-1*, *CG12403*) and *Vha68-2* (*CG3762*) are the best orthologs of human *ATP6V1A* proteins (DIOPT score 14 for both genes). Vha68-1 and ATP6V1A exhibit 83% identity and 91% similarity in primary amino acid sequence, and have similar size (614 and 617 amino acids, respectively), while Vha68-2 and ATP6V1A exhibit 83% identity and 92% similarity in primary amino acid sequence with similar size (614 and 617 amino acids, respectively).

#### 3.2 RNA extraction and quantitative real time PCR analysis

More than 25 flies for each genotype were collected and frozen. Heads were mechanically isolated, and total RNA was extracted using TRIzol Reagent (Thermo Fisher Scientific) according to the manufacturer's protocol with an additional centrifugation step (16,000 x *g* for 10 min) to remove cuticle membranes prior to the addition of chloroform. Total RNA was reverse-transcribed using PrimeScript RT-PCR kit (TaKaRa Bio), and qRT-PCR was performed using Thunderbird SYBR qPCR Mix (Toyobo) on a CFX96 real time PCR detection system (Bio-Rad Laboratories). The average threshold cycle value was calculated from at least three replicates per sample. Expression of genes of interest was standardized relative to GAPDH1. Primer sequences used in this study are provided below.

| Fly | Human | Forward primer sequence (5' to 3') | Reverse primer sequence (5' to 3') |
|---|---|---|---|
| *Gapdh* | *GAPDH1* | GACGAAATCAAGGCTAAGGTCG | AATGGGTGTCGCTGAAGAAGTC |

| | | | |
|---|---|---|---|
| *Vha68-1** | *ATP6V1A* | ACTACGCACCAAGGTCAAGG | CTTTGCCACTTCCAGGGTCA |
| *Vha68-1*** | *ATP6V1A* | ACCTCTTTCCGTGGAACTTGG | GCAGTTGTGTTGACACCTTTGG |
| *Vha68-2* | *ATP6V1A* | CAAATATGGACGTGTCTTCGC | CCGGATCTCCGACAGTTACG |
| *Eaat1* | *SLC1A2* | TGCTCTGTTCATCGCCCAAT | CGACGGCTATGATGAGGGAC |
| *VGlut* | *SLC17A6-8* | TTCATCGCCTCCAAGTTCCC | GCTGGATAGGTAACGCCCTC |
| *Atpα* | *ATP1A1-3* | ACATGGTGCCAGCCATTTCA | AAGCCGTTCTCAGCCATGAT |
| *Lcch3* | *GABRB1-3* | CCGAGACGTGTTCAACGACA | GGCTATGTCCGGTGCCATAA |
| *Grd* | *GABRA1-6* | TTTGGCTACACAACGTCGGA | GGTCGTGGTGGATCCTTGTT |
| *CG8916* | *GABRA6* | TTGAGTCCAAGAGCGGTGTC | CGTTTGGGTGGTTCTCTCCA |
| *Rdl* | *GABRP* | TGGCTCAATCGCAATGCAAC | GACCGTGGCGTATTCCAGTA |
| *CG12344* | *GLRA2* | CGAGAGCTTCTCGTCGAACA | GAAGTATACGGTGAGGCGGG |
| *GABA-B-R1* | *GABBR1* | CTCAGGGCGATCGTATTGCT | GGTGCGTAGAACATGGGTGA |
| *GABA-B-R2* | *GABBR2* | TGGCGGGTGCATTCGATATT | TCTCGTGATGCAAGGGTTCC |
| *GABA-B-R3* | *GABBR2* | AATTCGCACAGCAATCTGCC | ACAGCTCAAAGAGTCCGAGC |
| *Sh* | *KCNA1-7,10* | TGTCAGGTTCCTCGCATGTC | CTGACTGGCGCTTTTGGAAG |
| *Shab* | *KCNB1,2* | CGTGCTCGCGTTTAGTGATG | TTCTGGTACTCGGCGCATTT |
| *SK* | *KCNN1-3* | GGTTATCGAAAACGAACTGAGCA | CTTCCAAAGCATGGTAAGCTAC |
| *para* | *KCNB1,2* | ACGAGGATGAAGGTCCACAAC | ACGACGTATCGGATTGAATGG |
| *Nmdar2* | *GRIN2A-D* | GGCATCCCGGTTATCTCGTG | AGAACTGGTGCCACTTGTAGC |

#### 3.3 Climbing assay

Approximately 25 male flies were placed in an empty plastic vial. The vial was then gently tapped to knock all of the flies to the bottom. The numbers of flies in the top, middle, or bottom thirds of the vial were scored after 10 seconds. The percentages of flies that stayed at the bottom were subjected to statistical analyses.

#### 3.4 Histological analysis

Heads of male flies were fixed in 4% paraformaldehyde for 24 h at 4 ºC and embedded in paraffin. Serial sections (6μm thickness) through the entire heads were prepared, stained with hematoxylin and eosin (Sigma-Aldrich), and examined by bright-field microscopy. Images of the sections were captured with AxioCam 105 color (Carl Zeiss). For quantification, we focused on analyzing sections covering the central neuropil regions. We selected a section with the most severe neurodegeneration in each individual fly and the area of vacuoles was measured using Image J (NIH).

#### 3.5 Western blotting

Ten fly heads for each genotype were homogenized in Tris-Glycine SDS sample buffer, and the same amount of the lysate was loaded to 18% Tris-Glycine gels and transferred to nitrocellulose membrane. The membranes were boiled in PBS for 3 min, blocked with 5% nonfat dry milk, blotted with the anti-Aβ 6E10 antibody (Signet, Covance), incubated with appropriate secondary antibody and developed using ECL Western Blotting Detection Reagents (GE Healthcare Life Sciences). The membranes were also probed with anti-tubulin (Sigma-Aldrich) as the loading control in each experiment. Imaging was performed with ImageQuant LAS 4000 (GE Healthcare Life Sciences), and the signal intensity was quantified using Image J (NIH).

### DATA AND SOFTWARE AVAILABILITY

#### Data Resources

All the human postmortem brain sequencing data are available from the Synapse (https://www.synapse.org/) through accession numbers syn17008933 and syn17008927. RNA-seq data of the hiPSC-derived neurons are available from the gene expression omnibus (GEO) through accession number GSE128367 with secure token khuviiuyffkztyd. All other relevant data are available from the corresponding author upon reasonable request.

### Supplementary figures

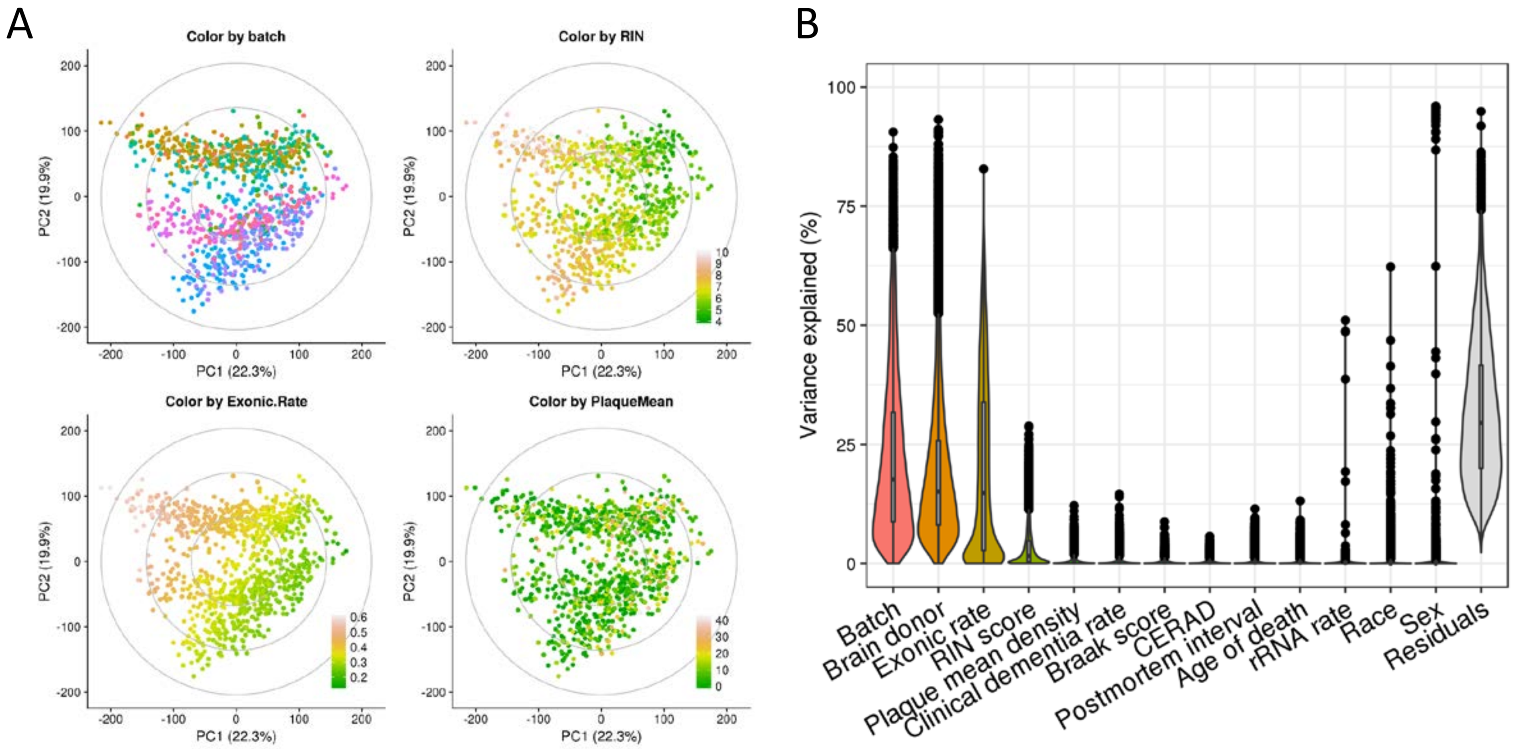


**Fig. S1. RNA-seq data correction for confounding factors. A**, Principal component (PC) plots showing the association between biological/technical variables and top PCs. **B**, Distribution of variance explained by the biological/technical variables according to variancePartition analysis. From left to right, variables were sorted in descending order by median fraction of variance explained.

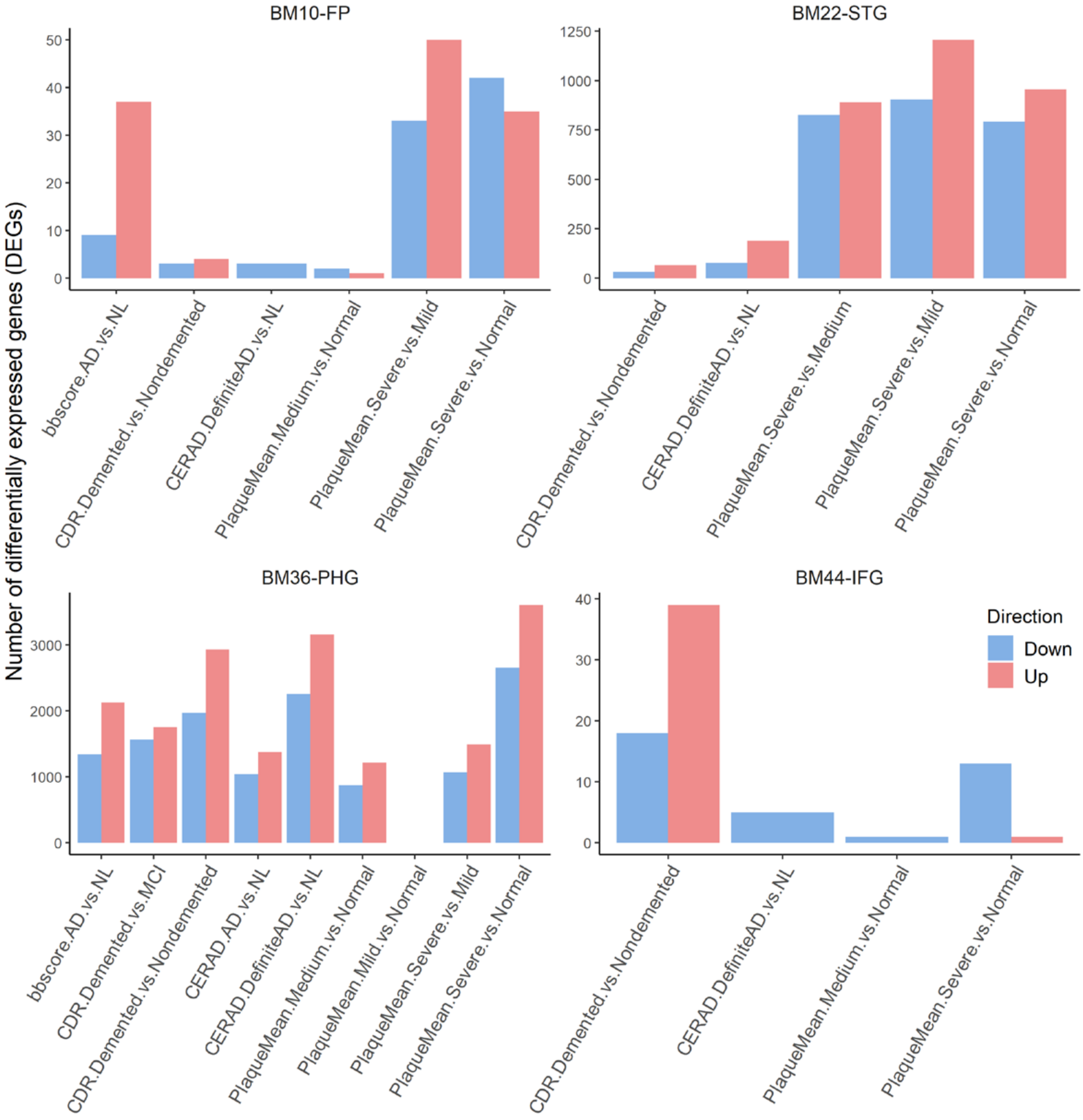


**Fig. S2. Summary of differential expression signatures.** Number of DEGs identified in each contrast for each region regarding 4 different cognitive/neuropathological traits. X-axis denotes the trait-specific contrast. Down, down-regulated DEGs; Up, up-regulated DEGs.

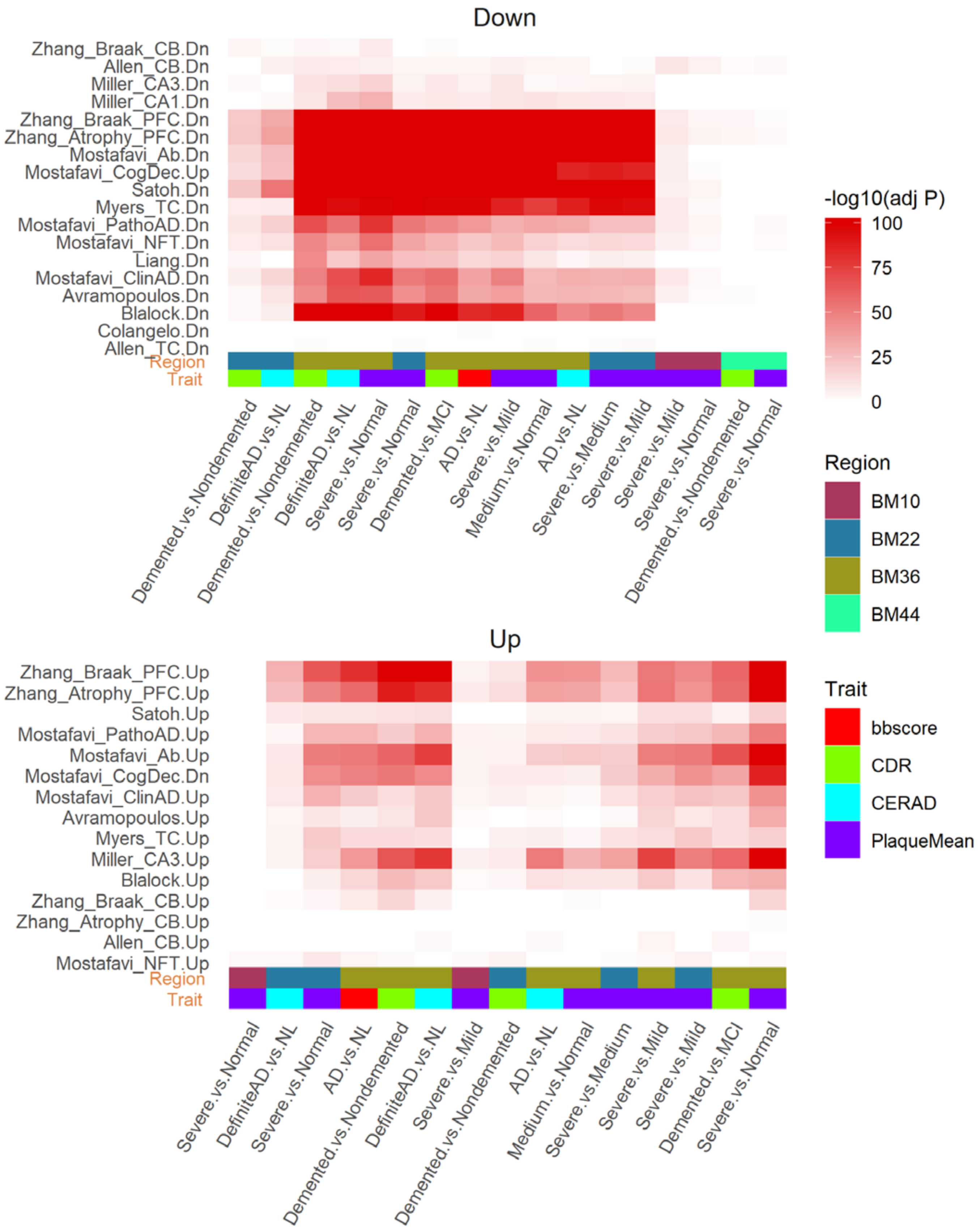
Down
Zhang_Braak_CB.Dn
Allen_CB.Dn
Miller_CA3.Dn
Miller_CA1.Dn
Zhang_Braak_PFC.Dn
Zhang_Atrophy_PFC.Dn
Mostafavi_Ab.Dn
Mostafavi_CogDec.Up
Satoh.Dn
Myers_TC.Dn
Mostafavi_PathoAD.Dn
Mostafavi_NFT.Dn
Liang.Dn
Mostafavi_ClinAD.Dn
Avramopoulos.Dn
Blalock.Dn
Colangelo.Dn
Allen_TC.Dn
Region
Trait
Demented.vs.Nondemented
DefiniteAD.vs.NL
Demented.vs.Nondemented
DefiniteAD.vs.NL
Severe.vs.Normal
Severe.vs.Normal
Demented.vs.MCI
AD.vs.NL
Severe.vs.Mild
Medium.vs.Normal
AD.vs.NL
Severe.vs.Medium
Severe.vs.Mild
Severe.vs.Mild
Severe.vs.Normal
Demented.vs.Nondemented
Severe.vs.Normal
Up
Zhang_Braak_PFC.Up
Zhang_Atrophy_PFC.Up
Satoh.Up
Mostafavi_PathoAD.Up
Mostafavi_Ab.Up
Mostafavi_CogDec.Dn
Mostafavi_ClinAD.Up
Avramopoulos.Up
Myers_TC.Up
Miller_CA3.Up
Blalock.Up
Zhang_Braak_CB.Up
Zhang_Atrophy_CB.Up
Allen_CB.Up
Mostafavi_NFT.Up
Region
Trait
Severe.vs.Normal
DefiniteAD.vs.NL
Severe.vs.Normal
AD.vs.NL
Demented.vs.Nondemented
DefiniteAD.vs.NL
Severe.vs.Mild
Demented.vs.Nondemented
AD.vs.NL
Medium.vs.Normal
Severe.vs.Medium
Severe.vs.Mild
Severe.vs.Mild
Demented.vs.MCI
Severe.vs.Normal
-log10(adj P)
100
75
50
25
0
Region
BM10
BM22
BM36
BM44
Trait
bbscore
CDR
CERAD
PlaqueMean

**Fig. S3**

**Fig. S3. Preservation of differential expression signatures in published bulk-tissue transcriptomic analysis datasets.** Public AD signatures enriched in the present down-regulated (top) or up-regulated (bottom) DEGs. In each panel, columns denote the present region-specific DEGs stratified by region, trait and contrast. The rows denote public AD signatures written in a format of “signature.direction”, where “direction” is either Dn (down-regulated) or Up (up-regulated), and “signature” includes: 1) Allen_CB, signature detected in the cerebellum region from Allen et al 2018, 2) Allen_TC, signature detected in the temporal cortex region from Allen et al 2018, 3) Avramopoulos, signature detected in the temporal lobe from Avramopoulos et al 2011, 4) Blalock, signature detected in the hippocampus from Blalock et al 2004, 5) Colangelo, signature detected in the hippocampus CA1 region from Colangelo et al 2002, 6) Liang, signature detected in multiple cortex areas from Liang et al 2008, 7) Miller_CA1, signature detected in the hippocampus CA1 region from Miller et al 2013, 8) Miller_CA3, signature detected in the hippocampus CA3 region from Miller et al 2013, 9) Mostafavi_Ab, signature correlated with B amyloid in the prefrontal cortex (PFC) region from Mostafavi et al 2018, 10) Mostafavi_ClinAD, signature correlated with clinical diagnostic of AD in the PFC from Mostafavi et al 2018, 11) Mostafavi_CogDec, signature correlated with cognitive decline in the PFC from Mostafavi et al 2018, 12) Mostafavi_PathoAD, signature correlated with AD pathology in the PFC from Mostafavi et al 2018, 13) Satoh; signature detected in the frontal cortex region from Satoh et al 2014, 14) Myers_TC, signature computed by comparing gene expression between AD and control with limma from the temporal cortex region from Webster et al 2009, 15) Zhang_Atrophy_CB, signature correlated with atrophy in the cerebellum region from Zhang et al 2013, 16) Zhang_Atrophy_PFC, signature correlated with atrophy in the PFC from Zhang et al 2013, 17) Zhang_Braak_CB, signature correlated with Braak staging in the cerebellum region from Zhang et al 2013, 18) Zhang_Braak_PFC, signature correlated with Braak staging in the PFC from Zhang et al 2013. The references of the public AD signatures are listed in supplementary text.

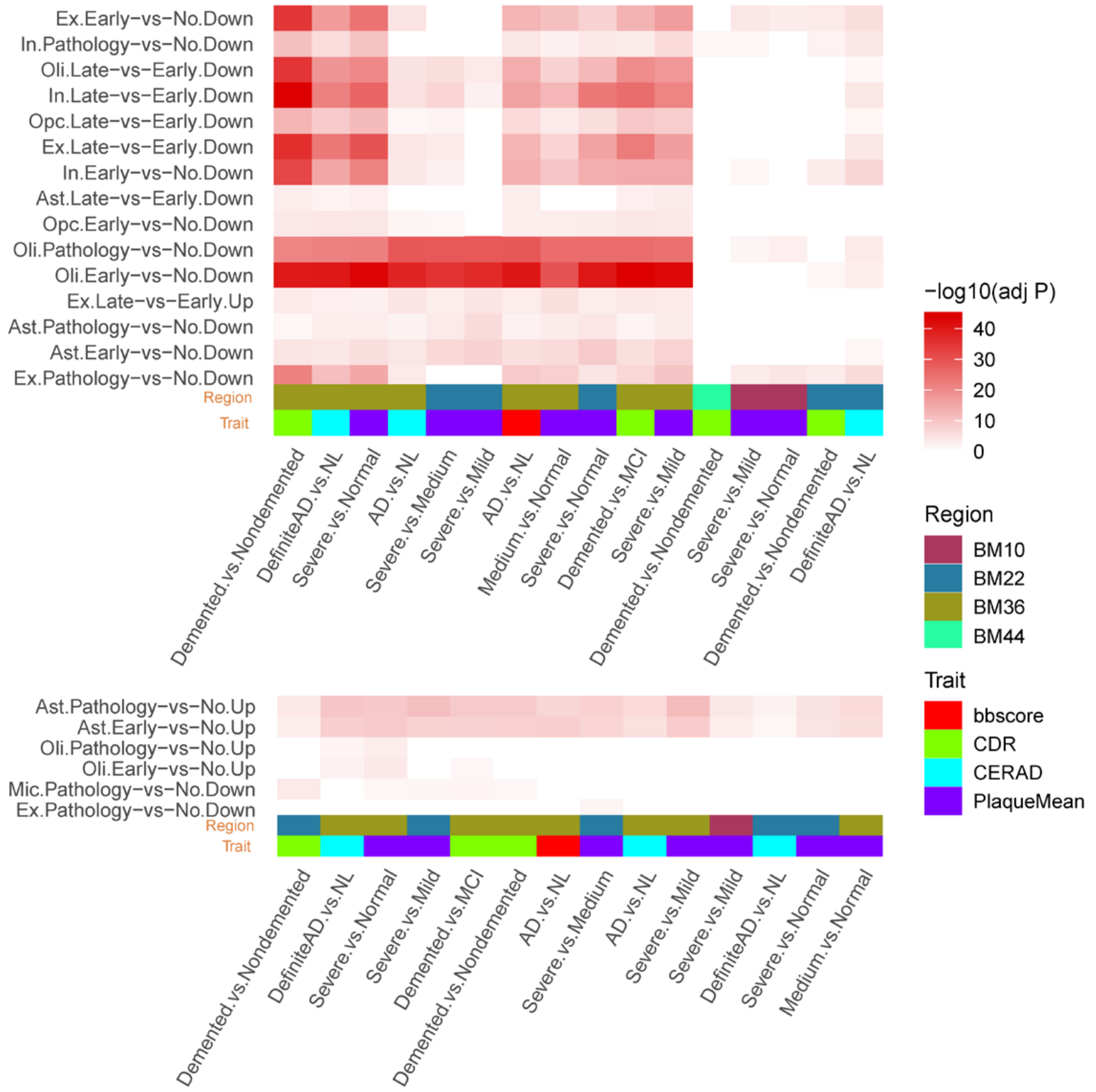


**Fig. S4. Preservation of differential expression signatures in a public single-nucleus RNA-seq (snRNA-seq) analysis of AD and control brains.** The present down-regulated (top) or up-regulated (bottom) DEGs were enriched in cell type-specific AD signatures detected by snRNA-seq (Mathys et al 2019). Columns denote the present DEG signatures identified from 4 brain regions regarding 4 different cognitive/neuropathological traits and rows denote cell type-specific DEGs. The snRNA-seq cell type-specific DEGs are denoted in a format of "cell.contrast.direction", where cell is either Ex (excitatory neurons), In (inhibitory neurons), Oli (oligodendrocytes), Opc (oligodendrocyte progenitor cells), Ast (astrocytes), or Mic (microglia), contrast is either Early-vs-No (early pathology versus no pathology), Late-vs-Early (late pathology versus early pathology), or Pathology-vs-No (early and late pathology combined versus no pathology), and direction is either Down (down-regulation) or Up (up-regulation).

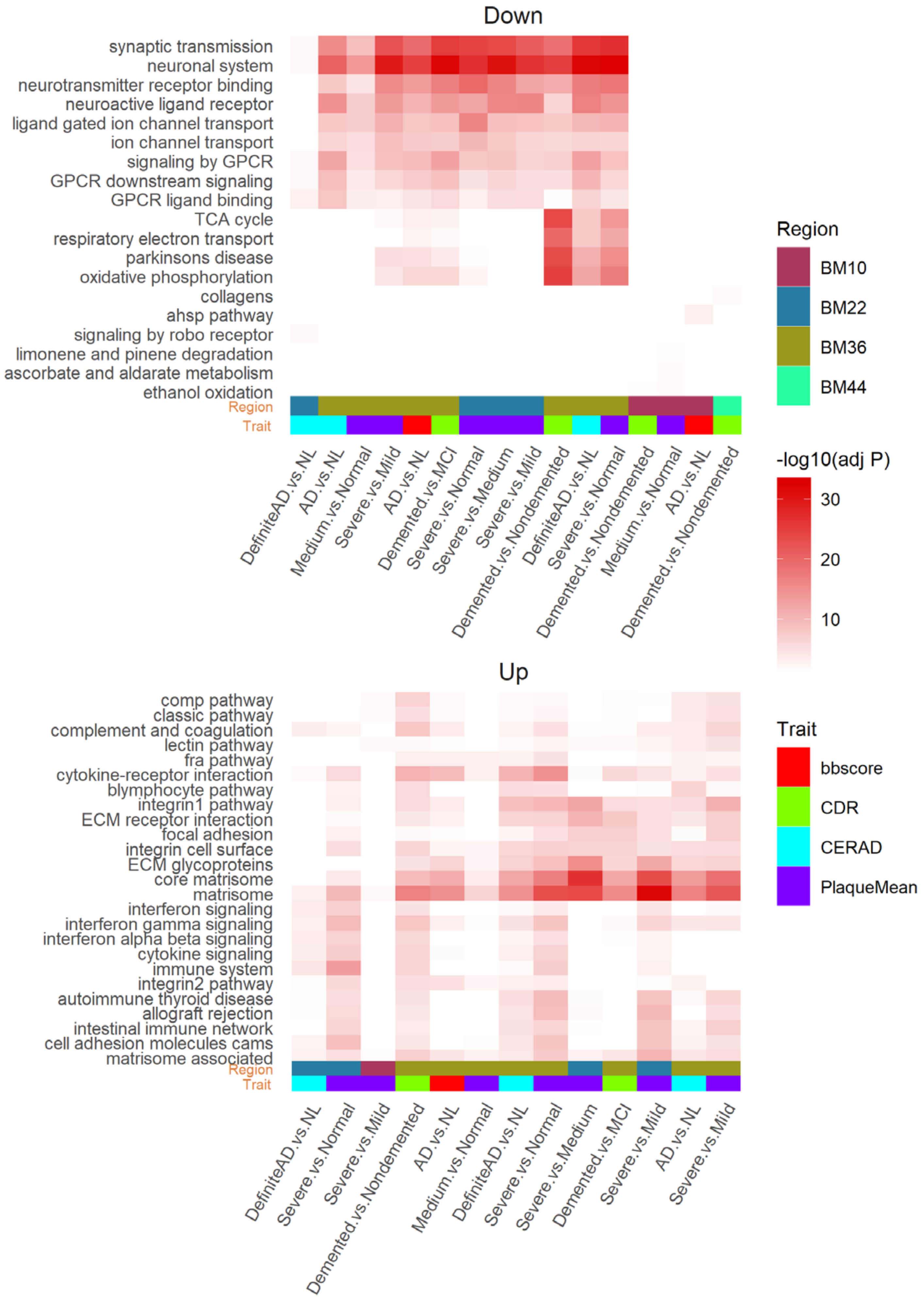


**Fig. S5. Summary of gene ontology (GO)/pathways enriched in differential expression signatures.** Top pathways enriched in the present down-regulated (top) or up-regulated (bottom) DEGs. Columns denote different set of DEGs from 4 brain regions regarding 4 different cognitive/neuropathological traits and rows denote GO/pathways. Down, down-regulated DEGs; Up, up-regulated DEGs.

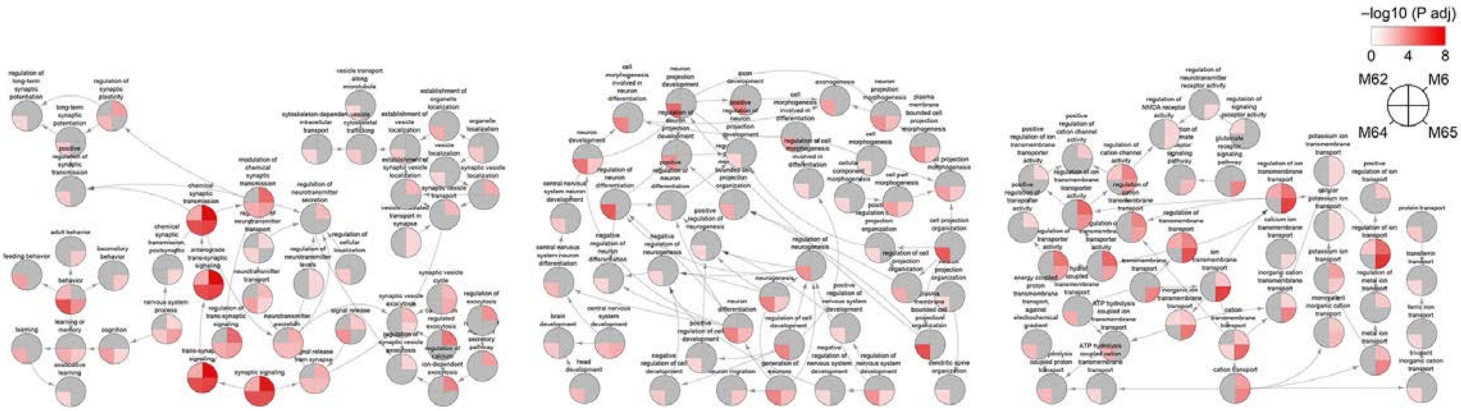


**Fig. S6. GO biological process (BP) hierarchy enrichment reveals distinct functional roles that the top ranked neuronal/synaptic modules may play.** Each node denotes a GO/BP term, with a pie-chart displaying the $-\log_{10}$(adjusted P value) of the FET enrichment for the 4 top ranked neuronal/synaptic modules (i.e. M6, M62, M64 and M65). Arrows denote the direction from a parent term to a child term. The GO hierarchy was extracted from the R/Bioconductor package GO.db and the GO/BP annotation gene sets were obtained from the R/Bioconductor package org.Hs.eg.db. From left to right, the three subplots are grouping terms in relation to synaptic function, neuronal development and transportation, respectively.

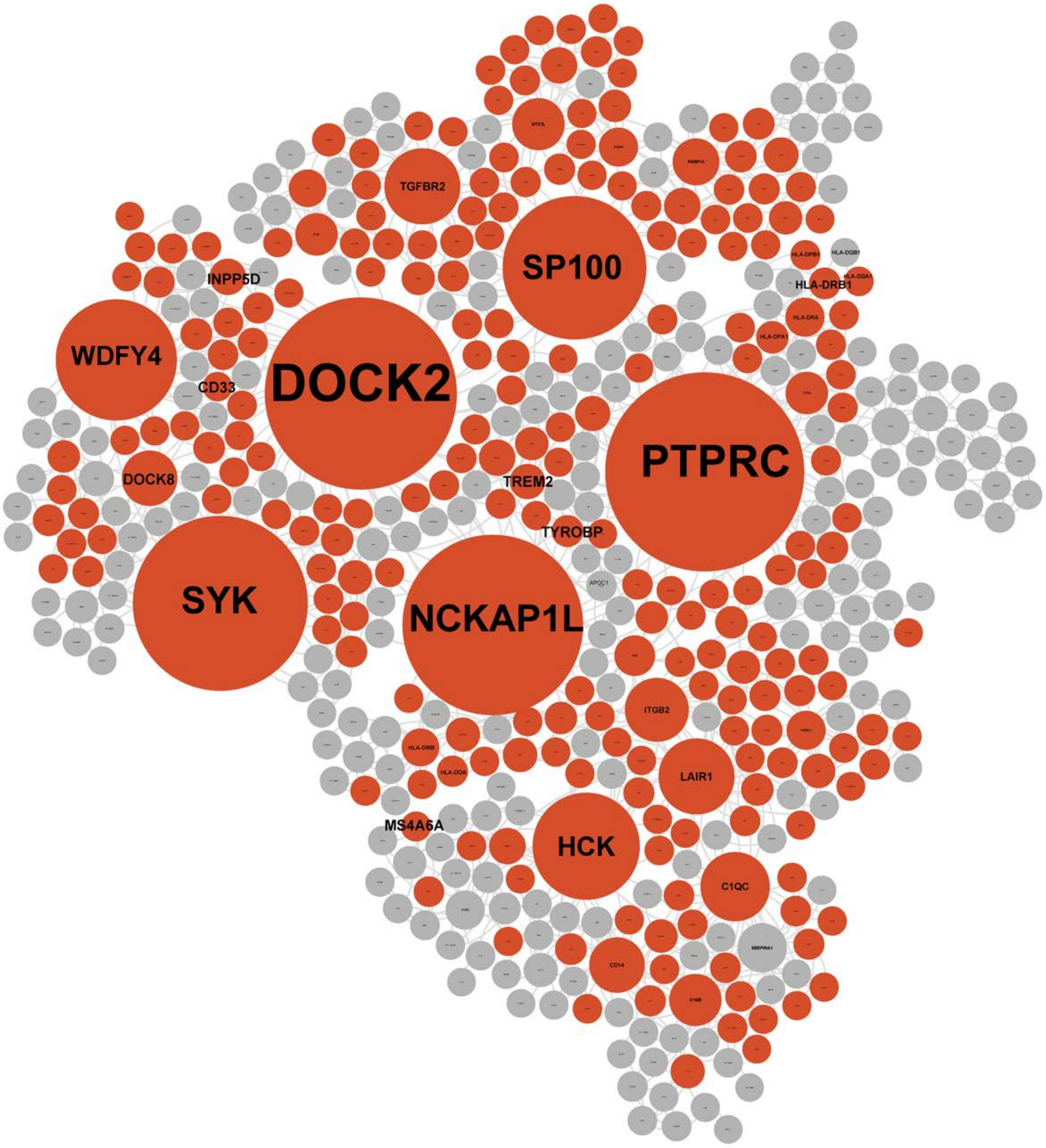


**Fig. S7. Topological structure of immune response module M153.** Node color denotes whether the gene is up-regulated (red), down-regulated (blue), or no change (grey) in demented brains. Node size is proportional to the connectivity within module.

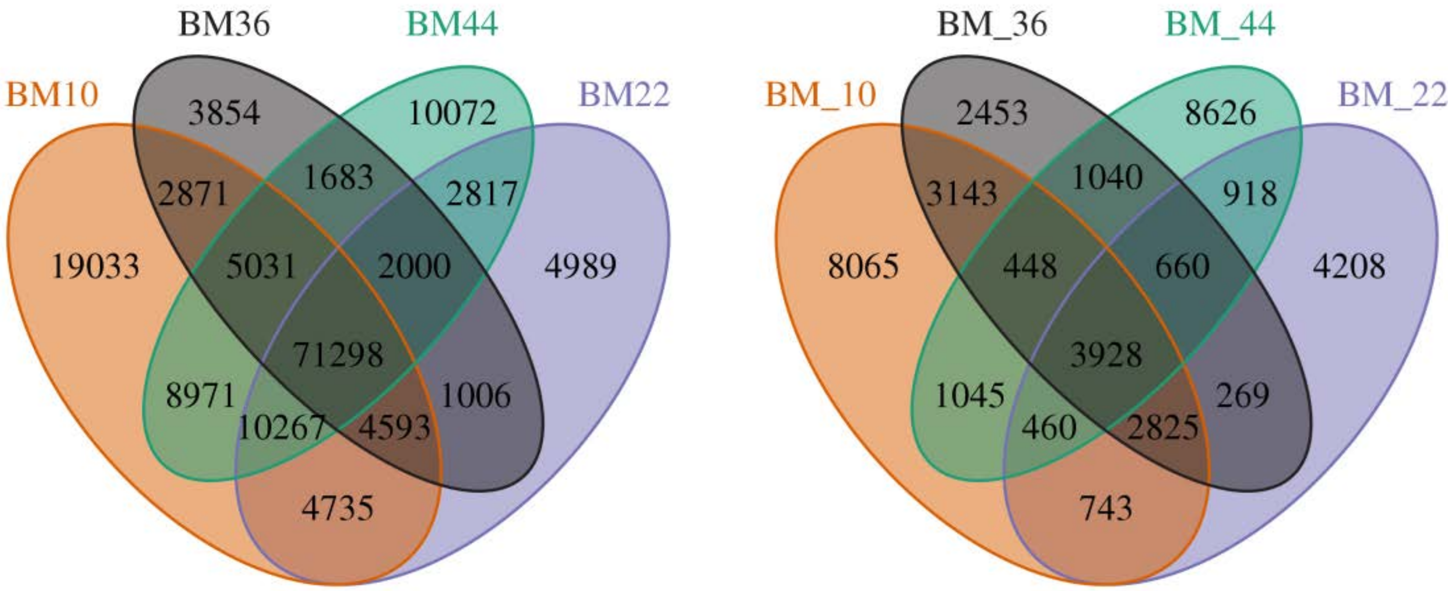


**Fig. S8. eQTLs are shared among different regions.** Venn-diagram showing the overlap among *cis*-eQTLs (left) and *trans*-eQTLs (right).

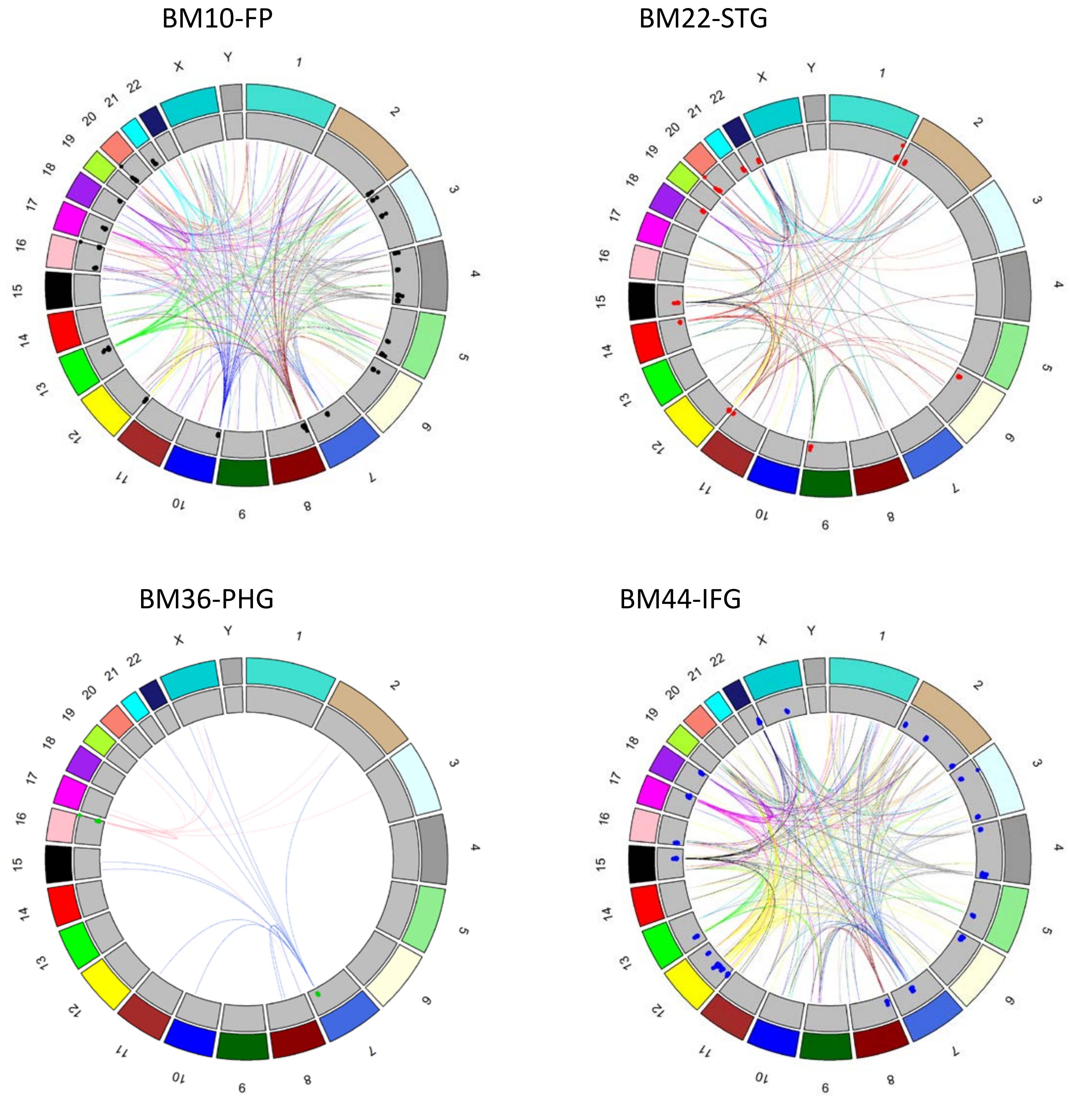


**Fig. S9. *Trans*-eQTL hotspots identified in the 4 brain regions.** The outermost first track shows the chromosome id. The second track denotes the color coding of the chromosomes. The dots in the third track denote the –log10(P value) of the *trans*-eQTL associations for the hotspots. Links in the middle connect the hotspots to the associated *trans*-eGenes. Links are colored by the chromosomal origination of the *trans*-eQTL hotspots.

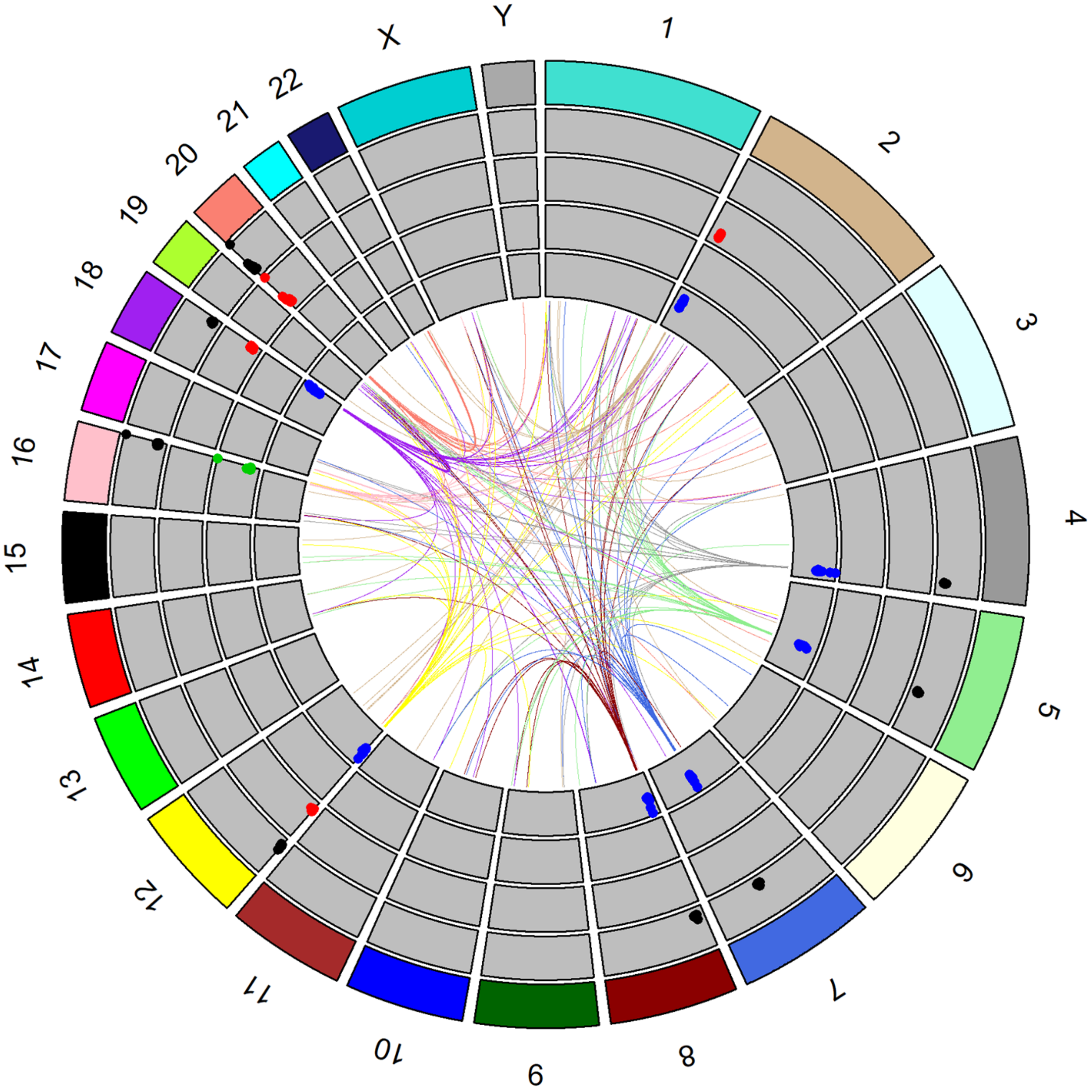


**Fig. S10. *Trans*-eQTL hotspots shared by at least two brain regions.** The outermost first track shows the chromosome id. The second track denotes the color coding of the chromosomes. The dots in the third to sixth tracks denote the –log10(P value) of the *trans*-eQTL associations for the hotspots in brain region BM10-FP, BM22-STG, BM36-PHG and BM44-IFG, respectively. Links in the middle connect the hotspots to the associated *trans*-eGenes. Links are colored by the chromosomal origination of the *trans*-eQTL hotspots.

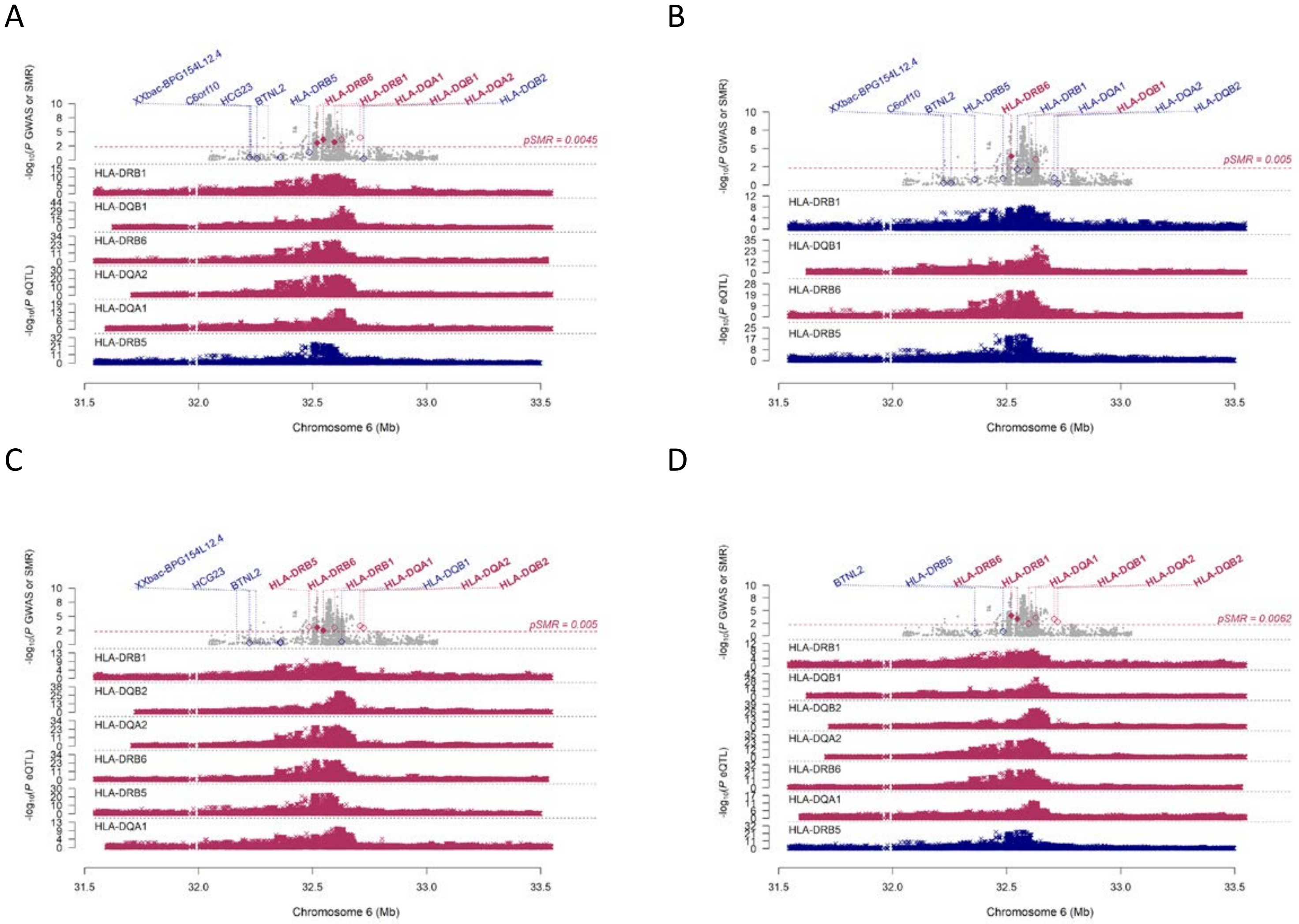


**Fig. S11. Summary data-based Mendelian randomization (SMR) analysis at the *HLA-DRB1/HLA-DRB5* locus. A-D** show the SMR analysis results integrating IGAP AD GWAS with eQTLs derived from brain regions BM10, BM22, BM36 and BM44, accordingly. For the top panel in each plot, dots represent the P values for SNPs from the IGAP AD GWAS analysis and diamonds represent the P values for genes from the SMR test. Filled diamonds highlight the genes surpassing the HEIDI test ($P \geq 0.05$). The genes with cis-eQTLs are listed on the top. Genes surpassing the SMR test were highlighted in red. Dashed line shows the region-specific Bonferroni corrected P value significance threshold of the SMR test controlling for the number of genes examined. Bottom plot shows the eQTL P values of SNPs from the present data.

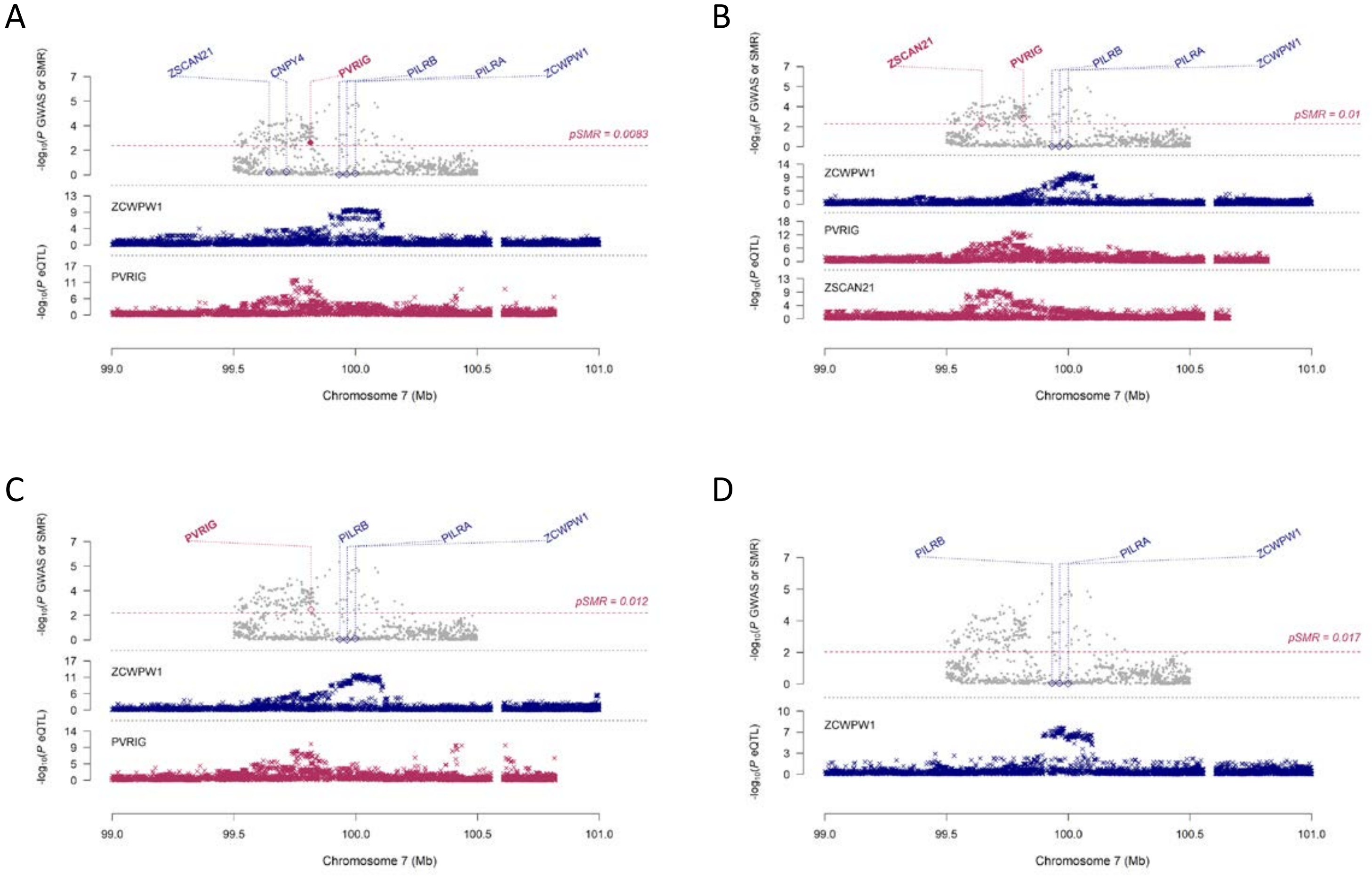


**Fig. S12. SMR analysis at the *ZCWPW1* locus. A-D** show the results from BM10, BM22, BM36 and BM44, accordingly. Figure legend is the same as in **Fig S11**.

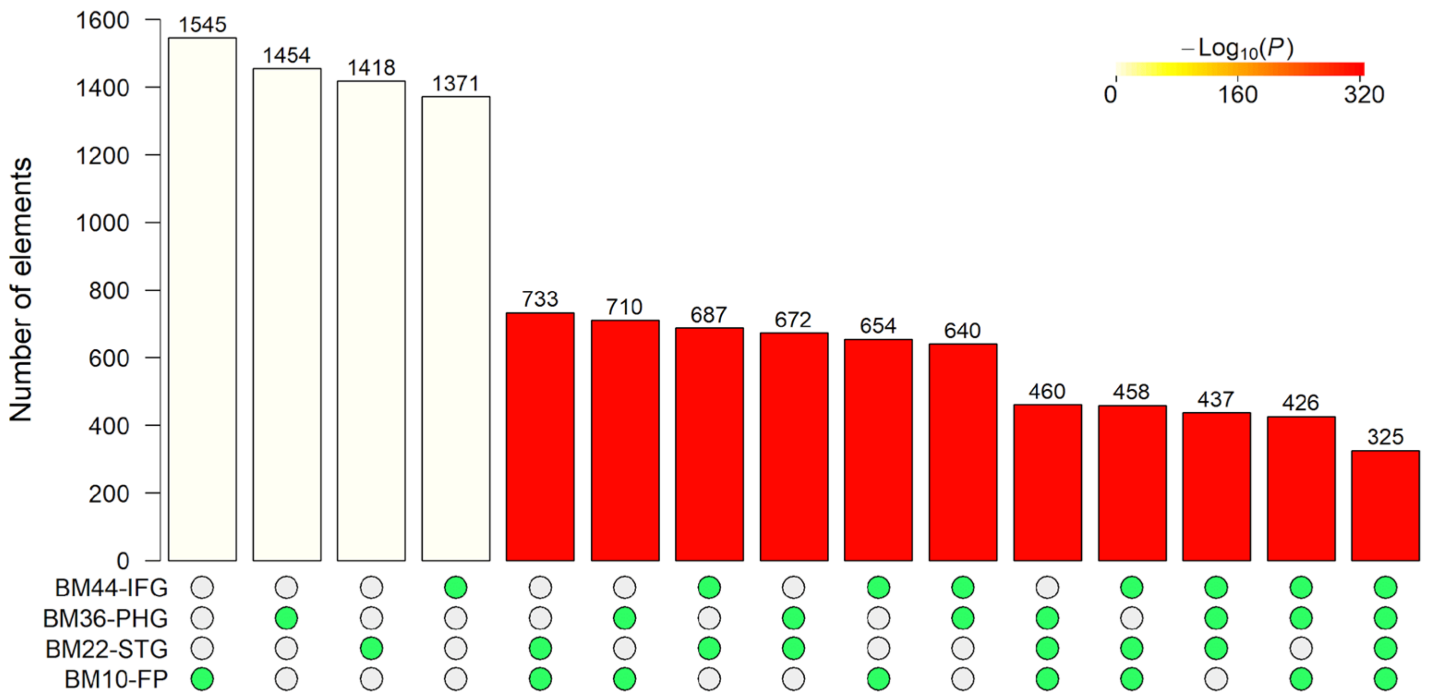


**Fig. S13. Bayesian network key drivers are strongly shared among brain region-specific networks.** The bar height denotes the number of key drivers overlapping in a given comparison as specified by the green circles underneath. The overlap size is also shown above the bar. The color denotes the P value significance of the overlap size.

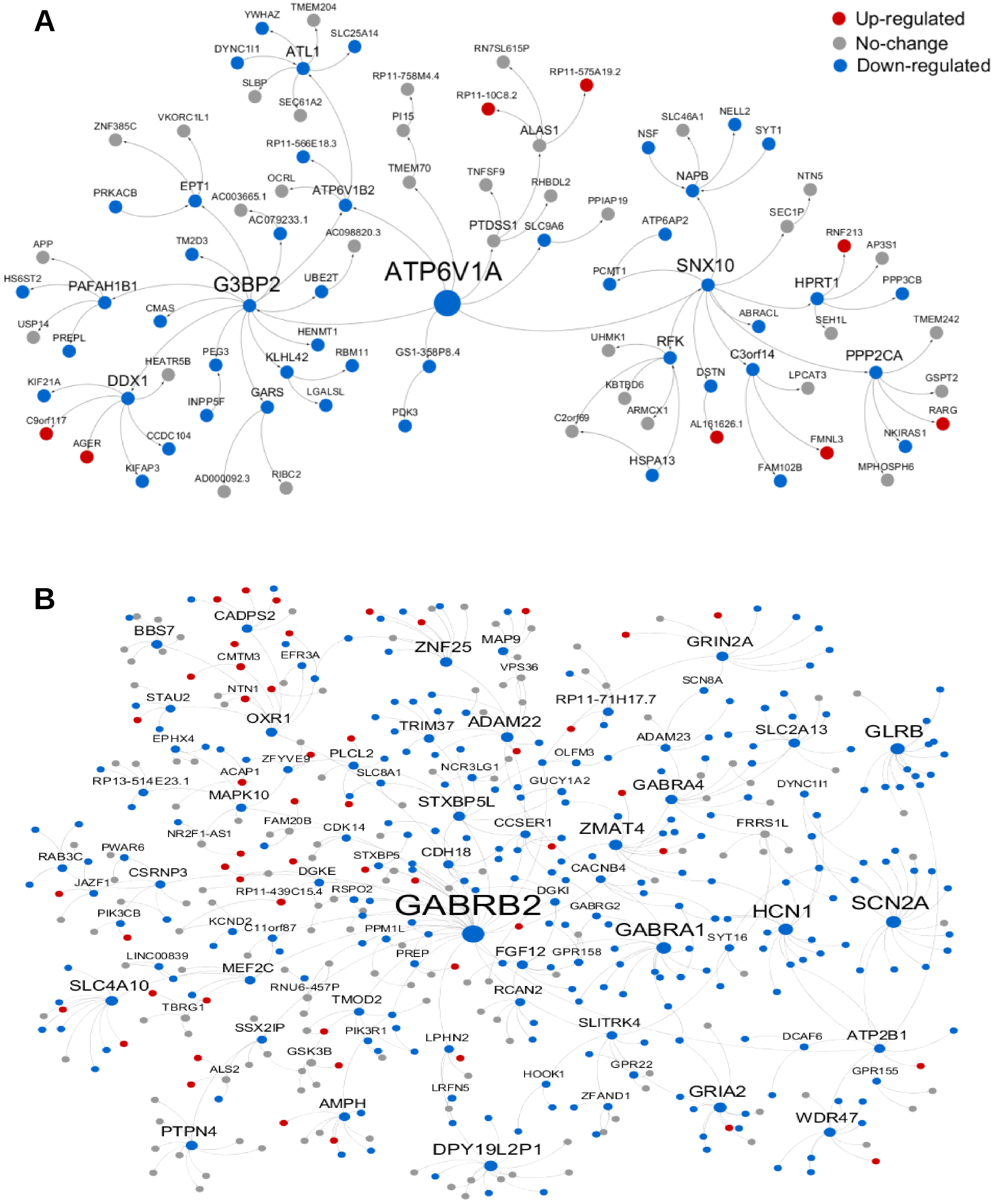


**Fig. S14. Network neighborhood of key driver genes *ATP6V1A* and *GABRB2* on the BM36-PHG BN.** Down-regulated genes in demented brains are enriched in network neighboring (A) *ATP6V1A* (FE = 5.4 and FDR = 3.7E-19) and (B) *GABRB2* (FE = 5.8, FDR = 1.1E-106). Node color denotes the direction of gene expression change in in the BM36-PHG region of patients with dementia (CDR ≥ 1).

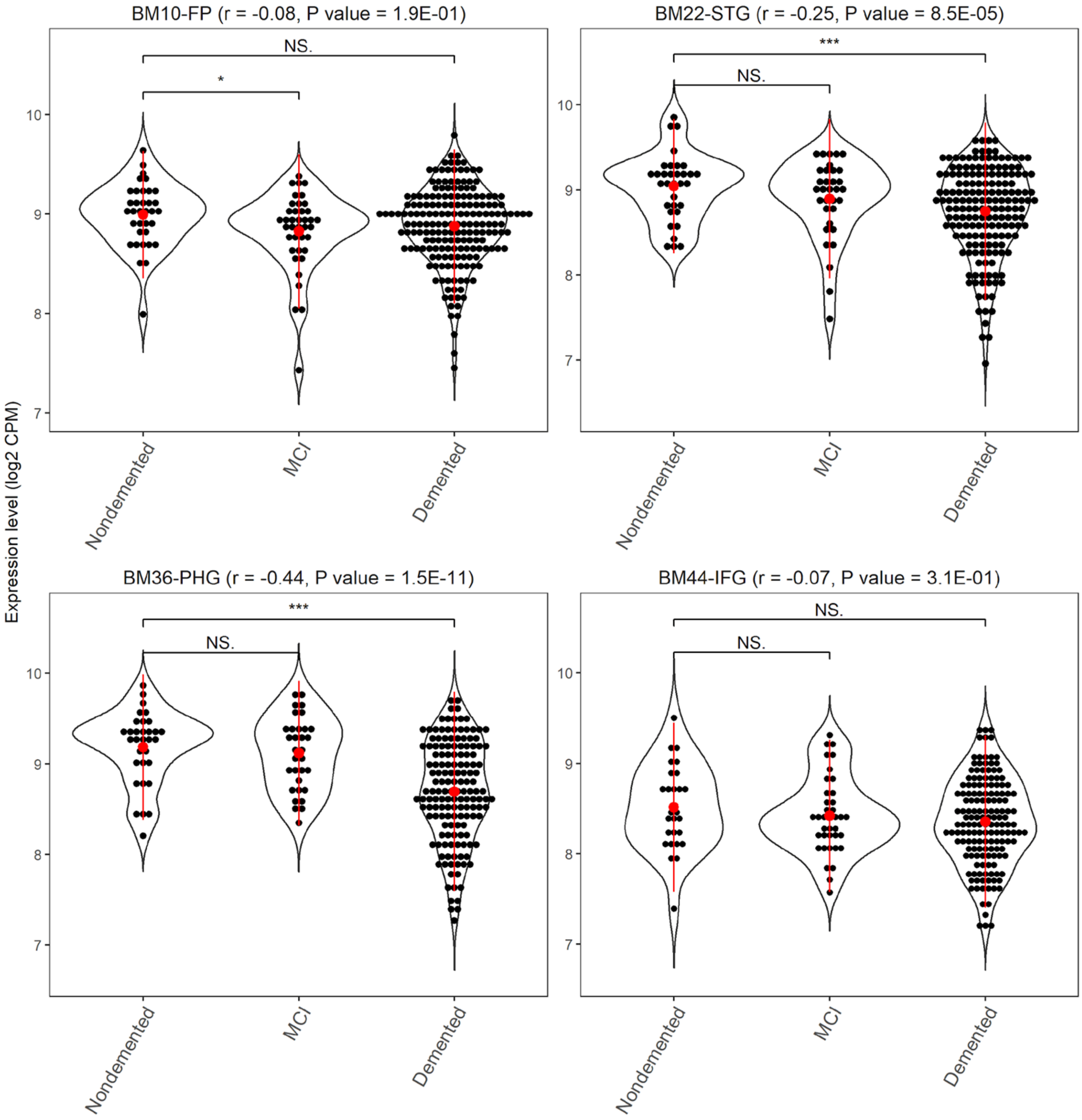


**Fig. S15. *ATP6V1A* is down-regulated in demented patients in multiple brain regions.** Significance bar represents *t*-test P value. *p < 0.05; ***p < 0.001, N.S., no significance. Correlation coefficient (r) and P value of the Spearman correlation between *ATP6V1A* expression and clinical trait CDR are also shown in the top of each sub-plot.

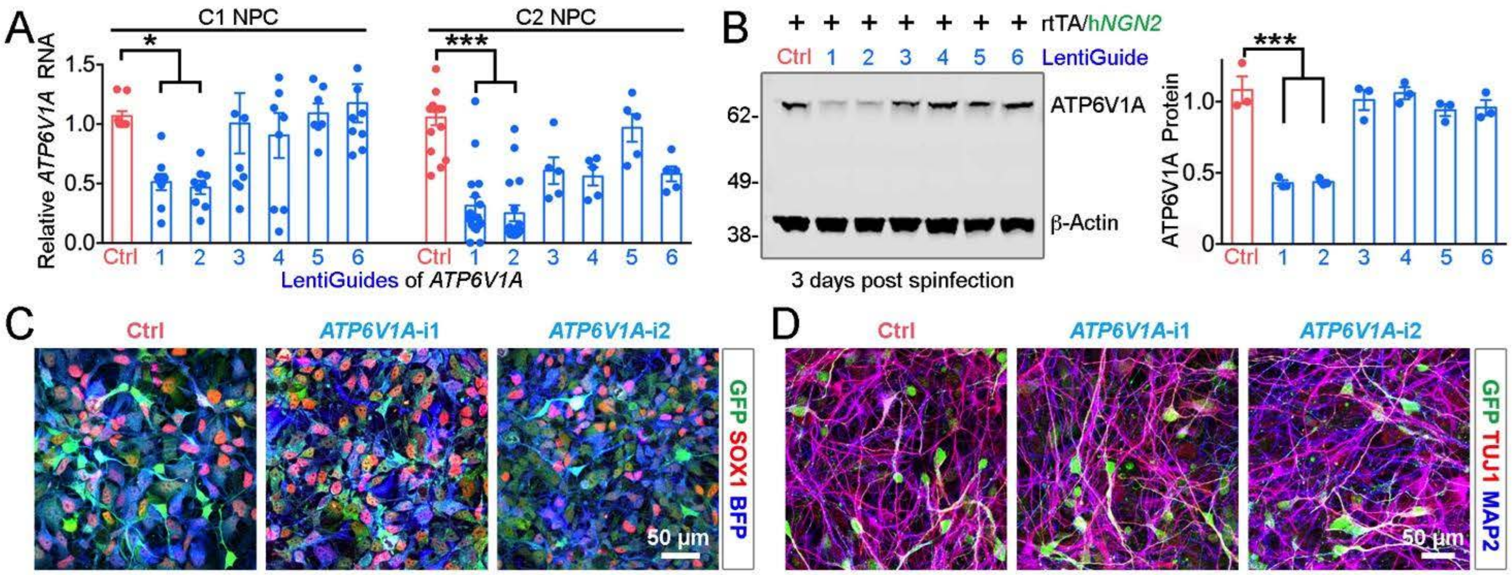


**Fig. S16. Evaluation of six gRNAs for repression of *ATP6V1A* in hiPSC-derived NPCs and *NGN2*-neurons. A-B**, Normalized relative RNA and protein levels (compared to an empty backbone control) following transduction of dCas9-KRAB NPCs with lentivirus-expressing gRNA targeting *ATP6V1A*. Red: control. Blue: CRISPRi. ANOVA; *p < 0.05 and ***p < 0.001; Error bars represent SE. **C-D**, *ATP6V1A* CRISPRi does not affect the neuronal differentiation of NPCs to *NGN2*-neurons. SOX1 is a neural stem cell marker (red); TUJ1 (red) and MAP2 (blue) are pan-neuronal markers. Bars, 50 µm.

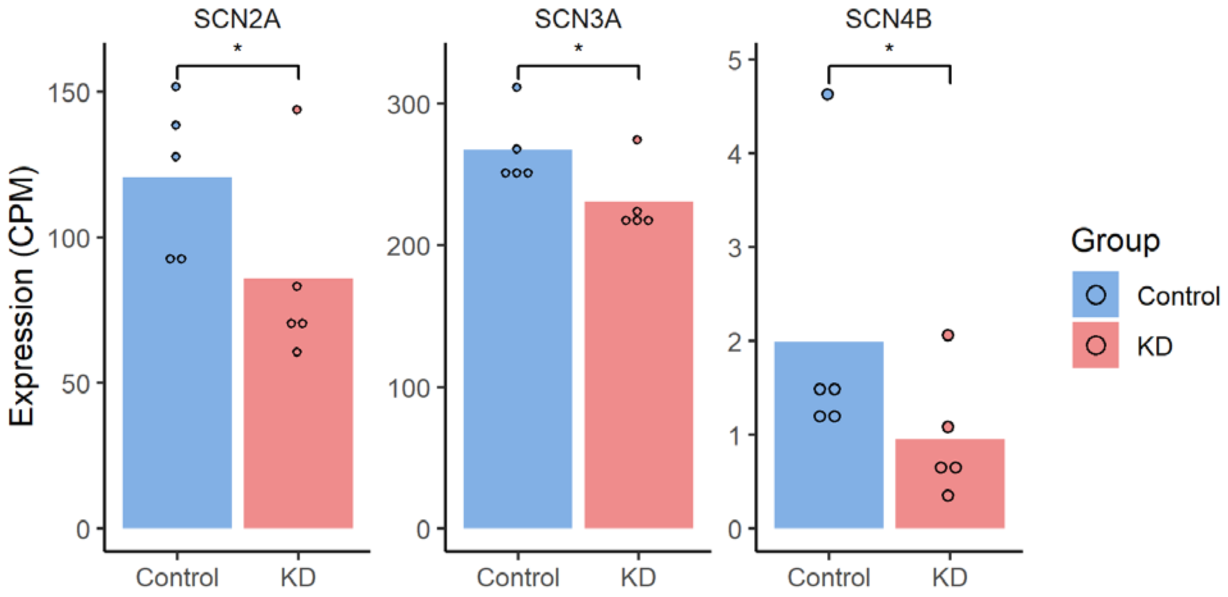


**Fig. S17. RNA-seq revealed reduced mRNA expression of voltage gated sodium channel subunits *SCN3A*, *SCN2A*, and *SCN4B* in *ATP6V1A* KD *NGN2*-neurons.** *$p < 0.05$ by Student's t-test.

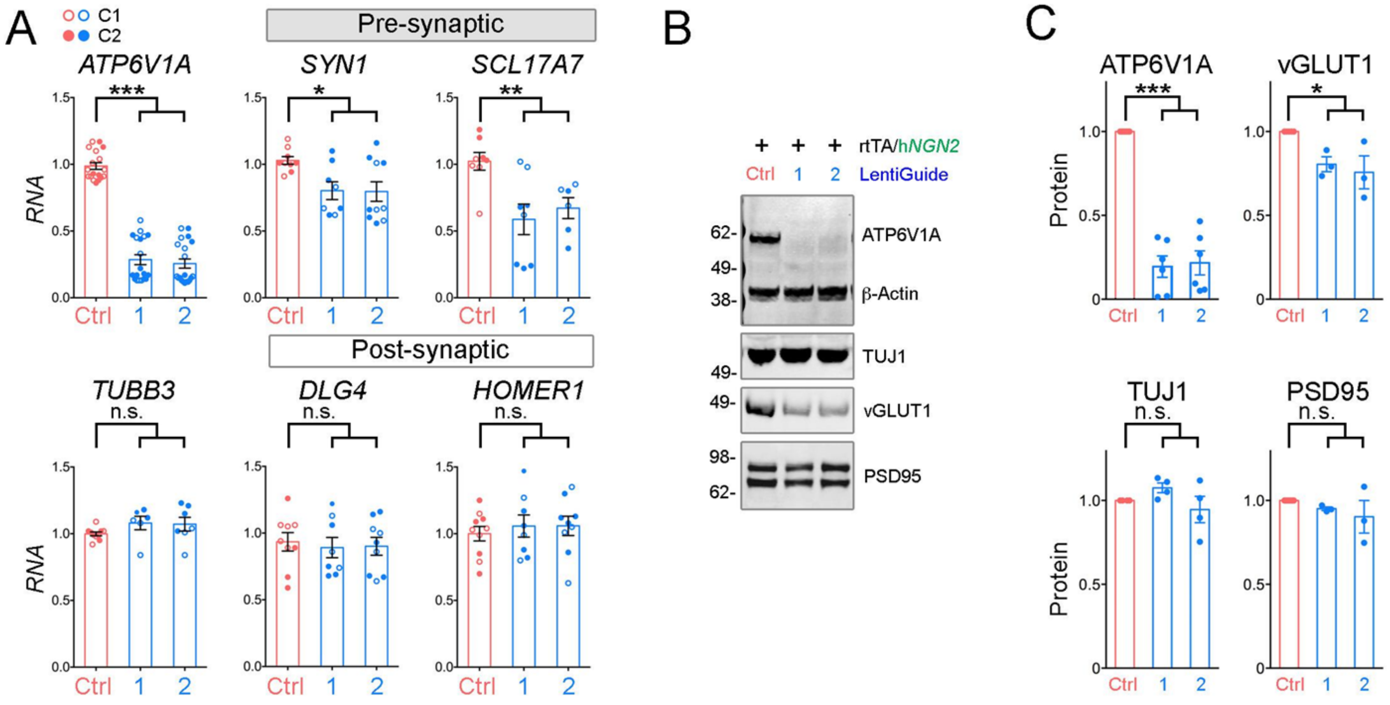


**Fig. S18. Expression of synaptic components in hiPSC-derived *NGN2*-neurons. A**, qRT-PCR analysis of expression of *ATP6V1A*, *TUBB3*, *SYN1*, *SCL17A7*, *DLG4* and *HOMER1* genes. n = 6-20 replicates. **B-C**, Western blot analysis and quantification of ATP6V1A, β-Actin, TUJ1, VGLUT1 and HOMER1 protein levels. Data represent the mean, n = 3 independent experiments. ANOVA; $*p < 0.05$; $**p < 0.01$; $***p < 0.001$; n.s., no significance; Error bars represent SE.

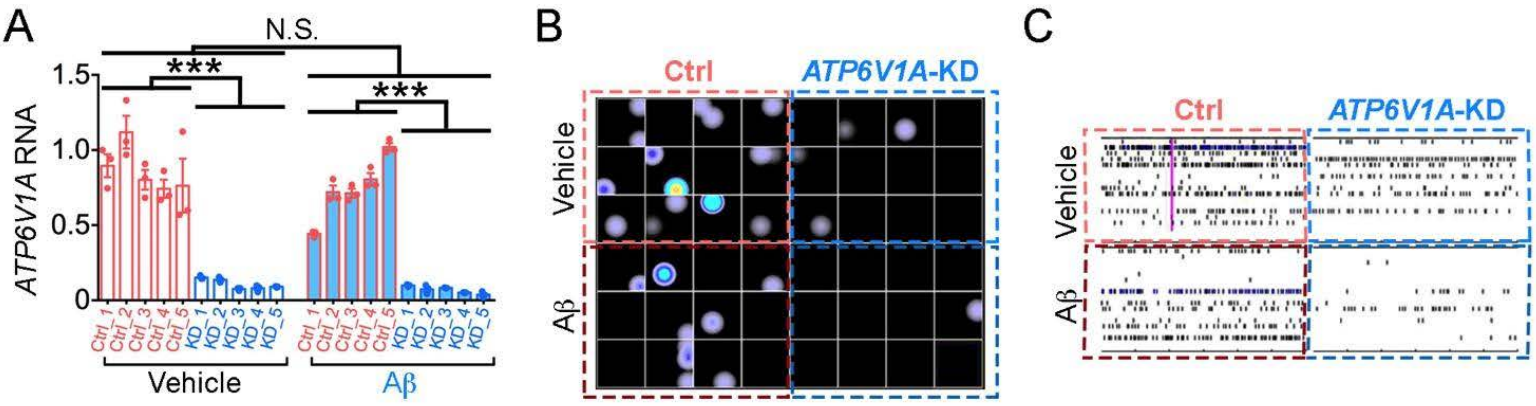


**Fig. S19. MEA arrays in *ATP6V1A*-deficient *NGN2*-neurons with or without exposure to Aβ. A**, *ATP6V1A* RNA levels across different samples. n = 3 replicates. ANOVA; ***p < 0.001; N.S., no significance; Error bars represent SE. **B**, Representative heat map recording of a CytoView MEA 48 plate. **C**, Representative raster plots of the spike events over 10 minutes of day 21 (D21) *NGN2*-neurons.

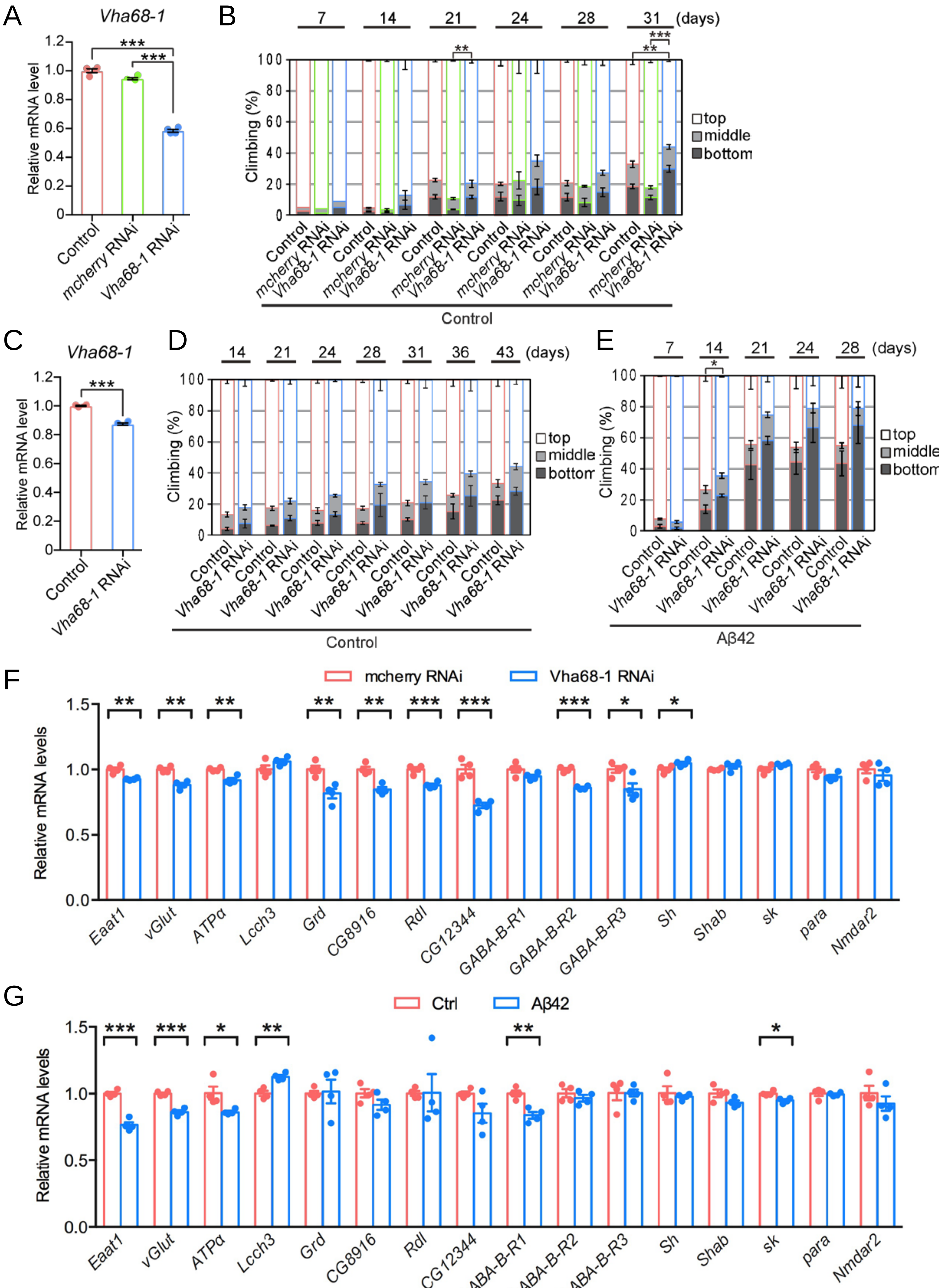
A
Vha68-1
Relative mRNA level
Control
mcherry RNAi
Vha68-1 RNAi
B
7
14
21
24
28
31
(days)
Climbing (%)
top
middle
bottom
Control
C
Vha68-1
D
14
21
24
28
31
36
43
(days)
E
7
14
21
24
28
(days)
Aβ42
F
mcherry RNAi
Vha68-1 RNAi
Relative mRNA levels
Eaat1
vGlut
ATPα
Lcch3
Grd
CG8916
Rdl
CG12344
GABA-B-R1
GABA-B-R2
GABA-B-R3
Sh
Shab
sk
para
Nmdar2
G
Ctrl
Aβ42

Fig. S20

**Fig. S20. Neuronal knockdown of *Vha68-1* exacerbates behavioral deficits caused by overexpression of Aβ42 peptide**. **A**, mRNA expression levels of *Vha68-1* in heads of flies expressing RNAi targeting *Vha68-1* (line #50726) were analyzed by qRT-PCR. $n = 4$, ***$p < 0.001$ by Student's t-test. **B**, Neuronal knockdown of *Vha68-1* (line #50726) by itself caused modest decline in climbing ability in aged flies. Average percentages of flies that climbed to the top (white), climbed to the middle (light gray), or stayed at the bottom (dark gray) of the vials. Percentages of flies that stayed at the bottom were subjected to statistical analyses. $n = 5$ independent experiments except for day 7 when $n = 2$, **$p < 0.01$ and ***$p < 0.001$ by Student's t-test. **C**, mRNA levels of *Vha68-1* in heads of flies expressing RNAi targeting *Vha68-1* (line #42888) were analyzed by qRT-PCR. $n = 4$, ***$p < 0.001$ by Student's t-test. **D**-**E**, Neuronal knockdown of *Vha68-1* (line #42888) did not alter climbing ability in control flies (**D**) but slightly enhanced locomotor deficits in Aβ42 flies (**E**). Percentages of flies that stayed at the bottom were subjected to statistical analyses. $n = 5$ independent experiments, **$p < 0.01$ and ***$p < 0.001$ by Student's t-test. The genotypes of the flies were: **A** and **B**, (Control): *elav*-GAL4/Y; +/CyO, (*mcherry* RNAi): *elav*-GAL4/Y; +/CyO; UAS-*mcherry* RNAi/+, (*Vha68-1* RNAi): *elav*-GAL4/Y; +/CyO; UAS-*Vha68-1* RNAi (line #50726)/+;, **C** and **E**, (Control): *elav*-GAL4/Y, (*Vha68-1* RNAi): *elav*-GAL4/Y; UAS-*Vha68-1* RNAi (line #42888)/+;, **D**, (Aβ42 and Control): *elav*-GAL4/Y; UAS-Aβ42/+; and (Aβ42 and *Vha68-1* RNAi): elav-GAL4/Y; UAS-Aβ42/UAS-*Vha68-1* RNAi (line #42888). **F** and **G**, qRT-PCR analysis indicated significantly reduced mRNA expression of 9 genes by neuronal KD of *ATP6V1A*/*Vha68-1* in fly brains (**F**) and 5 genes in Aβ42 fly brains (**G**). Ctrl, control. $n = 4$.

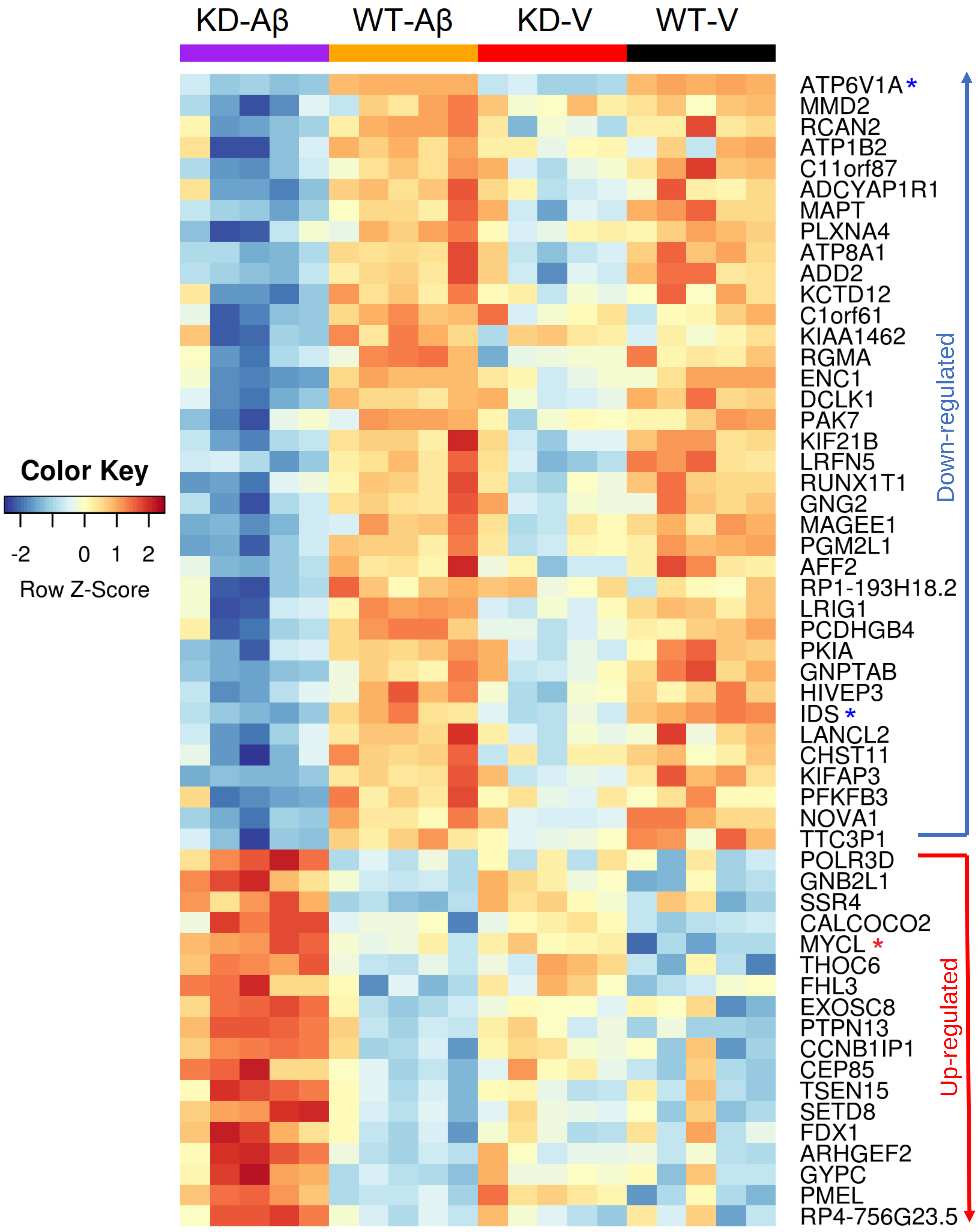


**Fig. S21. Heat-map showing genes significantly differentially expressed between *ATP6V1A* KD and WT in *NGN2*-neurons with or without Aβ treatment.** The expression values have been converted to z-score. Blue and red asterisks (*) denote, respectively, the genes consistently down- or up-regulated irrespective of Aβ treatment (i.e. KD-V vs WT-V and KD-Aβ vs WT-Aβ). Genes without any symbol annotation are detected only in Aβ treated cells (i.e. KD-Aβ vs WT-Aβ).

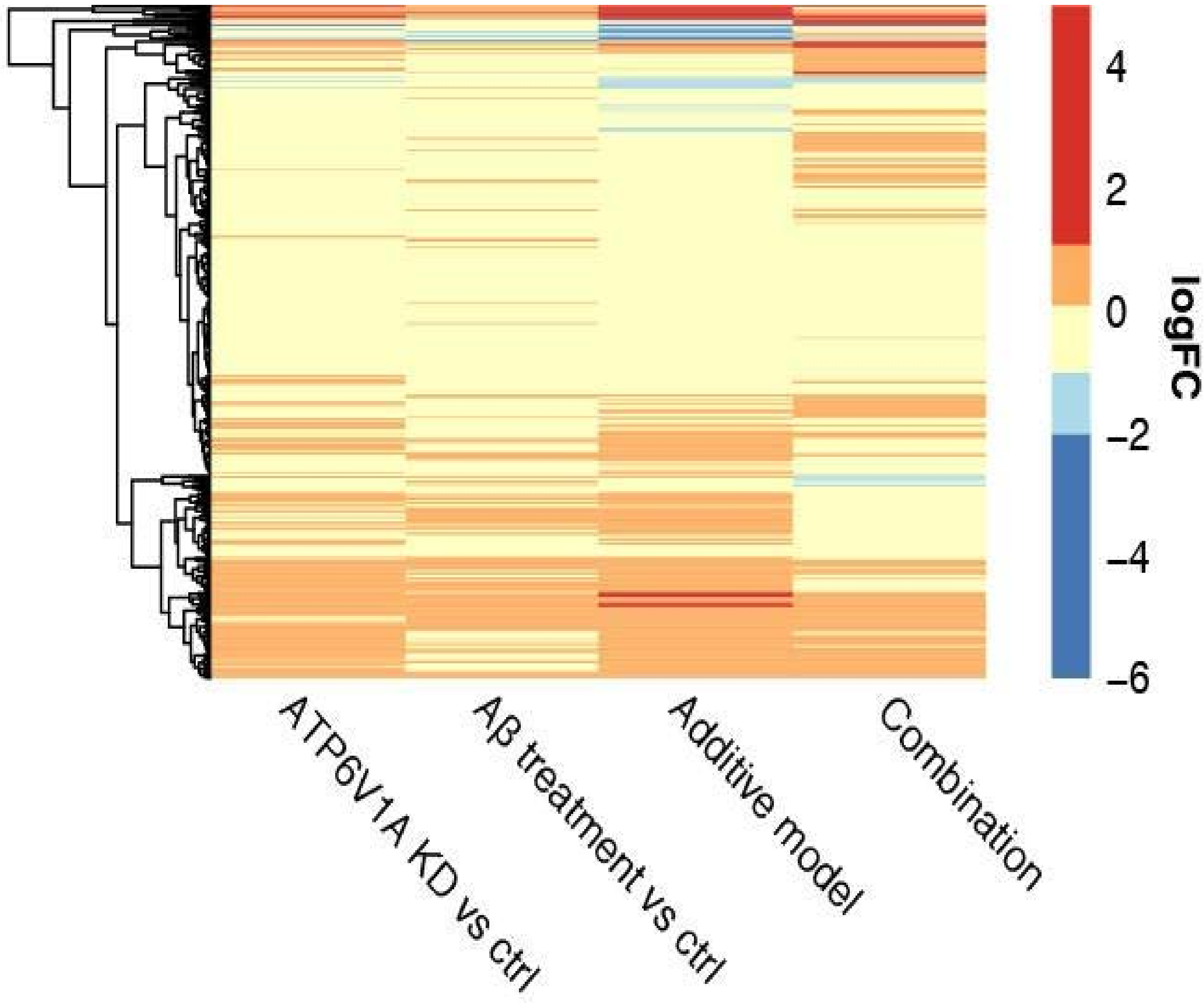


**Fig. S22. Hierarchical clustering of differential expression log2 fold changes (logFC) for all contrasts in *NGN2*-neurons treated with *ATP6V1A* KD and/or Aβ.** Genes were pre-aggregated using k-means (k=500). Color gradient represents logFC.
